# Unbiased lexicometry analyses illuminate plague dynamics during the second pandemic

**DOI:** 10.1101/2021.05.06.442897

**Authors:** Rémi Barbieri, Riccardo Nodari, Michel Signoli, Sara Epis, Didier Raoult, Michel Drancourt

## Abstract

Knowledge of the second plague pandemic that swept over Europe during the 14th-19th centuries, mainly relies on the exegesis of contemporary texts, prone to interpretive bias. Leveraging bioinformatic tools routinely used in biology, we here developed a quantitative lexicography of 32 texts describing two major plague outbreaks using contemporary plague-unrelated texts as negative controls. Nested, network and category analyses of a 207-word pan-lexicome over-represented in plague-related texts, indicated that “buboes” and “carbuncles” words were significantly associated with plague, signaling ectoparasite-borne plague. Moreover, plague-related words were associated with the words “merchandise”, “movable”, “tatters”, “bed” and “clothes”, while no association was found with rats and fleas. These results support the hypothesis that during the second plague pandemic, human ectoparasites were the major drivers of plague. Analyzing ancient texts using here reported method would certify plague-related historical records and indicate particularities of each plague outbreak, including sources of the causative *Yersinia pestis*.

## Introduction

Plague, a deadly zoonosis caused by the bacterium *Yersinia pestis* [1], has been incontrovertibly identified via paleomicrobiological methods in numerous historically described burial sites in Europe, ending decades-long controversies regarding the etiology of the so-called “Black Death” episode (1346-1353) and related episodes forming the 1346-18^th^ century second plague pandemic [2–7]. Ancient texts related to these plague episodes reported massive mortality [8–11], with an estimated ∼ 30 million deaths attributed solely to the “Black Death” episode (4, 11). Such figures have never again been observed during the plague outbreaks forming the 1894-current third plague pandemic [12, 13], leaving unknown the forces that shaped plague dynamics during the second plague pandemic.

Indeed, the sources and routes of the spread of *Y. pestis* in medieval European populations remain speculative, deriving as they do from the mathematical modeling of ancient plague episodes [13], which obviously cannot be verified, and from cumulative observations made during the third pandemic, which may not be applicable to the second pandemic [14, 15]. As an example, transmission of *Y. pestis* by rats and their flea ectoparasites or by the respiratory route, well described during the limited 20^th^ century plague outbreaks [13,16,17], may not have been sufficient to fuel the massive medieval mortality, pointing to the complementary role of human lice (*Pediculus humanus humanus*) and anthropophilic fleas (*Pulex irritans*) [13,18,19]. Accordingly, the relative contributions of rats, other mammals and animal and human ectoparasites to the dynamics of the second plague pandemic are poorly documented. Historical texts describing ancient plague outbreaks have been studied for more than one century [20–22], yet measuring the relative contributions of each factor during the second plague pandemic remains challenging in the absence of any quantitative analysis involving comparison with negative control texts, perpetuating the controversies regarding the dynamics of historical plague outbreaks [13,18,23,24].

To resolve some of these controversies, we designed a purposely naïve unbiased method for the automatic analysis of ancient plague texts derived from a method routinely used in biology to measure the relative expression of genes in a set of biological samples including negative control samples [25]. This model uses ancient plague-related text sets and negative control texts to measure the differential expression of plague-related words, creating a core-lexicome (core signature) of ancient plague and an accessory-lexicome specific to each ancient plague outbreak and opening an avenue for studying the pan- lexicome of plague. The method described here was developed based on positive control texts describing the paleomicrobiologically confirmed Great Plague that affected the city of Marseille and Provence at large in 1720-1722 [2,6,26,27] and was subsequently applied to texts describing the historical plague of 1629-1631, which ravaged Northern Italy, an episode still in need of paleomicrobiological confirmation [27, 28].

We cataloged words significantly associated with plague in these records and derived original knowledge regarding the sources and transmission of *Y. pestis* during two outbreaks attributed to the second pandemic in Europe.

## Results

### Building the Marseille word database

A word repertoire drew from 16 historical French texts describing the microbiologically documented 1720-1722 Great Plague of Marseille, France [2,6,26], and from 11 contemporary plague-unrelated French texts dating from the same historical period (Data file S1 and S2), after verification that the maximum Jaccard index between two texts was 0.08 using the “Jaccard” package in Rstudio (fig. S1A). This ensured that the 27 texts were original enough (without significant plagiarism) to be used as independent samples. This repertoire comprised a total of 2,049,566 words directly imported (without any modification) from Google Books into Rstudio, including 473,681 words (23.1%) poorly digitized due to damage resulting from preservation conditions and that therefore could not be included in further analysis. Then, 1,575,885 words (76.9%) representative of 10,315 unique words were cleaned by removing noncharacter letters and spaces, replacing ancient characters with homologous characters in the contemporary alphabet (i.e., the “long S” was replaced with “s”), filtering (with a French glossary of 336,631 words, conjugated verbs and proper nouns) and deleting words with fewer than five occurrences in all 27 texts (Complete script present as Supplementary Text 1). Altogether, the final 1,575,885-word database comprised 904,005 words from the 16 plague-related texts and 671,880 words from the 11 negative control texts.

### Analyzing the Marseille word database

This word database was used to compare the relative expression of words in the 16 plague-related texts and the 11 negative control texts (Fig. 2A). This comparison was performed by leveraging informatics methods and employing cut-off values commonly used in the field of transcriptomics (the analysis of RNA sequences, i.e., significant letter series using the universal biological code, in biological samples) (Fig. 1). Accordingly, using the DESeq2 package originally developed to identify dysregulated genes in transcriptome analyses, we generated a list of 70 words (log2foldchange >0 and *P* adjust<=0.05) (0.0007%), being overrepresented in plague-related texts compared to control texts as presented in the WordCloud (Fig. 2; Table S1; Data file S3). Two words are represented twice such as “Contagious” and “Infected”. Our results confirmed the significant report of the two plague signs, namely, “bubo” (*P* adjust= 5e-07) and “carbuncle” (*P* adjust= 0.003). Notably, these two signs have been continuously described, by Procopius in Antiquity [29], Guy de Chauliac in medieval times [30] and Yersin and Simond at the beginning of the third, current plague pandemic [31, 32]. The significant detection of these two words, constituting internal positive controls, validated our method and allowed us to consider enriched words as significantly associated with plague. Notably, the words “rats” and “fleas” appeared only 11 and 5 times in plague-related texts versus 5 and 3 times in negative control texts, respectively; the differences were not significant. Further careful examination revealed that the word “rat” was misleading in 5 cases, resulting from the abbreviation of the word “magistrate”; ultimately, there were 6 true occurrences in one text reporting dead rats in the streets and rats fleeing houses where plague was declared, but the author spoke of plague in general, and in any case, he does not mention rats during the Marseille plague [33]. Likewise, the word “flea” was mainly used to describe spots on the dura, observed during autopsy, that looked “like flea bites” and once to describe spots on the belly of a plague-patient as being “like flea bites”. In only one case, “flea” unambiguously referred to the ectoparasite, but in an out-of-plague context.

**Fig. 1.**
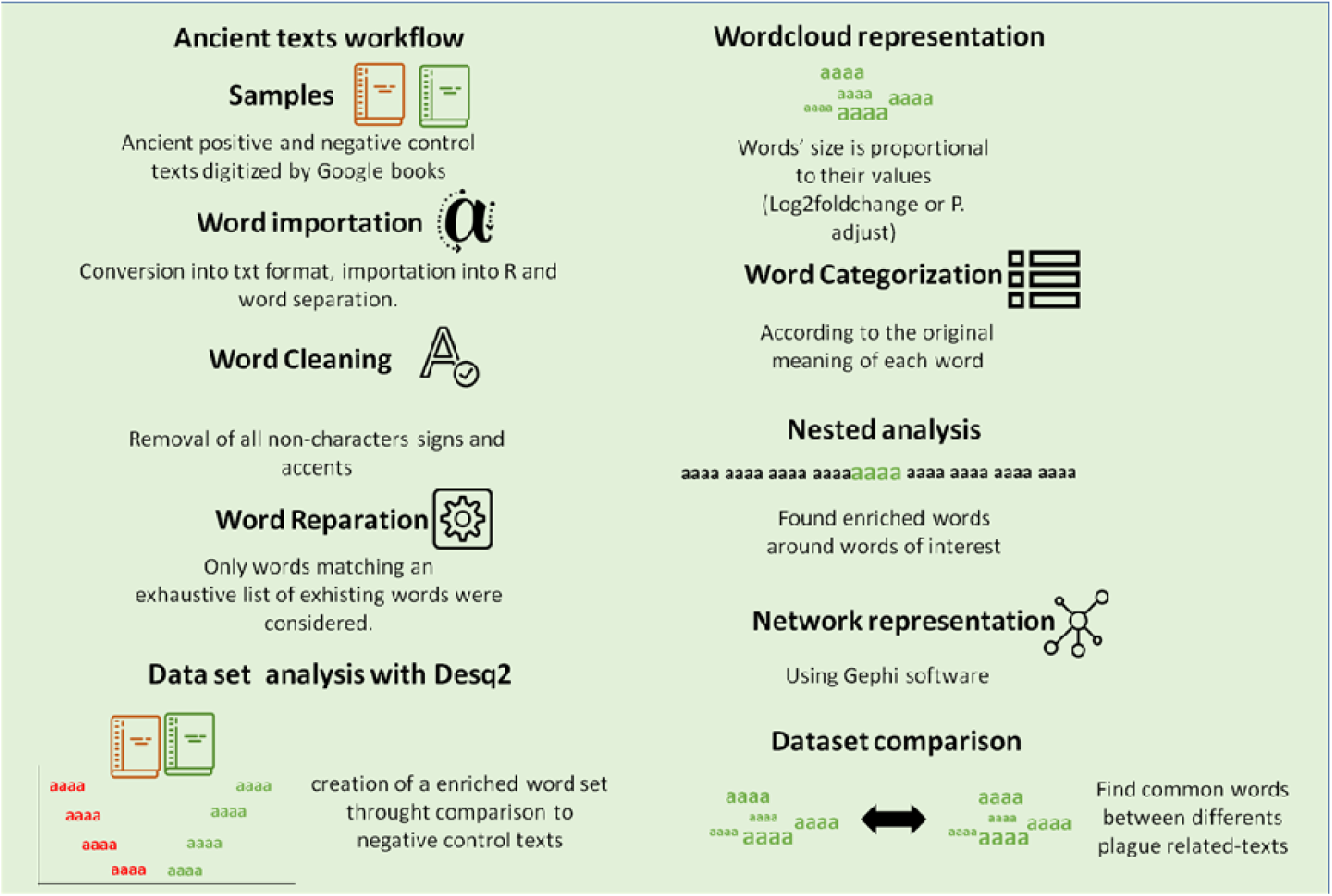
Workflow summarizing the main steps of the method developed in this study.

**Fig. 2.**
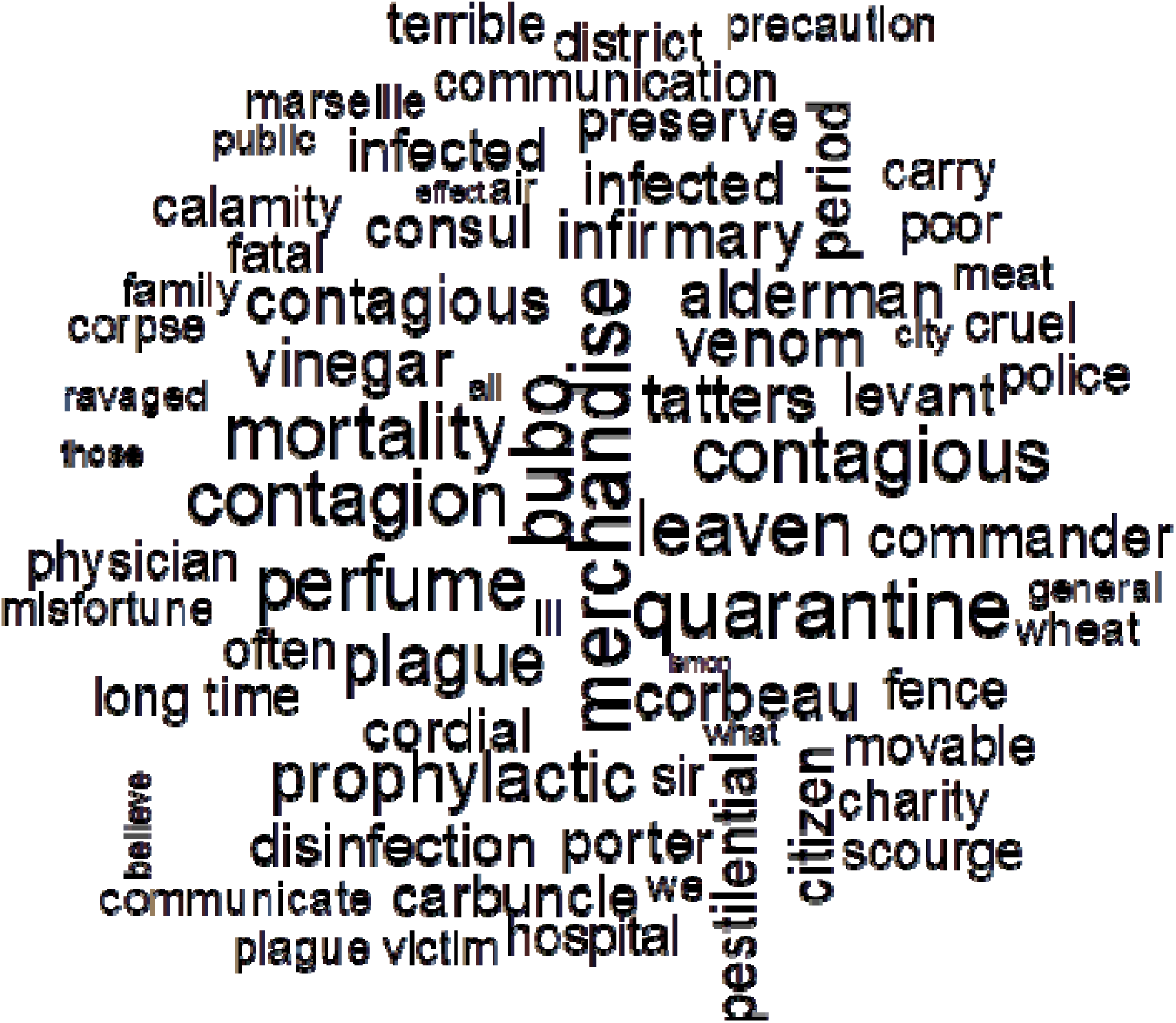
Wordcloud displaying the words that were overrepresented in the Marseille plague- related texts (Padj <=0.05 and log2foldchange >0). The size of the words represents their adjusted P enrichment in plague-related texts compared to negative control texts.

### Categorizing the Marseille word database

By analogy with bioinformatics, where clusters of orthologous genes (COGs) are defined as functional categories [34], the 70 plague words were classified into 19 categories of words, each category comprising words related to the same function in the plague-related texts (Data file S4). The category “Other” was the most abundant, comprising 15/70 (21%) unclassified words, followed by the categories “Consequence” (10/70 words, 14%), “Public Response” (8/70 words, 11%) and “Nature of plague” (7/70 words, 10%). Some words, such as “police”, “physician”, “hospital” and “infirmary” were classified into two different categories: “Prevention” and “Public Response”. Indeed, it appeared that these professions or places already existing before the plague surge were strengthened due to the catastrophic situation because of the construction of new infirmaries and hospitals and the increase in the number of doctors and police personnel. Faced with the assignation of some words to several categories, we specifically consolidated the category “Sources” by analyzing every sentence in which each word of this category was used to determine the number of its occurrence as a *bona fide* plague source. This analysis indicated that the word “movable” was used in 194/207 (94%) occurrences as a plague source; “tatters” was used in 196/225 (87%) occurrences as a plague source; “clothes” was used in 128/153 (83%) occurrences as a plague source; and “merchandise” was used in 561/810 (69%) occurrences as a plague source (Table 1).

**Table 1.**
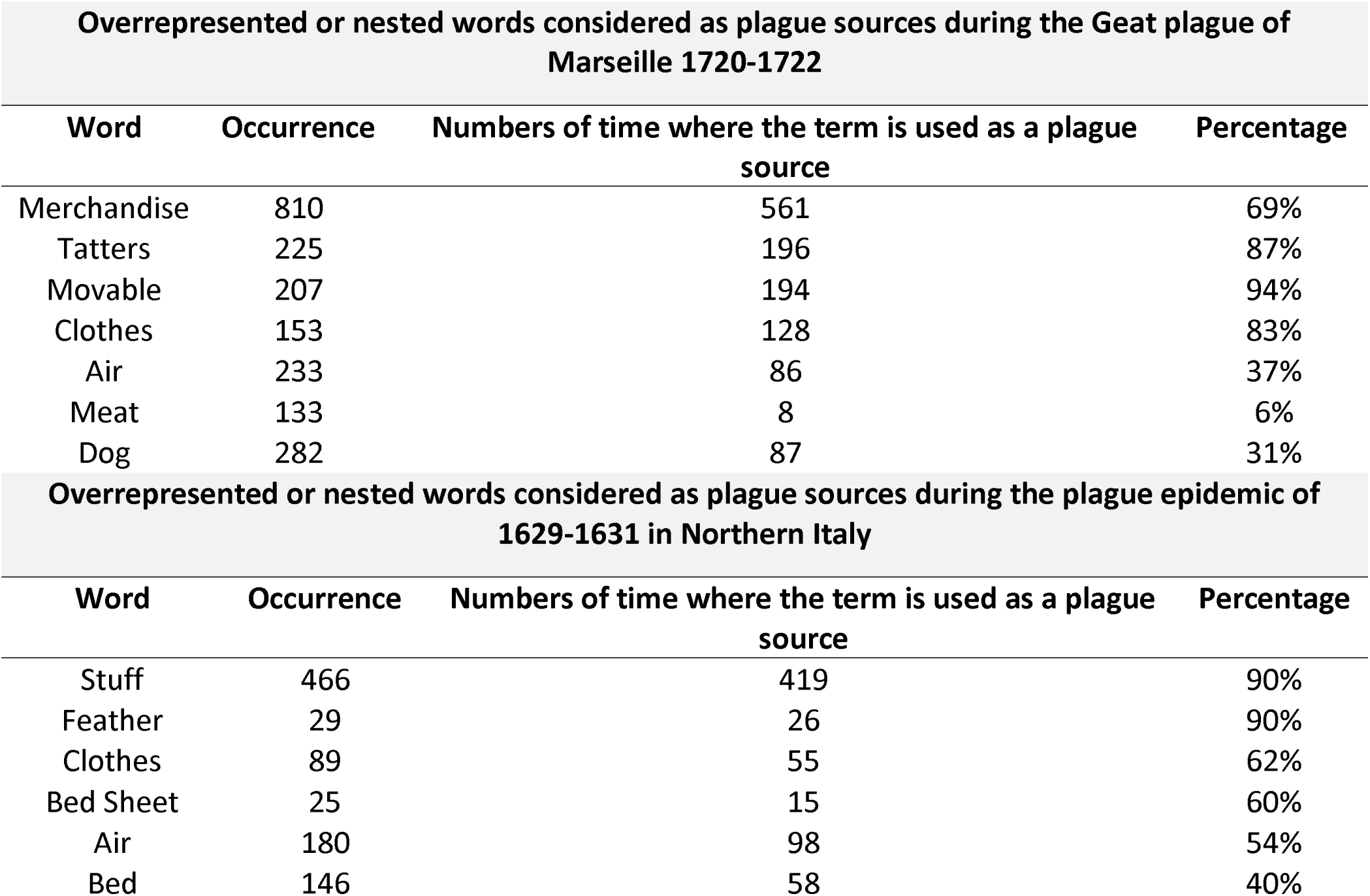
Overrepresented or nested words considered as plague sources in the analyzed records. Table representing the number of occurrences of each suspected plague source in plague-related texts from Marseille and Northern Italy featuring the percentage each word is used by ancient authors as a source of plague. Words for the “bona fide” analysis were selected depending on the results of the nested analysis, for the association with plague-related words.

| Overrepresented or nested words considered as plague sources during the Geat plague of Marseille 1720-1722 |  |  |  |
| --- | --- | --- | --- |
| Word | Occurrence | Numbers of time where the term is used as a plague source | Percentage |
| Merchandise | 810 | 561 | 69% |
| Tatters | 225 | 196 | 87% |
| Movable | 207 | 194 | 94% |
| Clothes | 153 | 128 | 83% |
| Air | 233 | 86 | 37% |
| Meat | 133 | 8 | 6% |
| Dog | 282 | 87 | 31% |
| Overrepresented or nested words considered as plague sources during the plague epidemic of 1629-1631 in Northern Italy |  |  |  |
| Word | Occurrence | Numbers of time where the term is used as a plague source | Percentage |
| Stuff | 466 | 419 | 90% |
| Feather | 29 | 26 | 90% |
| Clothes | 89 | 55 | 62% |
| Bed Sheet | 25 | 15 | 60% |
| Air | 180 | 98 | 54% |
| Bed | 146 | 58 | 40% |

### Marseille word database: nested and network analyses

Nested analysis was used to contextualize the 70 words overrepresented in the plague-related texts by extracting the 25 words downward and the 25 words upward of each of these overrepresented words and then testing the significance of each; WordClouds for each overrepresented word was generated using a *P* adj < 0.001 (Data file S5; fig. S3). As an illustration of this nested analysis representation, the word “merchandise” (Fig. 3) in the category “Source” was significantly associated with “Levant”, “smugglers”, “June”, “fur”, “ship”, “clear”, “unseating”, “exposed”, “purged”, “tatters”, “transport”, “porter” and “Jarre” (the name of the quarantine island in the Marseille gulf) (*P* adj < 0.001) (Fig. 3). Likewise, in the symptoms category, the word “carbuncles’’ (the gangrenous lesions consecutive to the blisters corresponding to a microbe’s gateway through the skin) [32] had 625 occurrences and was significantly associated with “groin”, “parotid”, “arms’’, “leg” and “thigh”, all words indicative of the topography of the lesions (*P*adj < 0.001). The word “bubo” (the lymph node proximal to the inoculation site (blister or carbuncle)) [32, 35] had 948 occurrences and was significantly associated with “groin”, “armpit”, “arm”, “throat”, “parotid”, “ear”, “maxillary”, “jugular” and “thigh” (*P*adj<0.001) (Fig. 3), all words designating localizations unsurprisingly similar to those associated with “carbuncles”, with 95% similarity. Concerning plague sources, the words “tatters”, “movable” (*P* adj<0.001), “cloth” and “air” (*P*adj<0.005) were found to be linked with “infected” and multiple other plague-related words (Fig. 4). The word “tatters” was associated to infection (e.g. “kill”, “corrupt”, “communicated”, “filth”, “contaminates”, “miasms”, “infect”, “infection” *P*adj<0.001), disinfection-related words (e.g. “purge”, “burn”, “aromatic”, “confiscation”, “punished” *P*adj<0.001), and also interesting words related to the dissemination of the plague and its origin (e.g. “smugglers”, “Turkish”, “Tripoli” *P*adj<0.001). The word “movable” was linked to words from the same lexical field as “tatters” (e.g. “tatters”, “cloth”, “clothes” *P*adj<<0.001). It was also associated with infection (e.g. “dangerous”, “infection” *P*adj<0.001) and disinfection-related words (e.g. “burn”, “purge”, “destroyed”, “disinfection”, “perfume”, “purify”, “confiscation” *P*adj<0.001). All these associations suggest that contemporaries considered tatters, clothes, and movable as plague sources that communicated plague across the city. Likewise, the word “merchandise” was linked to multiple disinfection related words (e.g. “purge”, “quarantine” *P*adj<0.001). The word “air” was dichotomously associated with contagion (e.g. “bad”, “corrupt” *P*adj<0.001, “infected” and “poisonous” *Padj*<0.005) and with purification (e.g. “purify”, “pure” *P*adj<0.001). In Fig. 4 it can be clearly seen that the word “infected” is linked with the words “tatter”, “cloth”, “movable”, “air” and “merchandise” (*P*adj<0.01). Other networks for potential sources of plague and the relation with other words are reported as fig. S5 and S6.

**Fig. 3.**
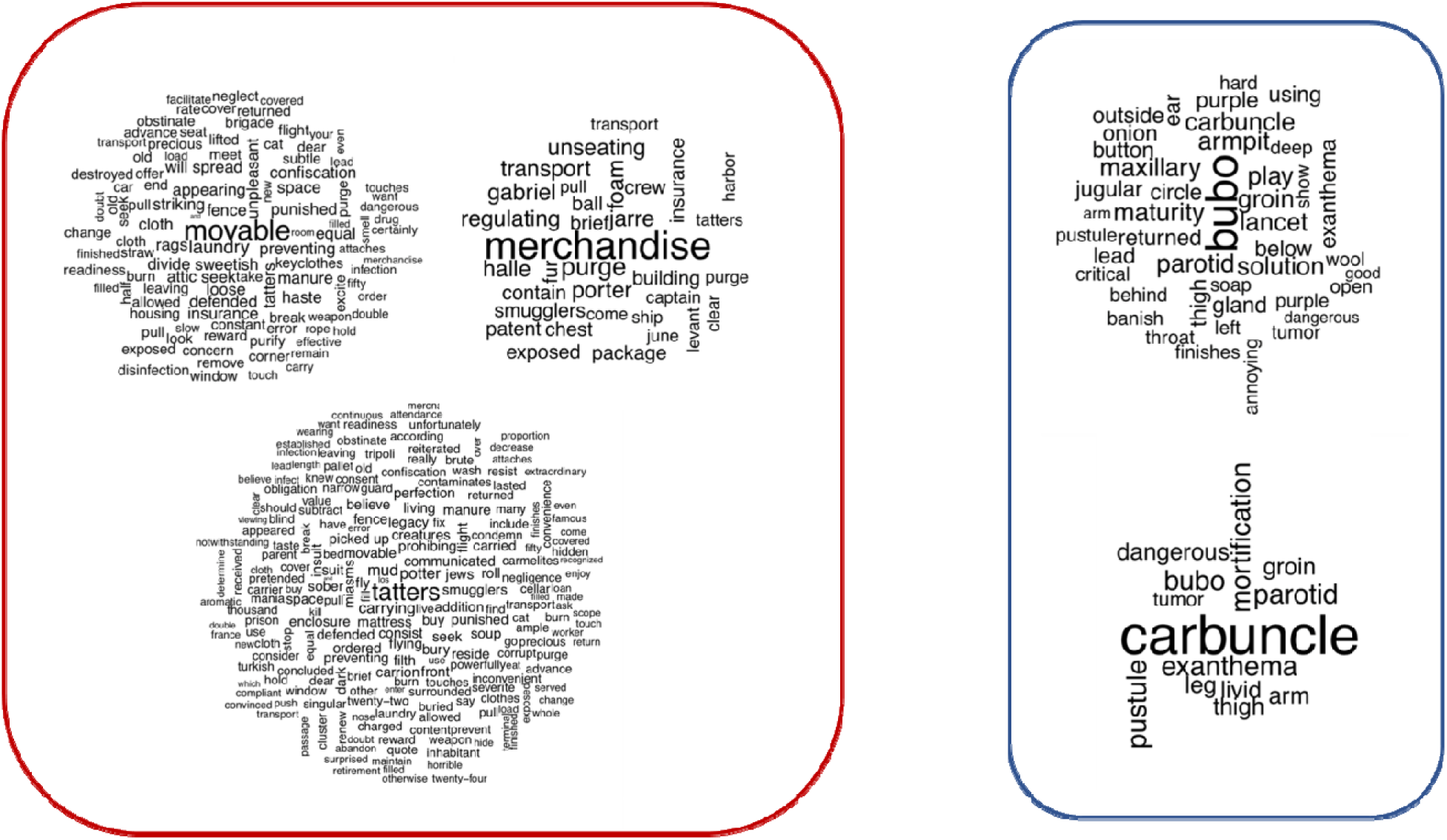
Wordcloud representations of the nested analysis displaying overrepresented words (Padj < 0.001) attached to plague sources (movable, tatters and merchandise) (red panel, left) and plague symptoms (buboes and carbuncles) (blue panel, right) during the 1720-1722 Great Plague of Marseille.

**Fig. 4.**
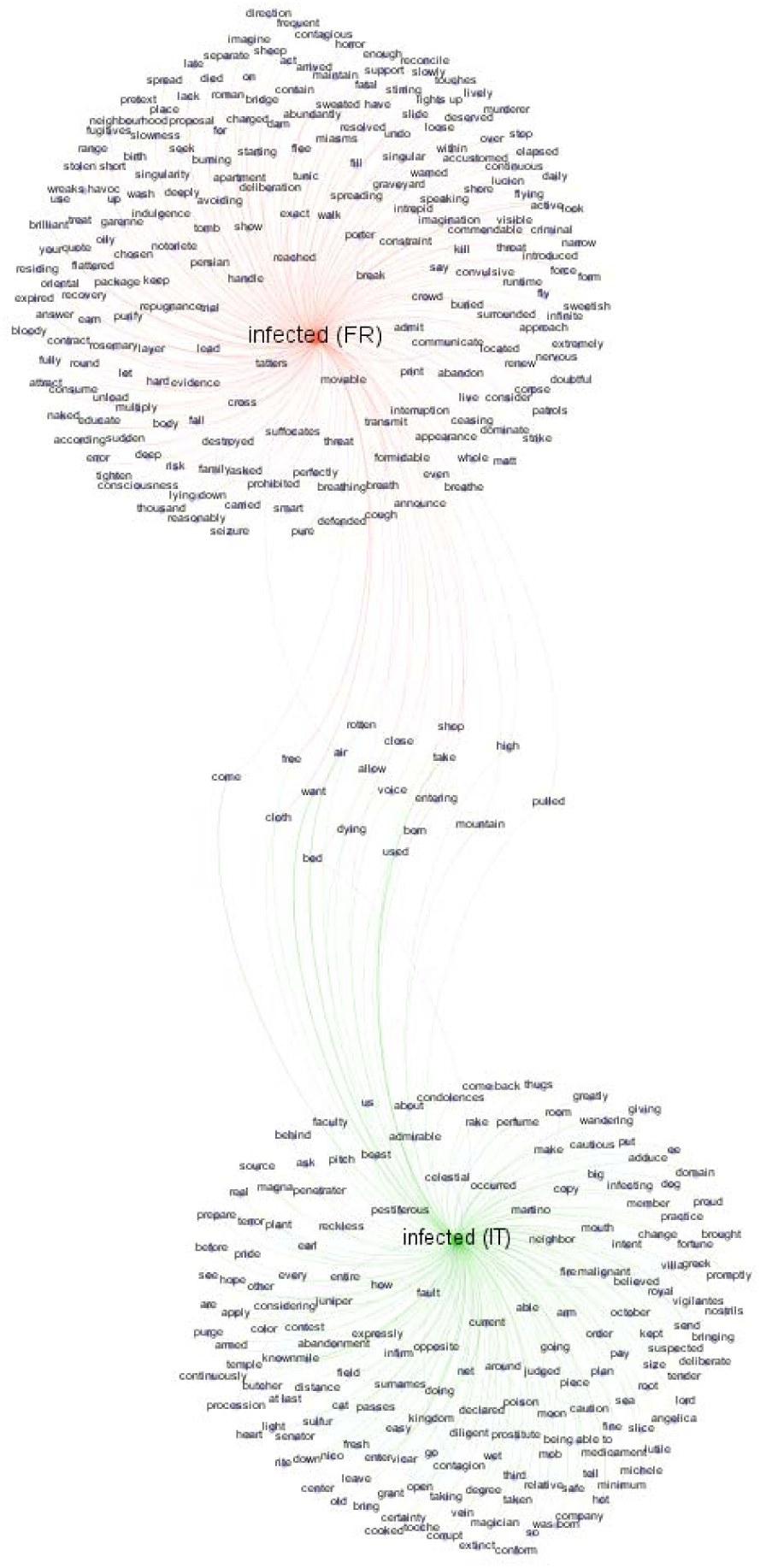
Network representation of the relations between words generated using Gephi software based on the adjusted P values from the nested analysis (P adj<0.01) of the word “infected” (in green for Marseille and in orange for Northern Italy). Word label dimensions are proportional to the number of links with other words.

### Documenting plague in Northern Italy

We then processed 16 historical texts describing an epidemic outbreak in 1629-1631 in Northern Italy (Data file S6), which was never microbiologically confirmed as plague [28]. Twenty Italian texts from the same historical period were used as negative control texts (Data file S7) (Jaccard index between two texts < 0.03) (fig. S1B). The repertoire of words included 1,574,910, of which a final word database of 1,063,483 words total, including 451,856 words from the plague-related texts and 611,627 words from the negative control texts. Using the DESeq2 package, we obtained a list of 147 overrepresented words (0.03%) (*P* adjust value <= 0.05 and log2foldchange>0) (Fig. 5) (Table S2; fig. S2B; Data file S8). Seven words are represented twice such as “contagion”, “could”, “did”, “infected”, “our” and “stuff” while “were” is represented three times. These 147 words were then classified into the same categories used for the Marseille text analysis (Data file S4), and the words in the “source” category were consolidated as explained previously (Table 1). In particular the Italian words “roba” and “robba”, which have been translated into the English word “stuff” and are used in a general way to indicate multiple things, such as movable, merchandise, food and especially clothes (Supplementary Text 2), were present more than 400 times in plague-related texts and in 90% of the cases the term was considered by the author as a plague source. Interestingly, the Italian word for “rat” was present only four times in the plague-related texts: it appeared twice in a figurative sense, once as a general premonition of plague (“If the underground animals flee their rooms, they come upon the earth, unable to live in the extreme putrefaction of their mother, which are worms, snakes, toads, rats, and being on the ground they die, then they are the cause of infection”) [36] and once in a statement that rats, together with dogs, cats, hens and doves, should be killed as a precaution to protect houses from the disease [36].

**Fig. 5.**
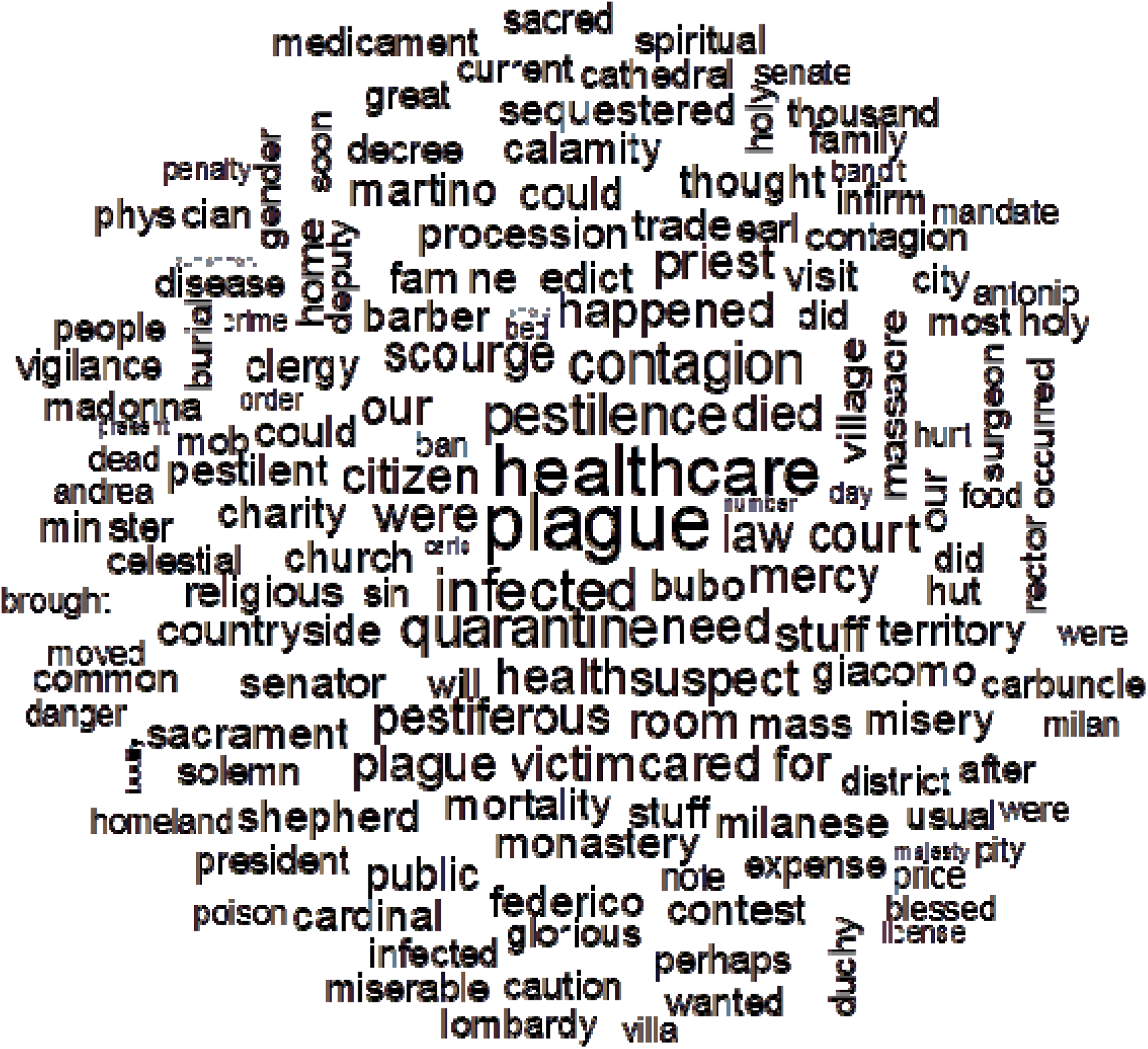
Wordcloud displaying words overrepresented in plague-related texts from Northern Italy displayed (P adjust value <=0.05 and log2foldchange >0). The size of the words represents their adjusted P enrichment in plague-related texts compared to controls.

Nested analysis on Italian overrepresented words indicated that words such as “clothes”, “beds”, “air” and “rooms” were directly linked to infection, disinfection and danger-related words, indicating clear attention to them as potential plague sources (Data file S9; fig. S4). In particular, the word “bed” was linked to infection (“infected”, “contagion”, “pestiferous” *P*adj<0.01), disinfection (“purge”, “perfume”, “air”, “boil” *P*adj<0.01), and “caution” (*P*adj<0.01). Similarly, the word “cloth” was found to be associated with plague-related words, such as “infected” (*P*adj<0.01), “caution” (*P*adj<0.001), and “poison” (*P*adj<0.01). The word “poison” was also associated with “clothes” (*P*adj<0.01). Moreover, the word “air” was linked to “infected” (*P*adj<0.001), “bed” (*P*adj<0.001), and “pestiferous” (*P*adj<0.01).

### Plague pan-, core- and accessory- lexicomes

To define the plague pan-, core- and accessory- lexicomes (the complete collection of words associated with plague, the plague words in common between the French and Italian lexicomes, and the plague words specific to each episode, respectively), all the French and Italian words were translated into English using the translation function in Microsoft® Word. Misleading translations of four words (three from French texts and 1 from Italian texts) resulted either from a truncated word (e.g., “conta” for “contamination”) or from an isolated letter (e.g., “o”), so that DESeq2 analysis incorporated a total of 68 unique words for Marseille and more than twice the number (139 unique words) for Northern Italy. This difference in the richness of the repertoire between the two datasets and the stronger association between words obtained from the nested analysis for French text respect to the Italian ones (Datasets S5 and S9), could be explained by the fact that the Italian texts in the Northern Italy dataset were largely written by nonmedical professionals such as religious figures, historians and economists (Dataset S6), while the French texts in the Marseille dataset were all written by doctors (Dataset S1). Additionally, the Italian texts described an epidemic covering multiple cities in multiple small countries in Northern Italy, while the French texts covered a much more geographically limited region. The plague pan-lexicome was composed of 10,743 words and included a core-lexicome of 2,645 words: 4,063 words are specific to Marseille’s plague and 4,035 to the Italian plague. Regarding overrepresented unique words, there were 207 in the pan-lexicome (Fig. 6); the core-lexicome had 17 unique words: 51 specifics to Marseille and 122 specifics to Northern Italy constituted the two accessory- lexicomes. Words significantly enriched in the Marseille plague texts had a significant probability of also being enriched in the texts related to the 1629-1631 Italian plague compared to control texts (Fisher’s exact test performed with two conditions, *P* val<0.001***; *P* val<0.00001****). The French and Italian datasets were then merged graphically to create networks of common and unique words. Words of interest present in both datasets were compared and placed together to form a network evidencing nested words that were unique or common to both datasets (*P*adj < 0.01). The network representing the significant symptoms of plague (bubo and carbuncle) showed overlap between the Marseille and Northern Italy datasets corresponding to the most common locations for buboes and carbuncles (“groin”, “armpit”, “throat”, “parotid”, “leg”, and “arm”) (Fig. 7). Interestingly, three words “air”, “bed” and “cloth” from the category “Sources” overlapped in the network for “infected” (Fig. 4). No other potential plague sources were observed in common between the Marseille and Northern Italy datasets.

**Fig. 6.**
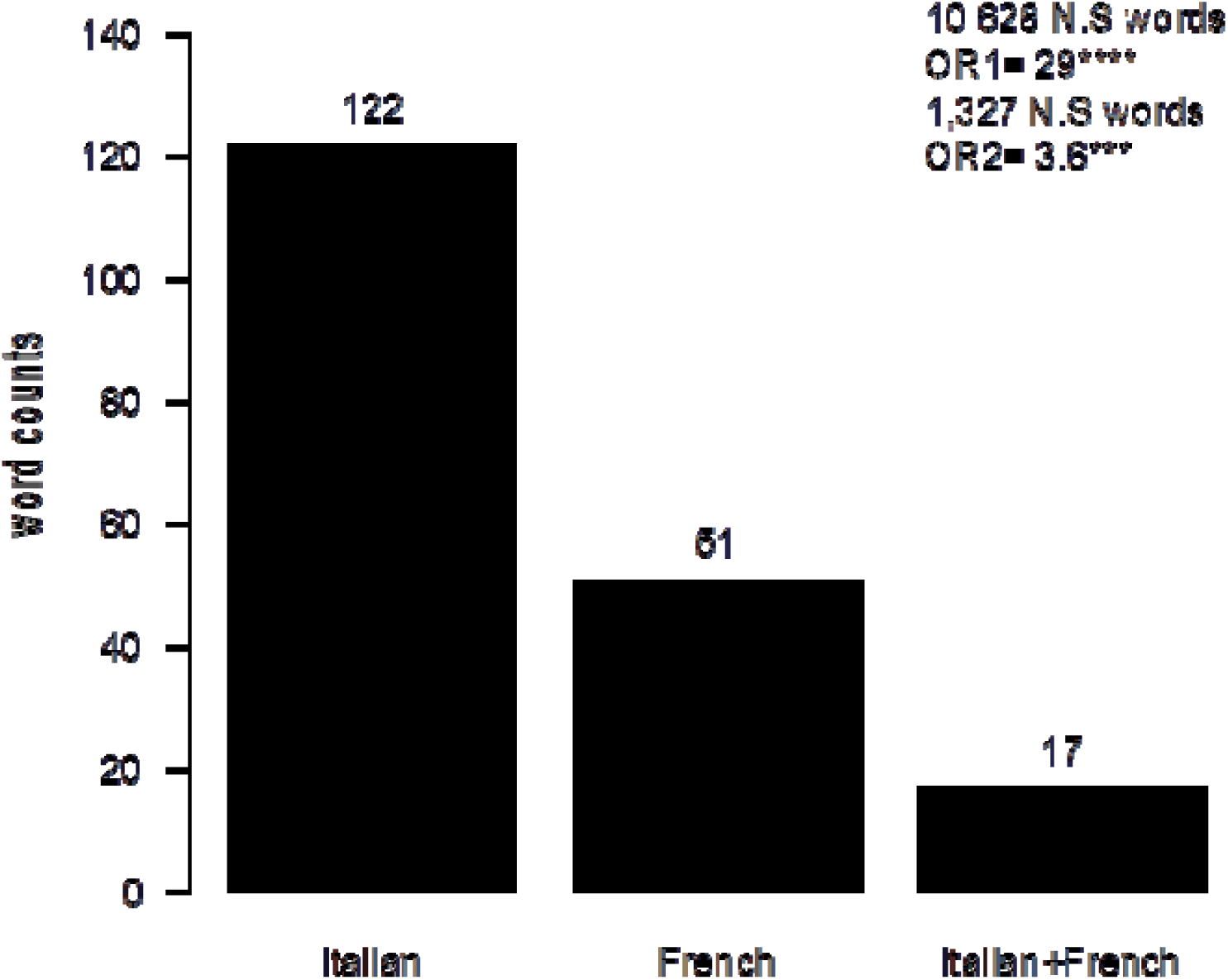
Barplot displaying the numbers of unique significantly over−represented words from Italian (IT) or French (FR) or shared between both. The overlap of significantly over−represented words between Italian and French texts was tested using fisher test on all non−significant words (OR1) or non−significant words with a baseMean > than the lower quartile of significant words (OR2). Pval<0.001***; Pval<0.00001****

**Fig. 7.**
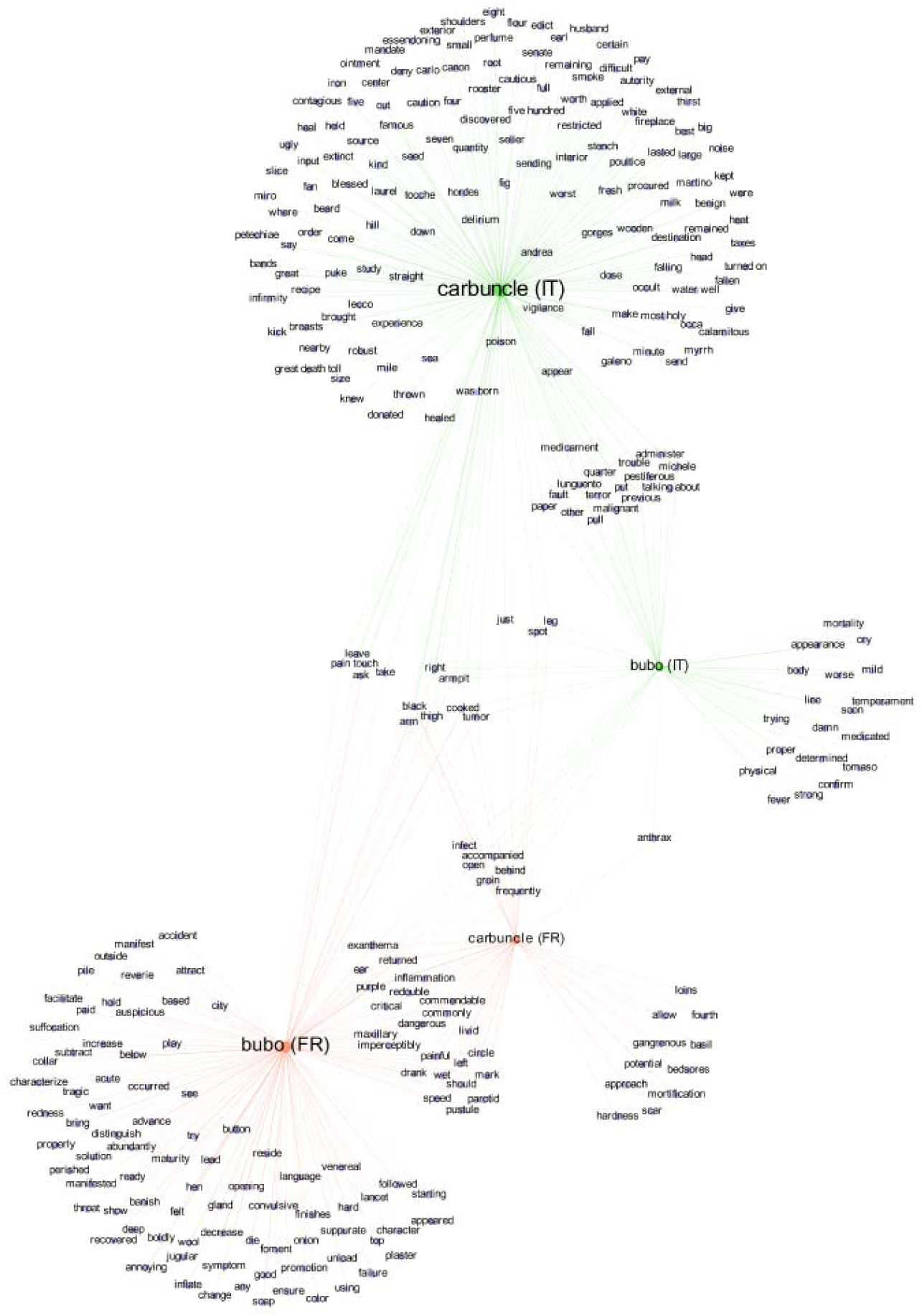
Word network representation of words related to two plague symptoms, “carbuncle” and “bubo”, from the Marseille plague (displayed in green) and the Northern Italy plague (displayed in orange). Network generated with Gephi software based on the adjusted P values from the nested analysis (P adj<0.01). Word label dimensions are proportional to the number of links with other words.

## Discussion

Updated lexicographic analyses of ancient texts describing deadly outbreaks proved efficient for identifying them as plague-describing texts and provided valuable information regarding plague dynamics during the second pandemic.

We developed and used a method derived from automatic bioinformatics to obtain quantitative information from ancient texts. Our data, coupled with paleomicrobiological analysis [2,6,26], confirm that these ancient texts speak of a plague caused by *Y. pestis,* as revealed by the significant presence of the words “buboes” and “carbuncles”, the two main symptoms of plague during the second pandemic. These results resolve a years-long debate as to the etiology of plague in historical sources. Indeed, plague was initially interpreted as a hemorrhagic fever [23,37,38] because of the absence of rats and their fleas and because some researchers thought that the term “Black Death” denoting the medieval plague partly reflected the symptoms of the disease. In reality, this term was anachronistic and remains largely metaphorical (reflecting the idea of disaster) [39]. Furthermore, through our analysis, we ascertained that contemporaries of the second plague pandemic did not consider animals as sources of plague. In no case did the historical texts significantly mention animals as a source of plague, disrupting the traditional picture of plague as a primary zoonosis during the second pandemic [18]. In particular, the words “rat” and “fleas”, repeatedly put forward as the source and vector of *Y. pestis* during the second plague pandemic [18, 40], did not reach significance and were not enriched in plague-related texts, relativizing the role played by rats and fleas in the transmission of the plague during the second pandemic. Whereas we cannot estimate the predictive negative value of words, such an absence cannot be taken as supporting the prominent role of rats and their ectoparasites during the epidemic episodes.

Instead, contemporaries traced plague outbreaks in Marseille and Northern Italy to imported stuff, a general word encompassing the significantly enriched words “movable”, “tatters”, “clothes”, and “merchandise”; specific references were made to fur imported via the maritime route in Marseille, with a possible unanticipated role of smuggling, and clothes brought by the French and Imperial Allemans soldiers into Northern Italy during the war of Mantua [41–43]. All such clothing was in direct contact with plague-stricken people, and contemporaries insisted on the dangers that these clothes represented in spreading plague. In addition to discussing the Marseille and Northern Italy outbreaks, numerous historical texts reported interhuman transmission of plague through the transportation of movables, merchandise or clothes belonging to plague-stricken persons [44]. Accordingly, the people of those times saw infective potentiality in textiles and other merchandise that came from infected places or with which plague victims had been in contact. Indeed, the trafficking of the clothes of deceased plague victims was a major problem in epidemic periods, given that in preindustrial Europe, wages were so low that buying clothes was a luxury that ordinary people could only afford a few times in their lives [45]. There are multiple direct accounts of episodes in which buying, selling and stealing clothes was reported as having triggered plague in houses, small villages or even cities [41,45,46]. In Milan, the purging and sequestering of all the clothes and movable of plague victims were two primary prophylactic measures. The results herein reported can be put in perspective by those obtained on typhus by the Nobel Prize winner Charles Nicolle: during a 1909 typhus outbreak in Tunis, Nicolle demonstrated for the first time the role of body lice as vectors of infectious diseases in an epidemic context [47]. He found that patients’ lice-infested clothes and their feces infected other people [48, 49]. Nicolle noted that bathing and changing clothes stopped contagion [48]. Nicolle’s observations shed light on our results by showing that when an epidemic disease is transmitted through clothing, it is actually transmitted via human ectoparasites such as body lice, acknowledged vectors of typhus and plague [19, 48]. The fact that the vector role of lice was demonstrated in 1909, late in the history of infectious diseases, partially explains why historians neglected body lice in the interhuman transmission of plague, applying the rat-rat flea model, despite all the evidence [13].

In texts from both Marseille and Northern Italy, the words “carbuncle” and “bubo” were significantly enriched. Interestingly, the word “carbuncle”, referring to the skin ulcer following the introduction of *Y. pestis* through skin [32], seems to be very representative of the second plague pandemic, whereas it is rarely described during the third pandemic and no longer reported in contemporary cases of plague. Notably, “buboes” and “carbuncles” were localized all over the body, including the face and the neck (specifically ear, jugular, parotid gland, and maxillary regions), localizations that do not support a role for fleas because rodent fleas are known to bite mainly the lower part of the body (legs and thighs) [Barbieri in press, 2020]; rather, they suggest a role for human ectoparasites, including body lice and human fleas, as vectors of Y*. pestis*, which could explain the rapid spread of the epidemics [19,50–52], as supported by epidemiological [13, 17], historical [53] and archaeological studies [18].

In summary, new data generated by a mostly unbiased lexicographic tool provided contrasting knowledge regarding the dynamics of the second plague pandemic, challenging the proposed hypothesis that these episodes resulted from the reactivation of local plague foci [6] by suggesting human ectoparasites as vectors of the deadliest and most widespread plague pandemic in human history.

## Materials and Methods

### Ancient texts

Texts were retrieved from Google Books, Archive.org and Buisante.parisdescartes using the following keyword combinations: plague & Marseille; plague & Italy with specific period ranges such as plague & Marseille & 1720-1820 or plague & Italy & 1629-1680. A total of 32 historical texts describing the 1720-1722 Great Plague of Marseille (16 complete books) and the 1629-1631 Italian plague, also known as the Manzoni plague, after the famous novel written by Alessandro Manzoni in 1827 [54] (13 complete books and 3 book chapters), were retrieved (Data files S1 and S6). Additionally, we retrieved 11 contemporary negative control texts for Marseille and 20 contemporary negative control texts for Northern Italy, all written during the 17^th^-18^th^ century and describing surgery, plants, universal history, industry and architecture; these texts were scanned within the framework of the “Google Books” program undertaken by Google and the Archives program (Data files S2 and S7).

### French texts’ disposition

To filter the raw text words (see the historical text mining analysis section), we used a list of French words comprising 336,631 words, conjugated verbs and proper nouns compiled from the Français-Gutenberg Dictionary. The symbol “⎧” (“long S”, which was used in most languages in Europe until the industrial revolution) was automatically substituted with an “s” only if the new word with an s existed on the list of French words. To standardize the words, plural forms ending with a “s” or “x” were converted to the singular form only if the singular word was present on the list of French words.

### Italian texts’ disposition

Italian texts (plague-related texts and controls) presented ambiguities that could lead to errors in the analysis. Indeed, most Italian books present, at the upper part of each page, the name of the book or chapter. Another problem of Italian books is the repetition of the last word of each page at the beginning of the next one. To avoid problems due to the overrepresentation of some words, these two types of repetitions were manually corrected by removing the name of the book/chapter from each page, together with the last word of each page. As reported above, the symbol “⎧” (“long S”) was automatically substituted with an “s”. However, we observed that optical character recognition (OCR) programs tend to recognize the “⎧” symbol as an “f” and not as an “s”. To avoid the loss of the Italian word “peste” (“plague”) due to OCR mistranscription, the word “pefte” (no meaning) was changed to “peste” in all the texts. Accordingly, the only words identified were those present on the list of Italian words (including singular and plural forms, conjugated verbs, proper names, and surnames = 931,657 words). Of these words, 427 (32%) were discarded. Moreover, one-letter words were removed, and singular and plural forms of the words were unified as best as possible into only one word following Italian grammar rules for plural formation (Supplementary Text 3). Plague-related and control text word counts were compared using the DESeq2 R package with default parameters (version 1.22.2).

### Historical text mining analyses

After the raw texts were imported into R (see Supplementary data) in .txt format, all the words were separated and converted to lowercase; accents, empty cells and non-letter characters such as “?”, “!”, “,”, “[.]”, “-”, “_”, “%”, “$”, “€”, “#”, “\\”, “+”, “*”, “:”, “;”, “>”, “<”, “§”, “&”, “•”, “«”, “»”, “[{}]”,”’”, “\”, and “‘” were removed. The number of occurrences was computed for each word in the 63 texts. Regarding the plague-related texts, 3,624,476 words were identified, with 2,049,566 words from the French texts and 1,574,910 words from the Italian texts. Given that we were not able to systematically repair ancient mistranscribed words, only words matching those in exhaustive list of French words (including singular and plural forms and all conjugated verbs = 336,631 words) and the exhaustive list of French cities, towns and places or Italian words were considered. Of ∼ 3.5 million words, 2,721,742 were conserved (75%), resulting in 25,659 and 44,475 unique words for the French and Italian texts, respectively. Then, we applied a cut-off value for removing all unique words present fewer than 5 times in the French and Italian texts preliminary to the DESeq2 analysis, resulting in 10,315 and 10,645 unique words for the French and Italian texts, respectively. Subsequently, plague-related and control text word counts were compared using the DESeq2 R package with default parameters (version 1.22.2). This approach identified 70 French and 147 Italian terms that were overrepresented in plague-related texts compared to control ones (*P* adjust < 0.05 and log2foldchange >0) and 213 French and 313 Italian words that were present > 4 times in plague-related texts compared to controls (Log2foldchange >2) (Data files S3 and S8).

### Nested analysis

In a second step, we used a “nested” analysis to identify the words that lay in close proximity to overrepresented words in the plague-related texts. We extracted words located +/-25 words apart from significant terms and compared them to the remaining part of the texts using DESeq2. Significant words (adjusted *P* value ≤ 0.001 and log2FoldChange > 0) were plotted using the “wordcloud” R package (version 2.6) (Data files S5 and S9).

### Word translation

All the French and Italian words were translated into English using the translation function in Microsoft® Word. Each overrepresented (only the words with *P* < 0.05 or log2foldchange > 0) and nested (words associated with overrepresented words) words were controlled by the operators for mistranslation and then corrected. Words with no English translation were maintained as they were; this was the case for one French word, “corbeau” (a person responsible for burying plague victims), and 4 Italian words, namely, “monatti” (people responsible for burying plague victims), “espurgatori” (people responsible for stuff purging), “untori” (people accused of intentionally spreading the disease using “special” ointments), “caldiera” (a tool used to untie the cocoons of silkworms), with no direct English translation. Words exhibiting an “out of context” translation were also manually corrected, such as “carbone”, “tacchi”, and “unti” in Italian and “preservatif” in French texts.

### Network Representation using Gephi

Graphical representation of the nested analysis results was performed using the software Gephi 0.9.2 [55]. This software was used exclusively for representation purposes of the data and statistics obtained by the nested analysis. The results of this analysis were converted into nodes and edges to generate the network. At each node, a word was obtained from the nested analysis, and the edges between two words represented the association between these words. Only words with a strong statistical significance (*Padj* < 0.01) were represented. The spatial position of the words in the network has no statistical meaning. To maximize the efficiency of the comparison between the French and Italian datasets, for the construction of the comparison networks, the words nested with the word of interest and the words with which the word of interest was nested were used. Therefore, an edge was created each time a word of interest was associated with another word with a significance determined by a *P*adj < 0.01.

## H2: Supplementary Materials

### Supplementary materials and methods

Fig. S1. Jaccard index analyses for French texts corpus (A) (plague related-texts and control texts) and for Italian texts corpus (B).

Fig. S2. Volcano plot obtained from the analysis of the French texts corpus (A) and Italian texts corpus (B), generated using the DESeq2 package (log2foldchange >0 and P adjust<=0.05).

Fig. S3. WordClouds generated from the nested analysis of French plague-related overrepresented words.

Fig. S4. WordClouds generated from the nested analysis of Italian plague-related overrepresented words.

Fig. S5. Word network representation for selected words from the Great Plague of Marseille (1720-1722).

Fig. S6. Word network representation of words associated to potential sources of plague.

Table S1. Statistical table summarizing the number of raw, filtered, used and unique words for each French plague-related texts and control texts.

Table S2. Statistical table summarizing the number of raw, filtered, used and unique words for each Italian plague-related texts and control texts.

Data files S1. Historical French text related to the 1720-1722 Great Plague of Marseille.

Data files S2. Plague-unrelated French texts dating from the same historical period of the Great Plague of Marseille which were used as controls.

Data files S3. All French words raw data generated by Deseq2 analysis.

Data files S4. List of the overrepresented plague-related words with specific classification into 19 categories of words.

Data files S5. All French words raw data generated by “nested” Deseq2 analysis.

Data files S6. Historical Italian text related to the 1629-1631 plague epidemic in Northern Italy.

Data files S7. Plague-unrelated Italian texts dating from the same historical period of the plague epidemic in Northern Italy which were used as controls.

Data files S8. All Italian words raw data generated by Deseq2 analysis.

Data files S9. All Italian words raw data generated by “nested” Deseq2 analysis.

## Acknowledgments

### Funding

This work was supported by the French Government under the Investissements d’avenir (Investments for the Future) program managed by the Agence Nationale de la Recherche (ANR, fr: National Agency for Research) (reference: Méditerranée Infection 10-IAHU-03). This work was supported by Région Le Sud (Provence Alpes Côte d’Azur) and European funding FEDER IHU PRIMMI. The authors acknowledge Pierre Pontarotti (IHU-CNRS) for fruitful discussions about text analysis.

### Author contributions

R.B. and R.N. performed data processing and analysis, interpreted the data, drafted the manuscript, and prepared figures. M.S. and S.E. interpreted the data and drafted the manuscript. D.R. and M.D. designed the study, interpreted the data, and drafted the manuscript. All authors reviewed the manuscript.

### Data and materials availability

All data needed to evaluate the conclusions in the paper are present in the paper and/or the Supplementary Materials.

---

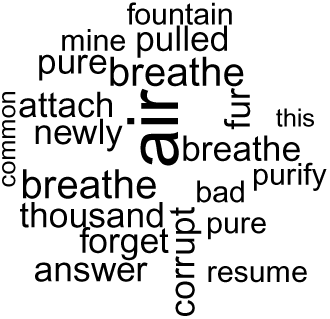
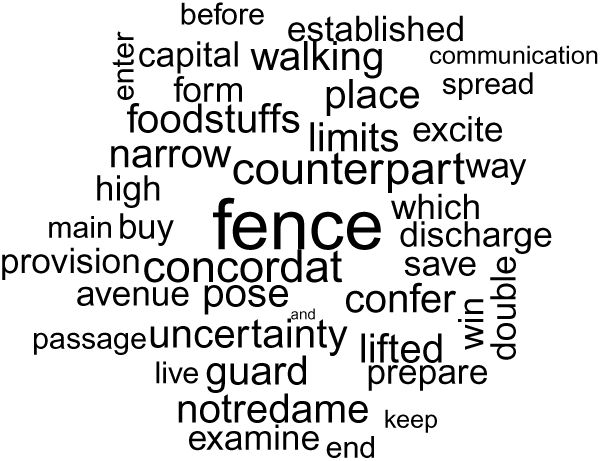
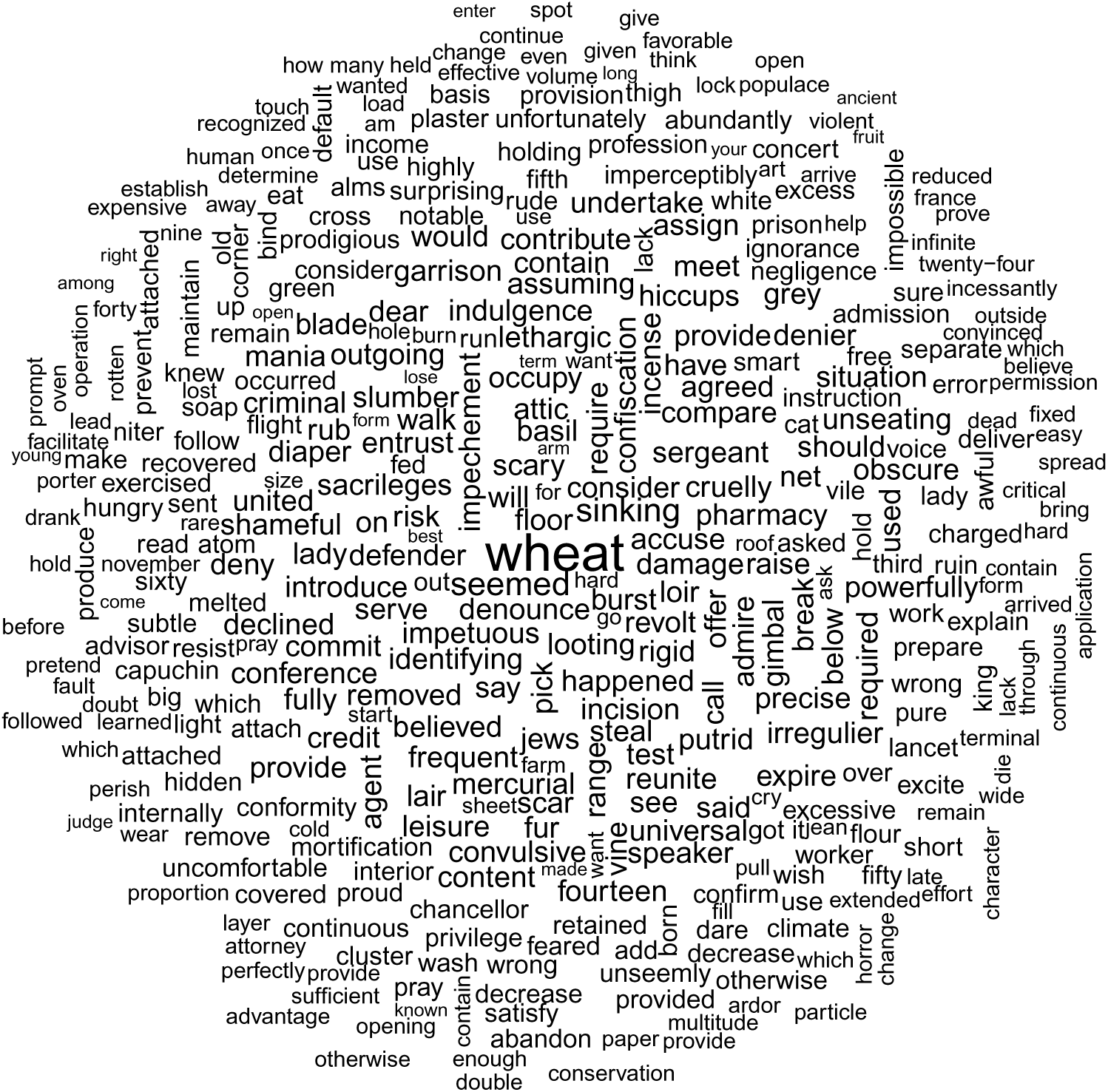
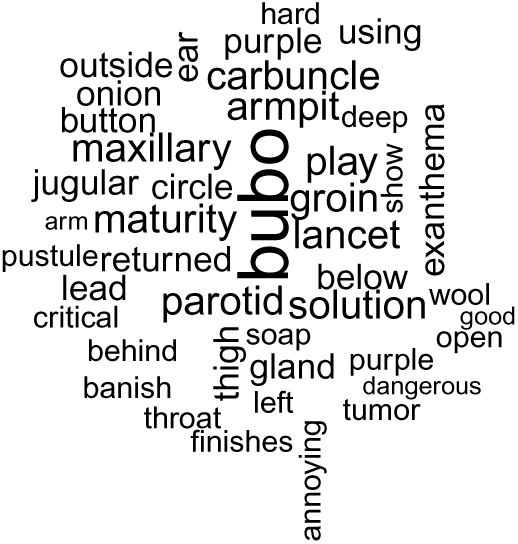
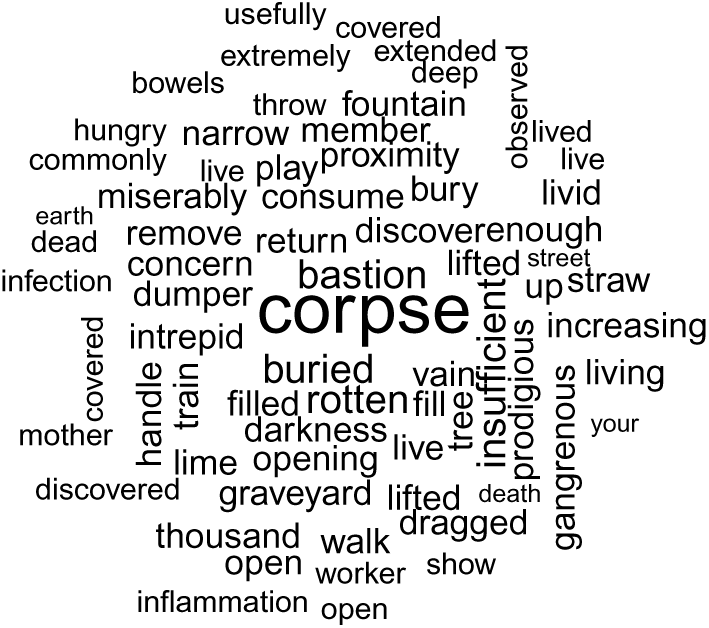
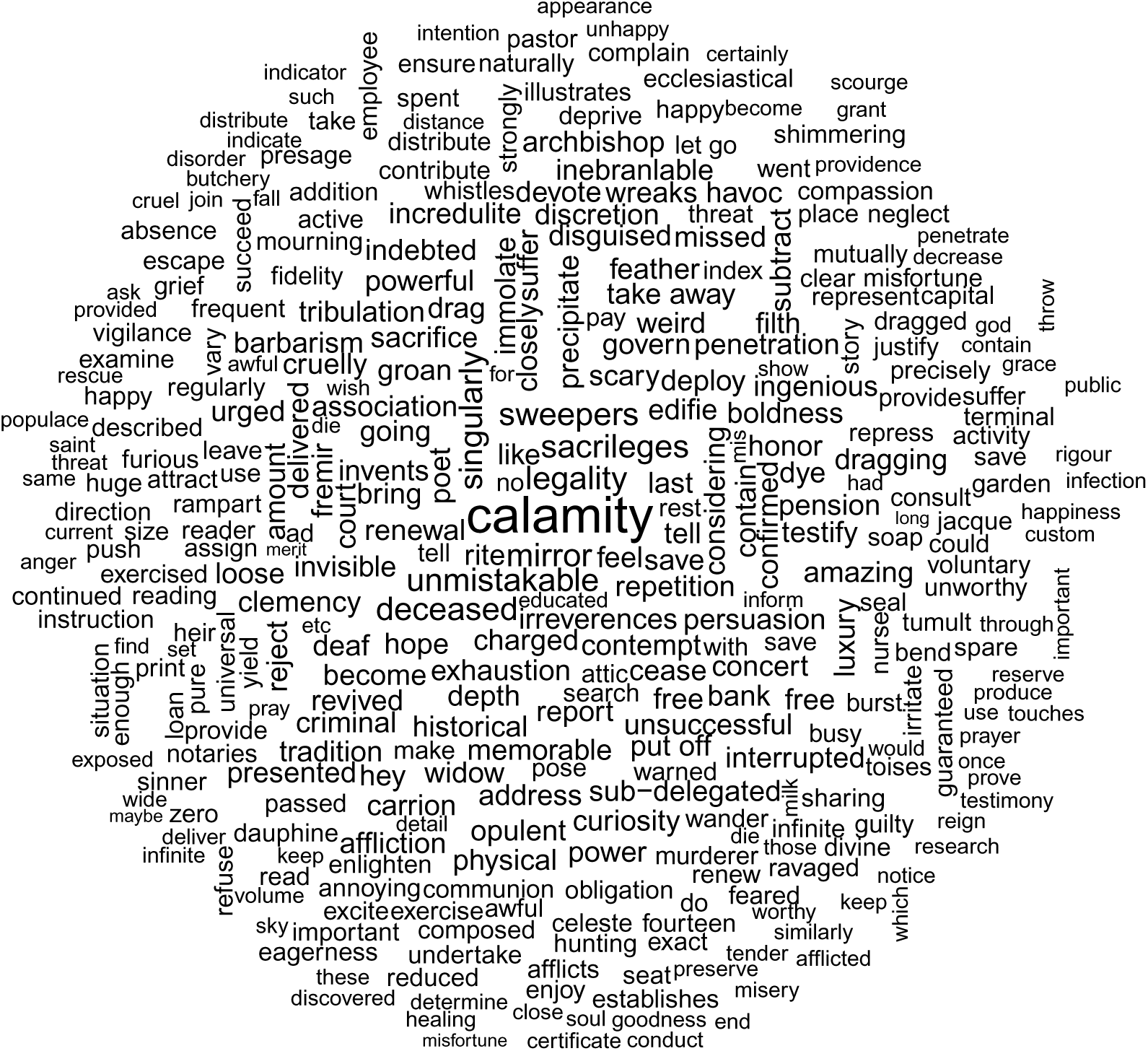
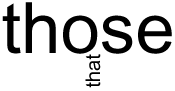
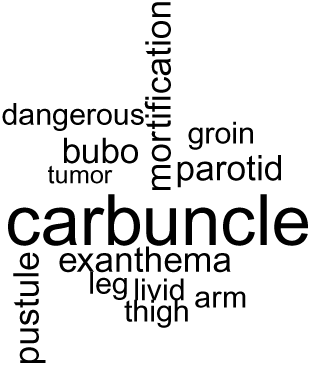
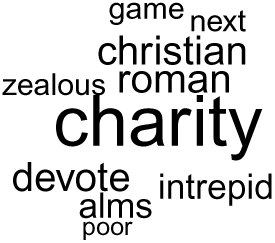
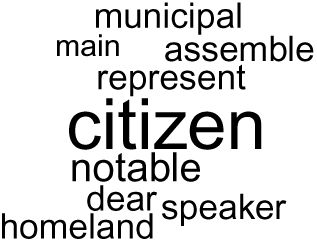
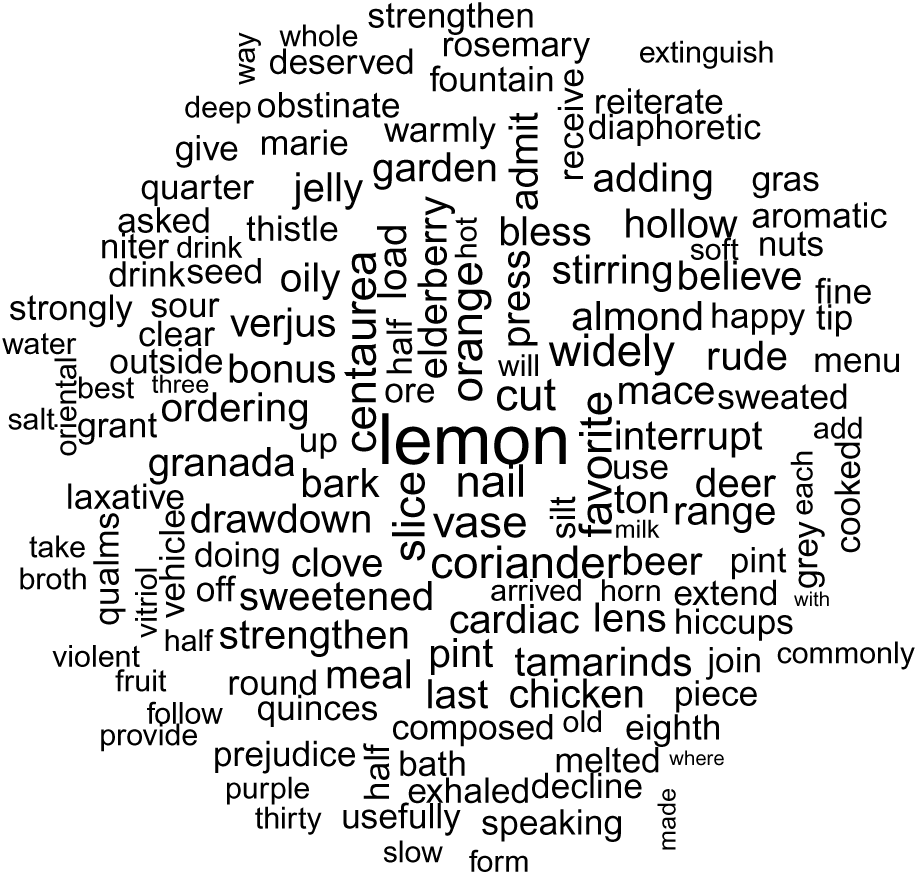
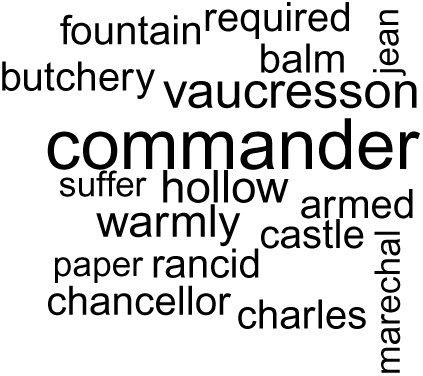
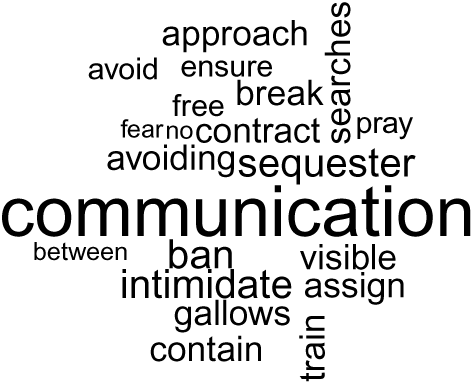
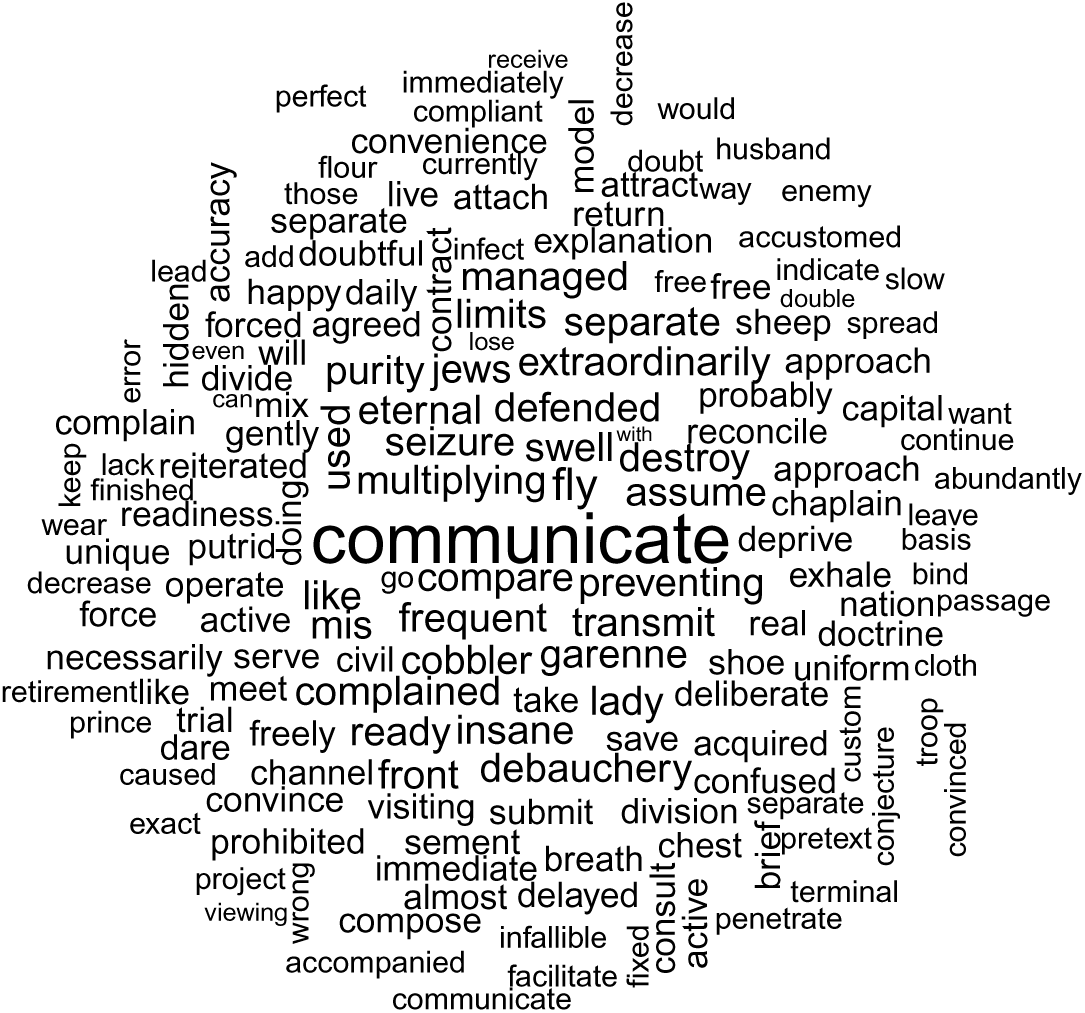
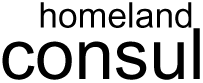
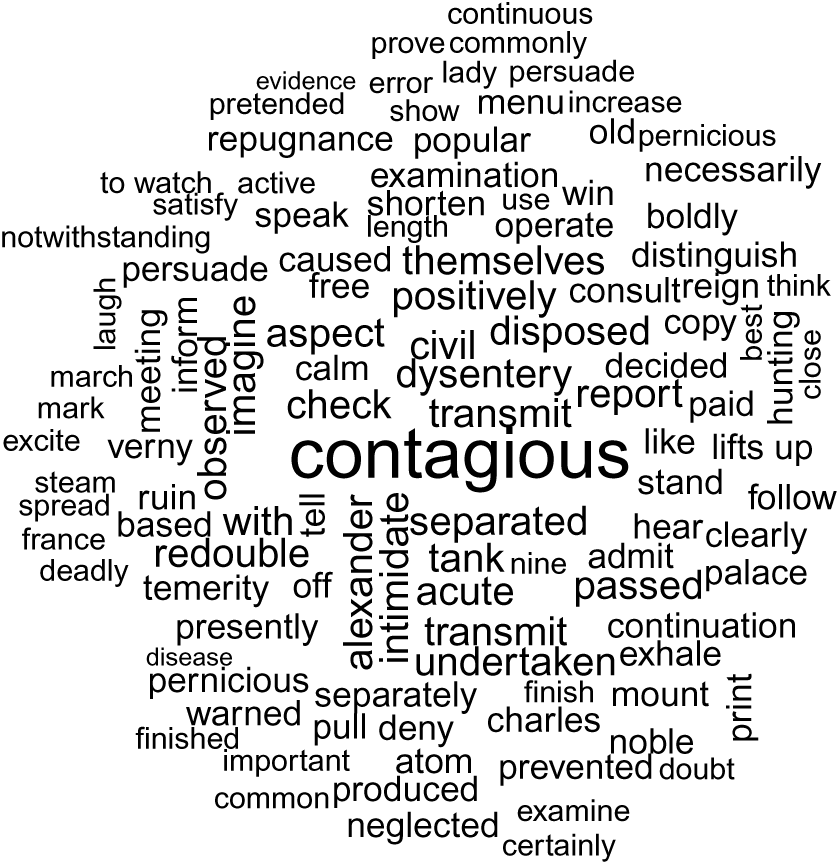
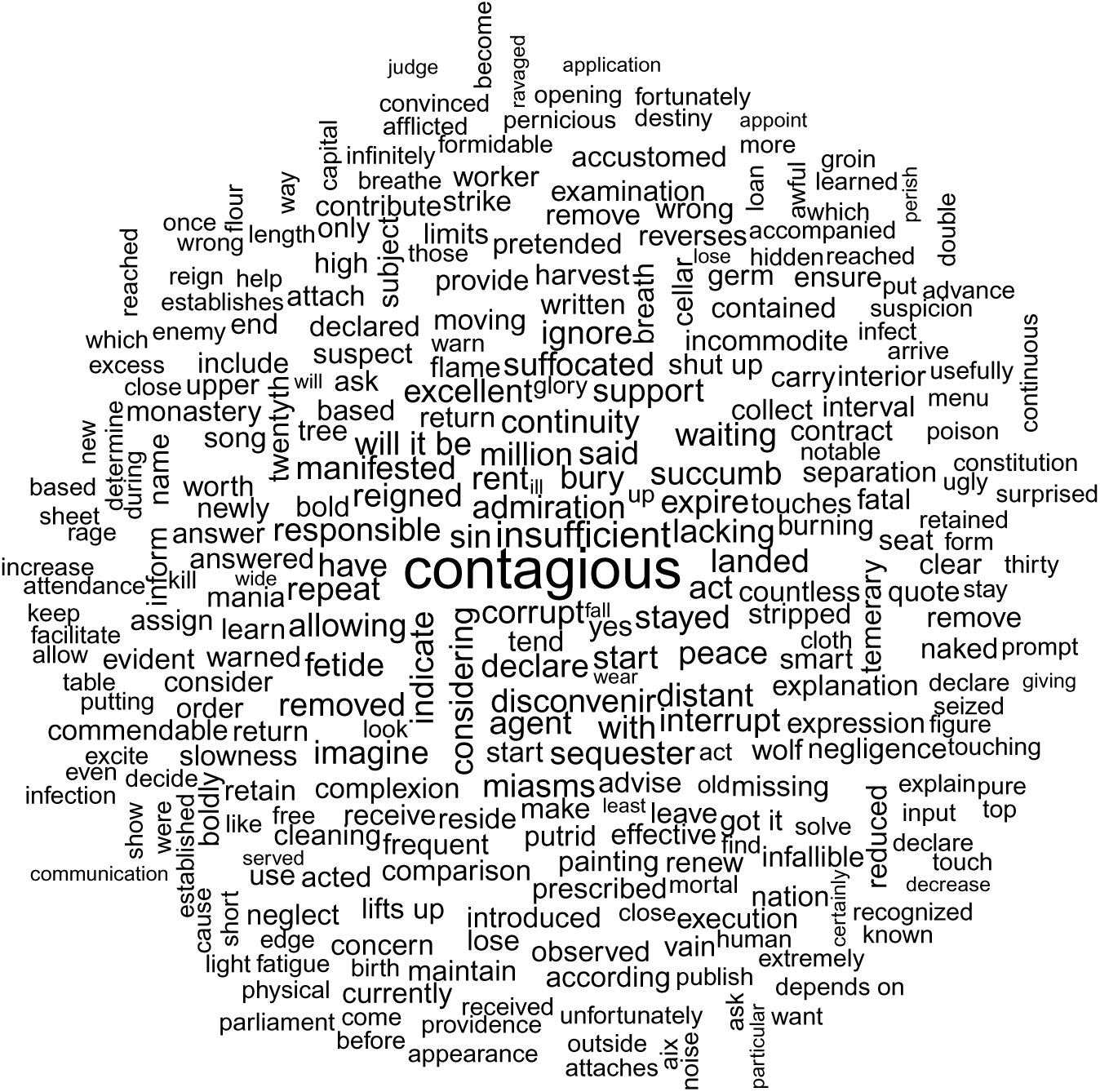
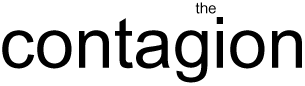
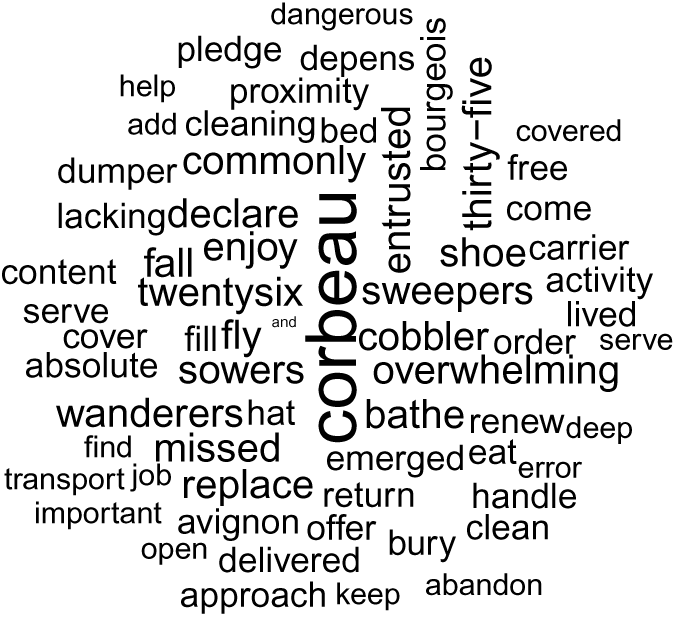
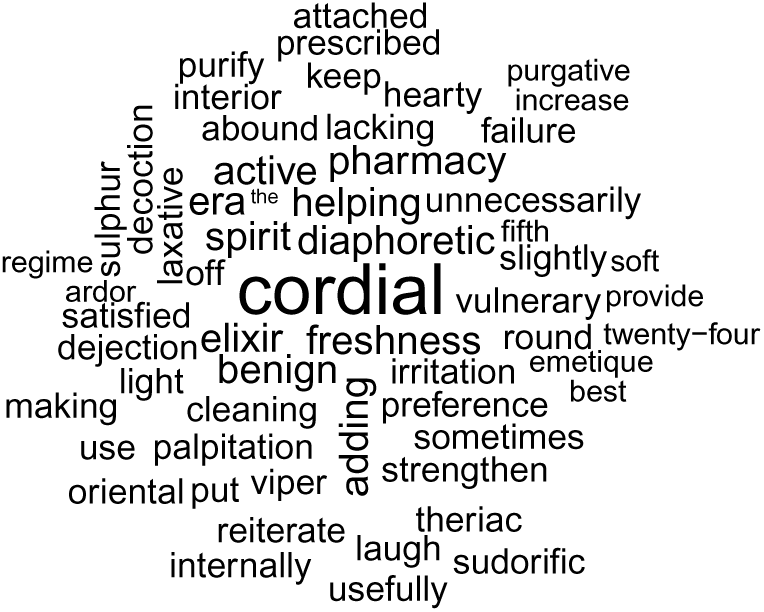
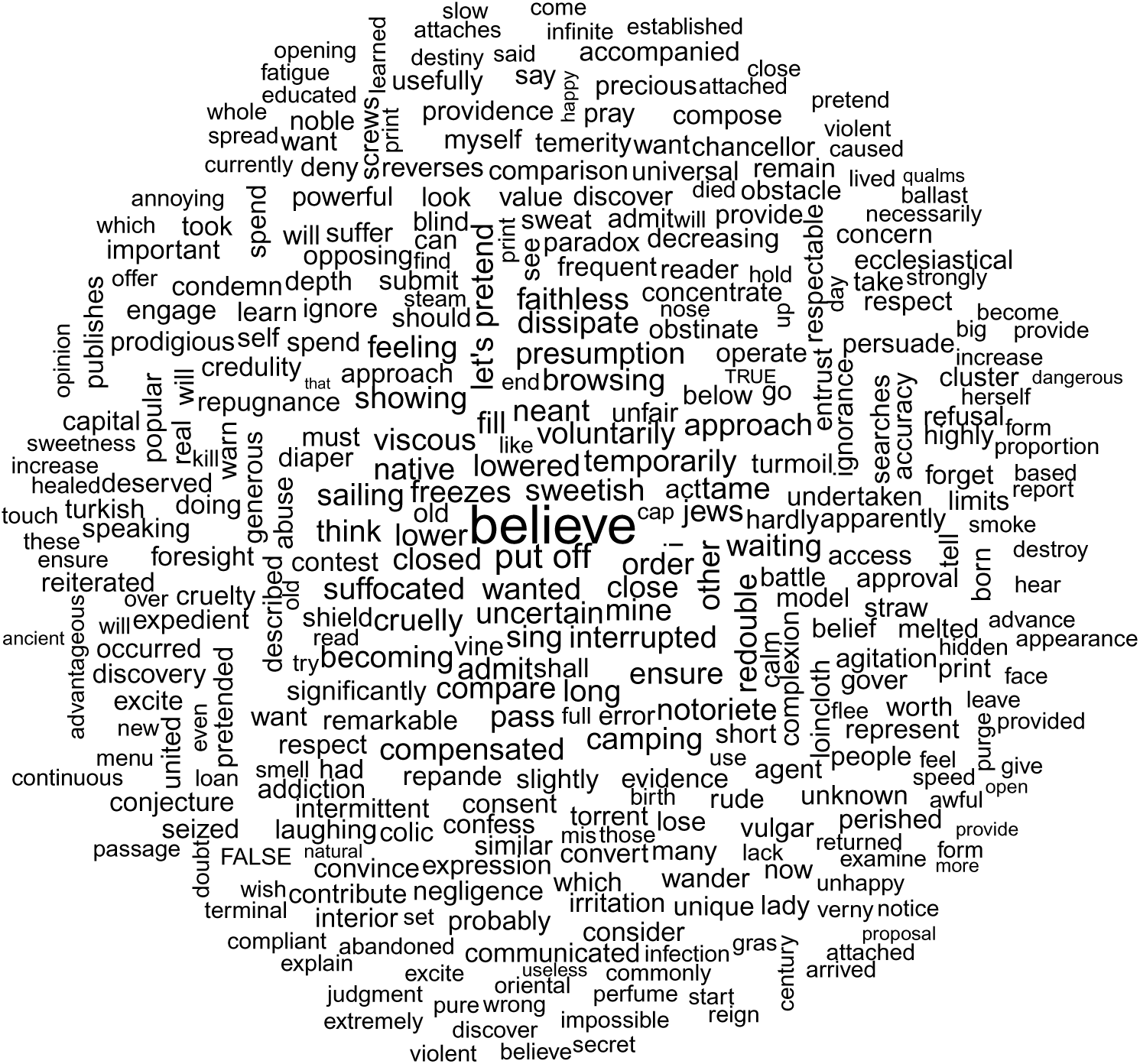
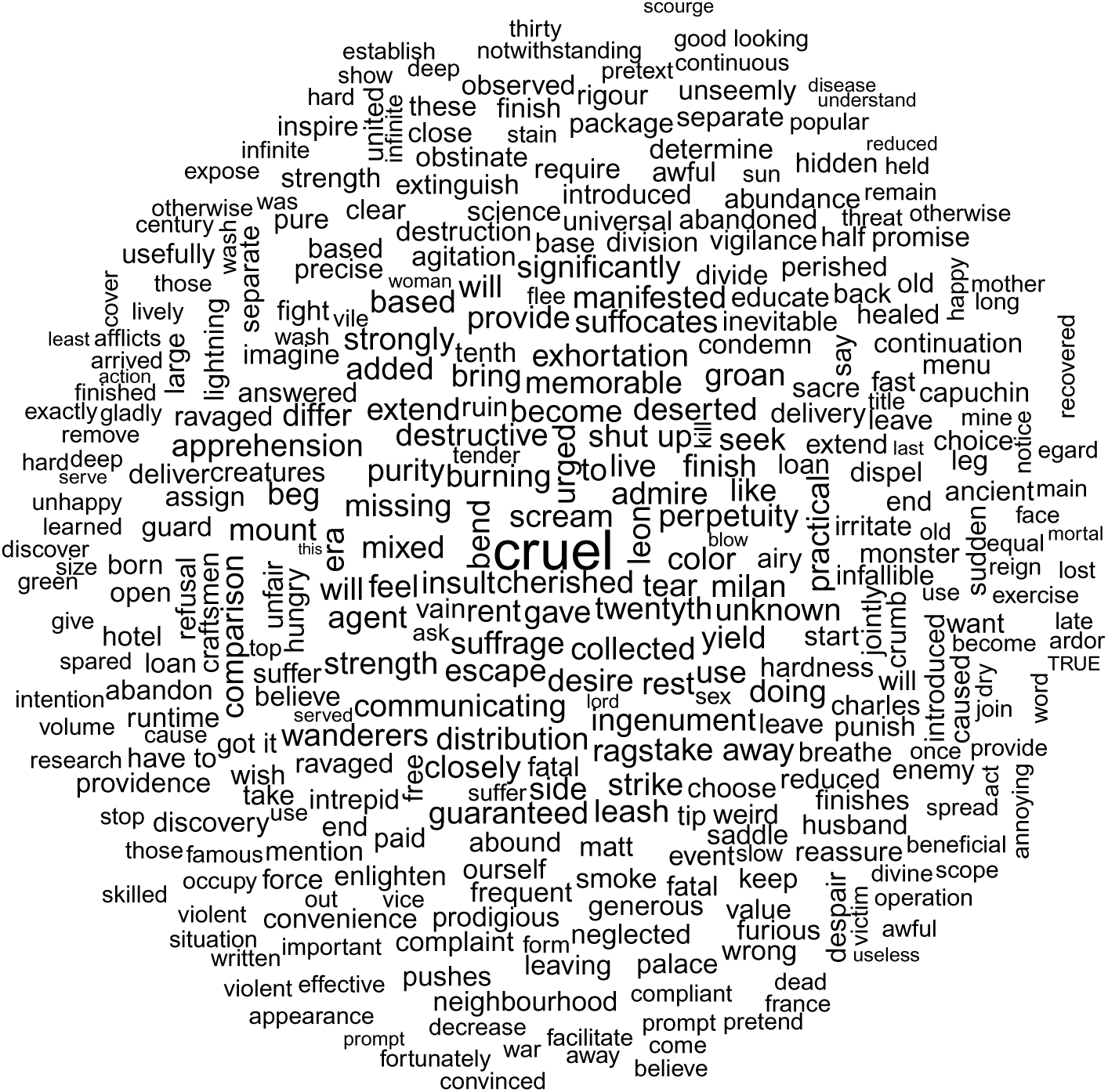
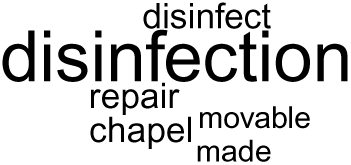
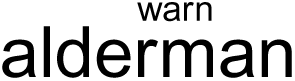
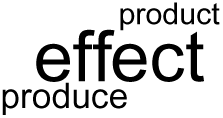
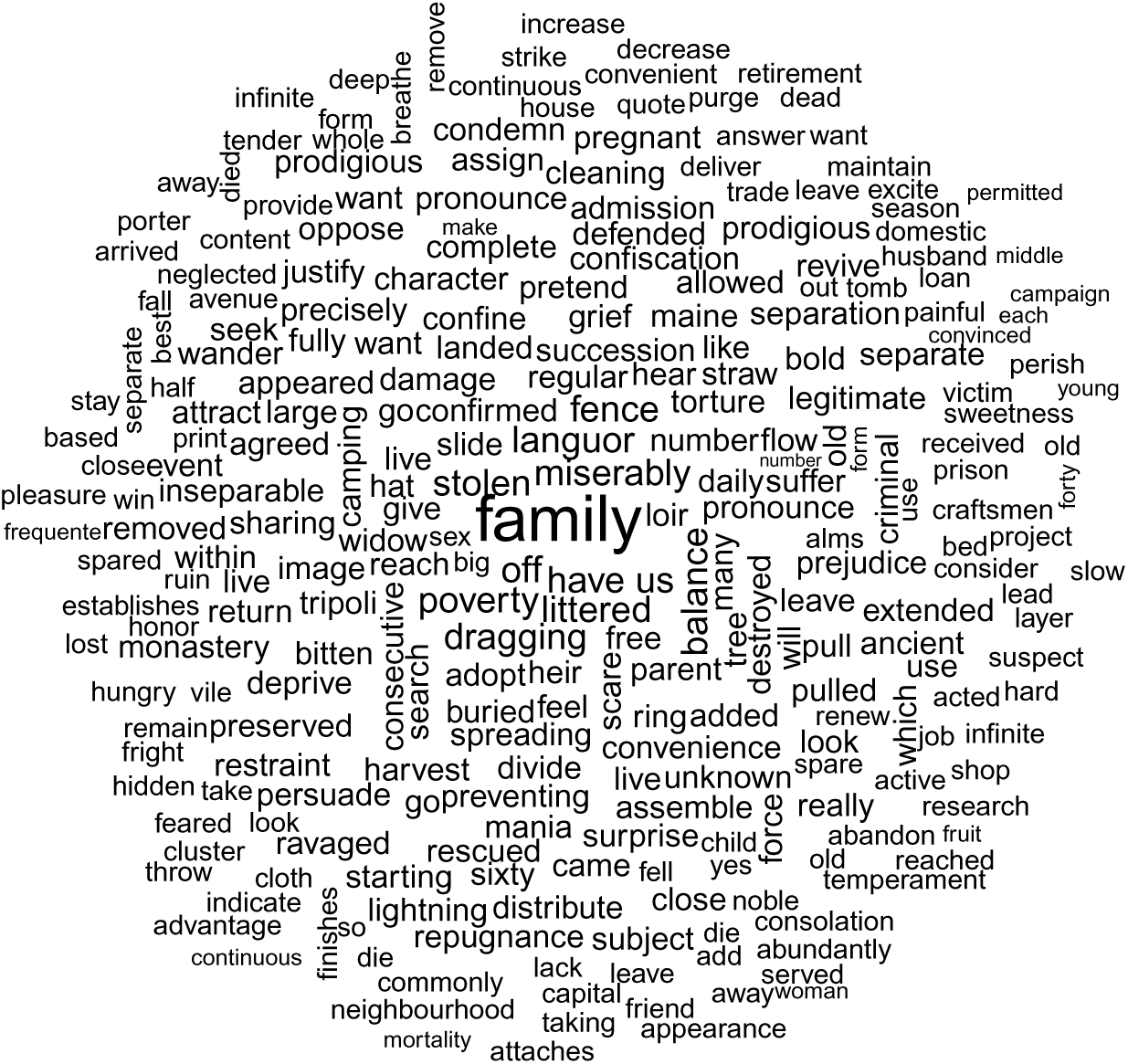
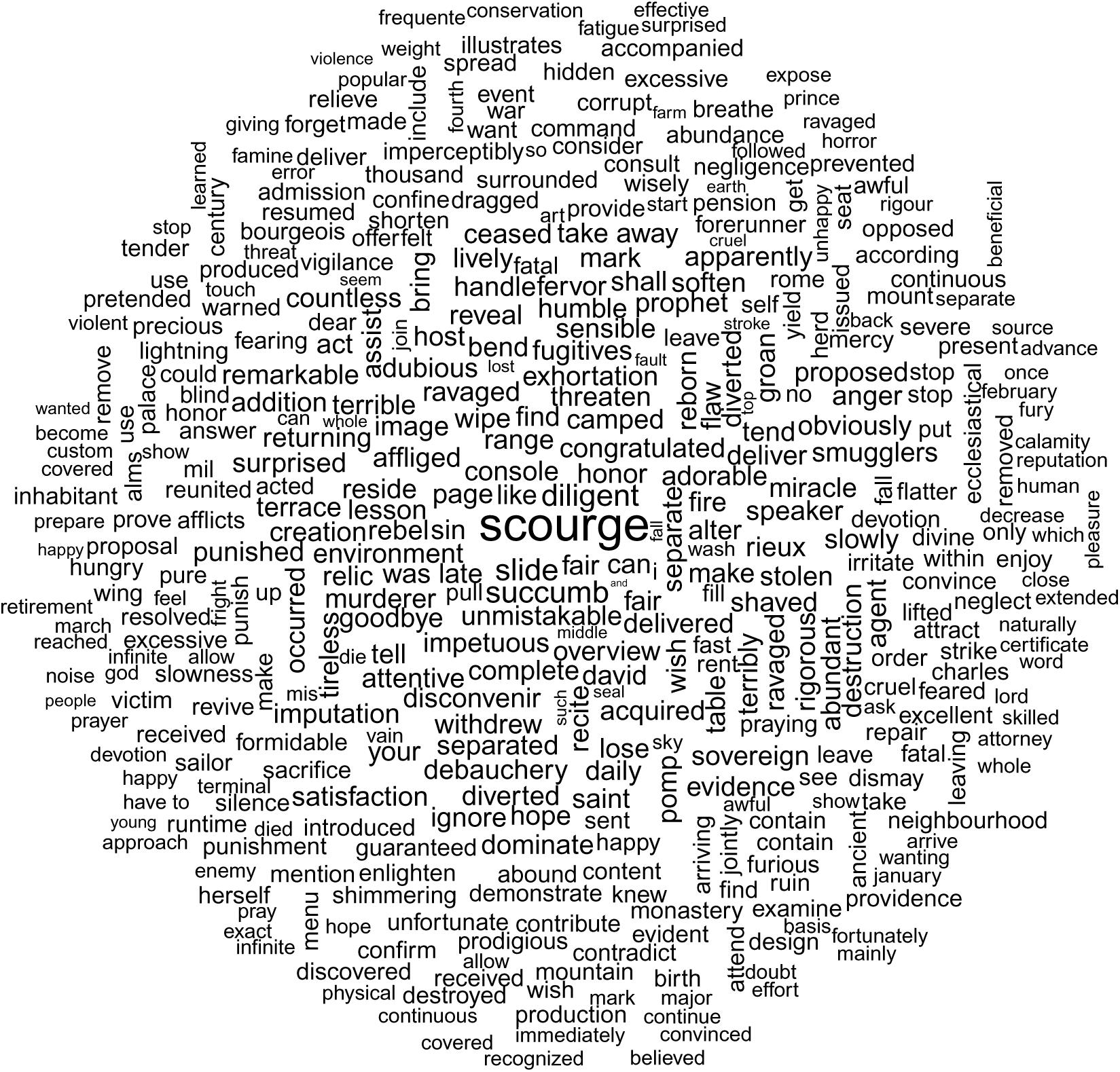
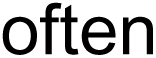
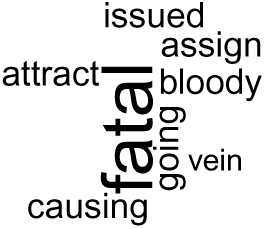
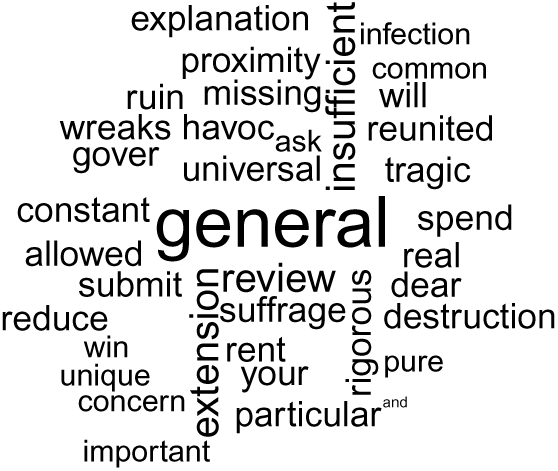
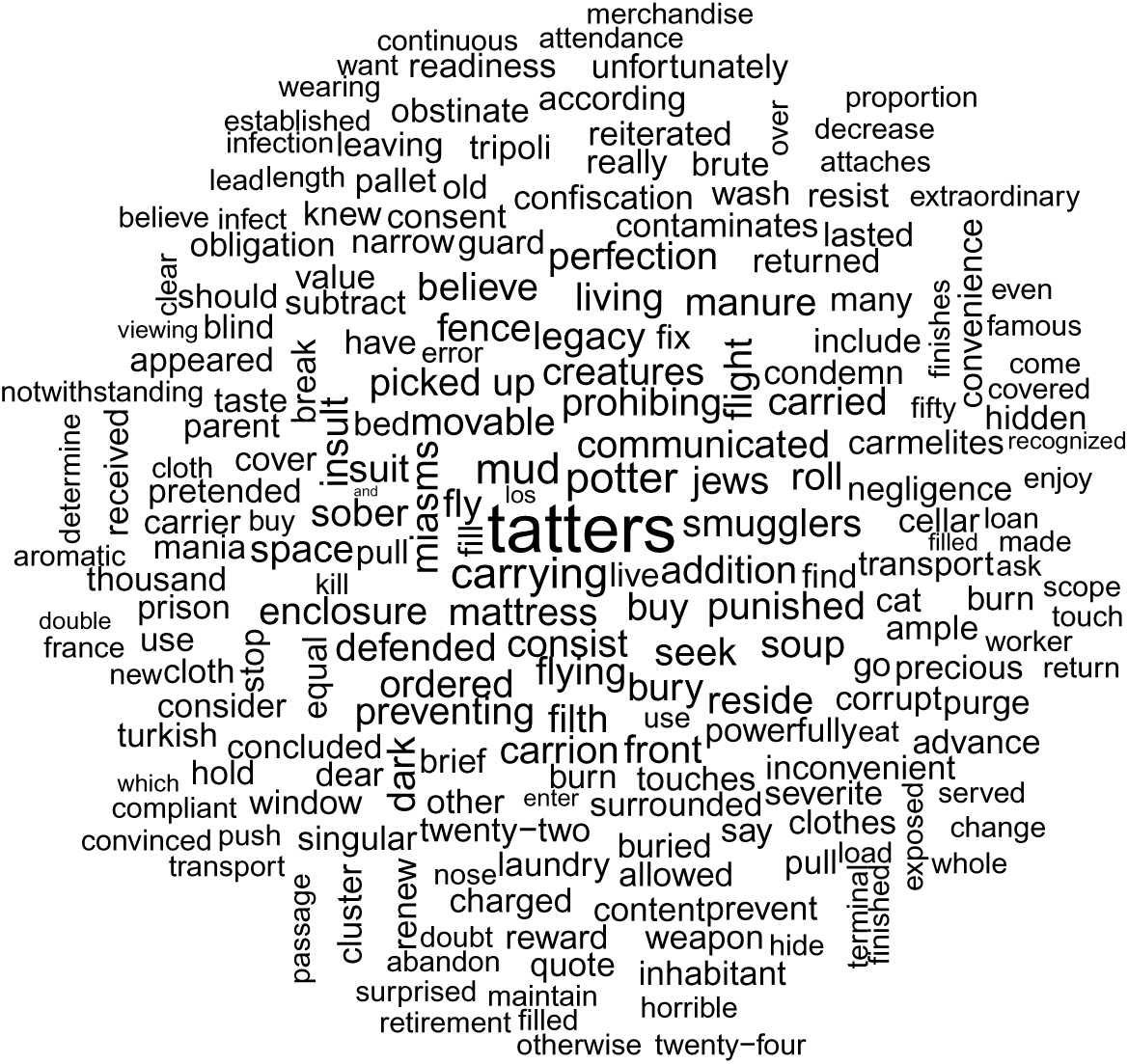
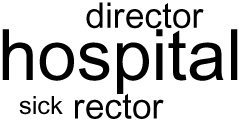
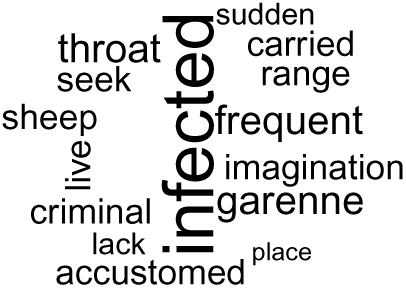
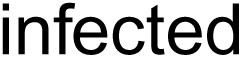
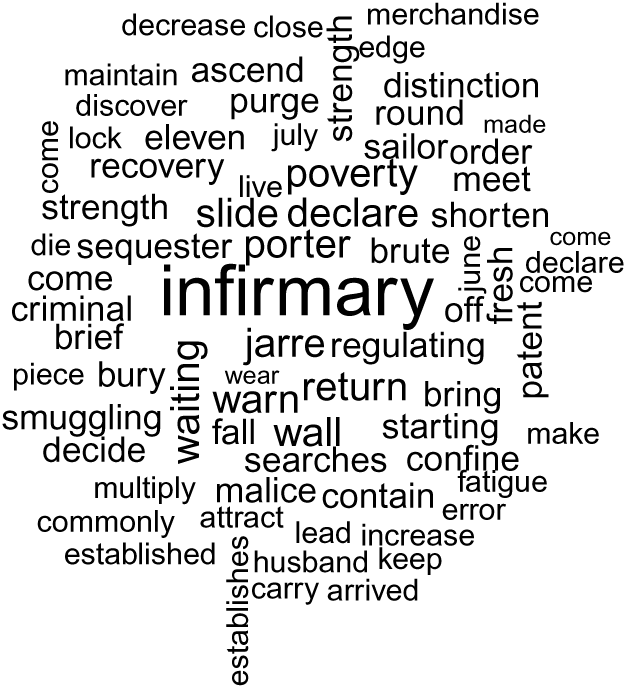
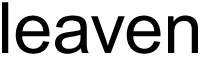
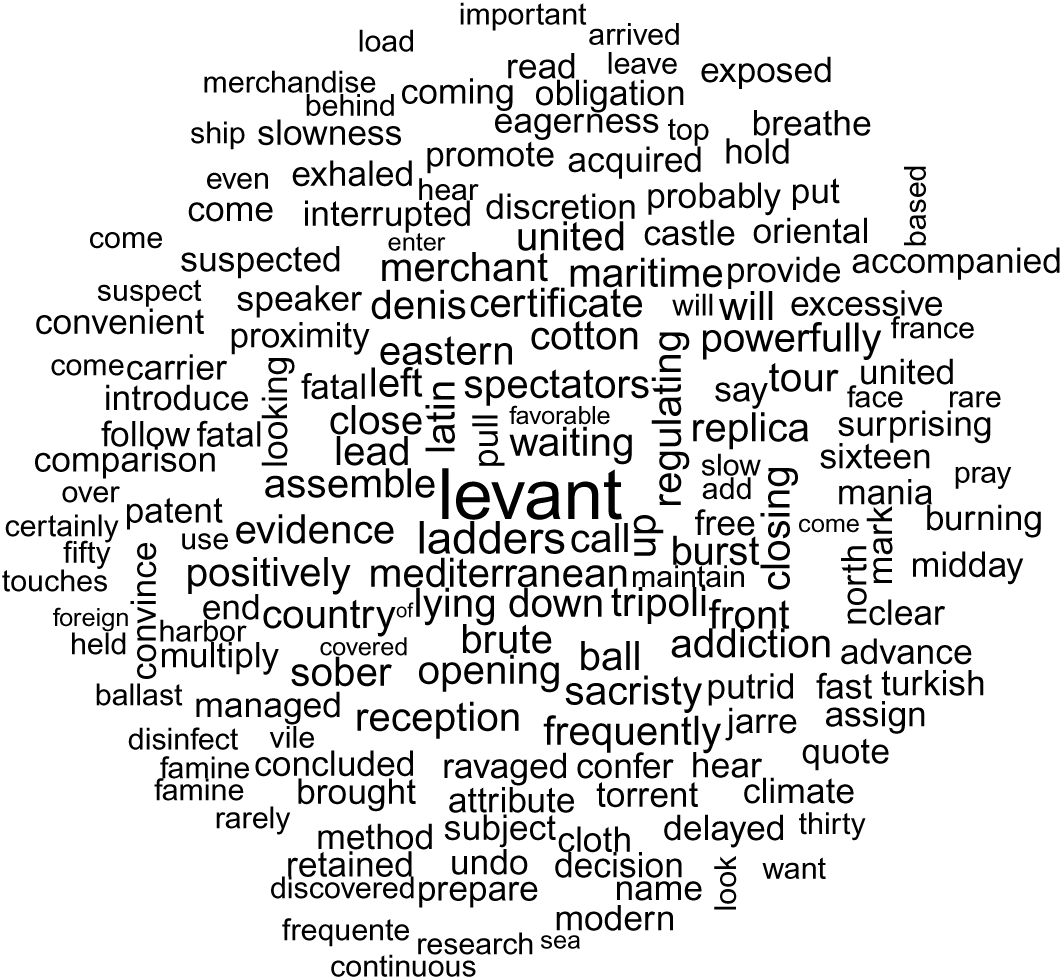
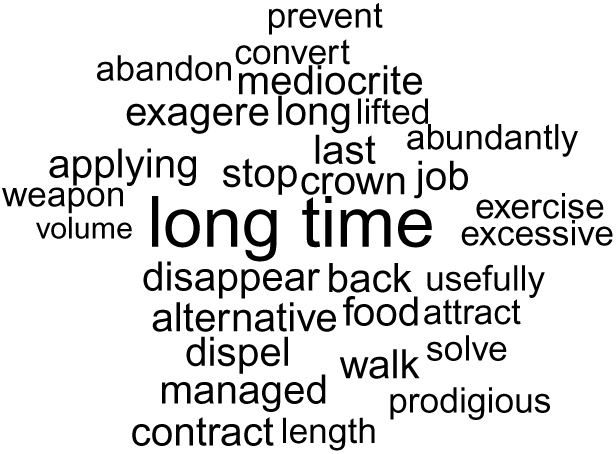
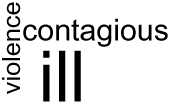
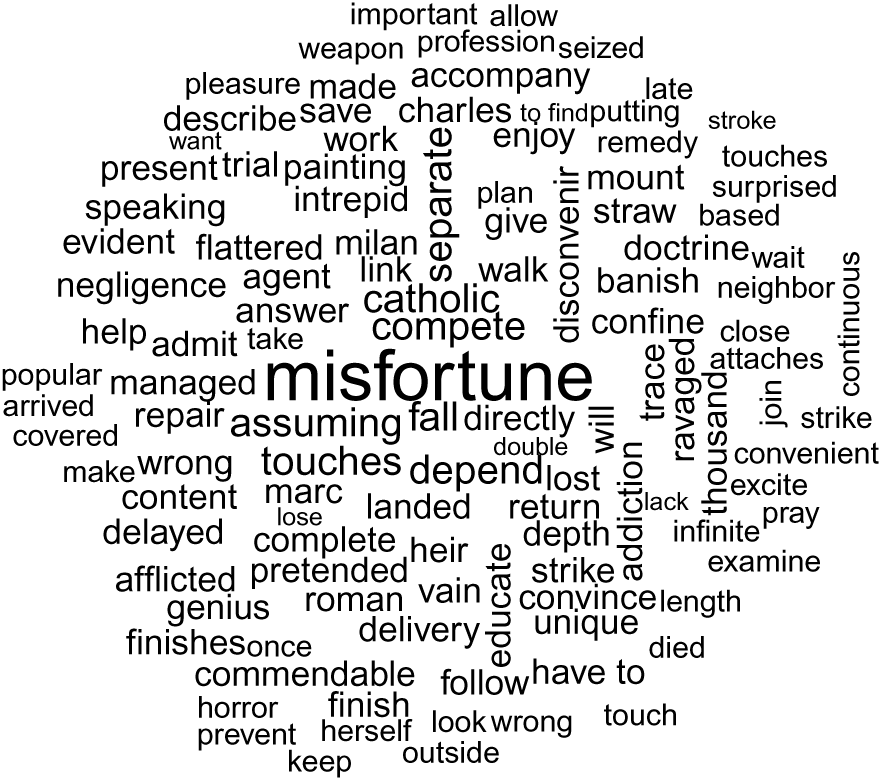
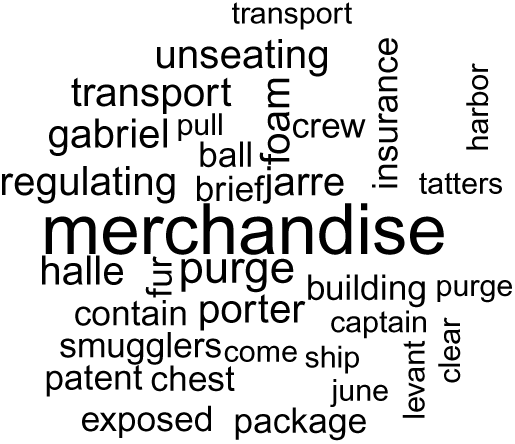
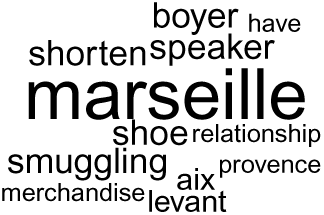
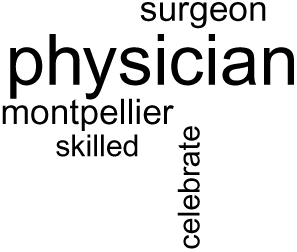
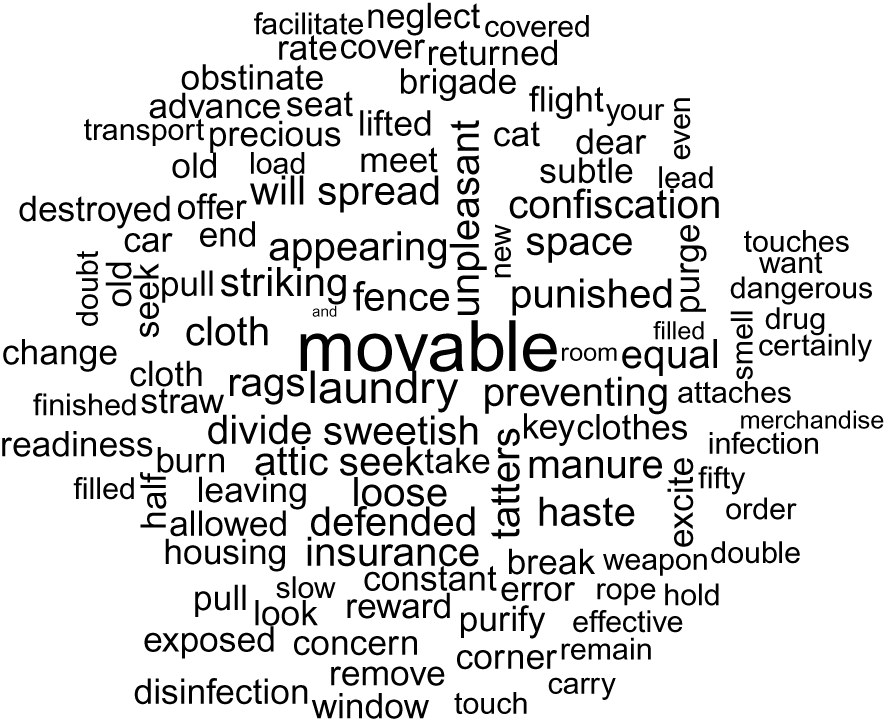
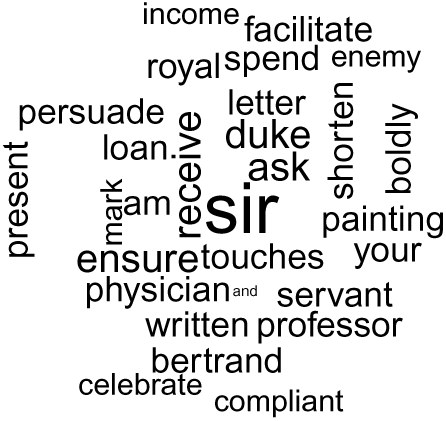
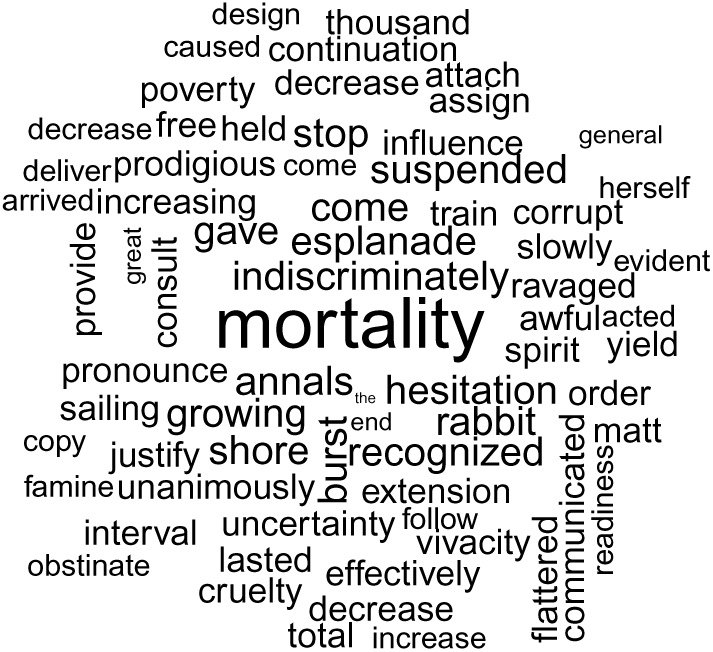
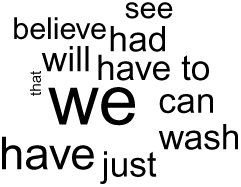
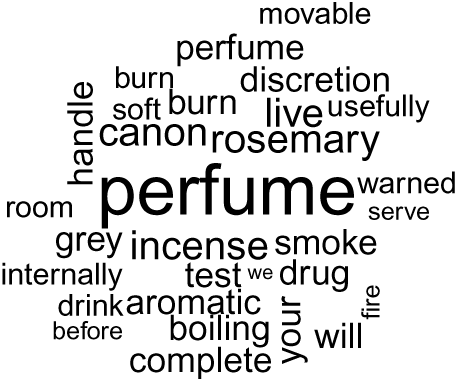
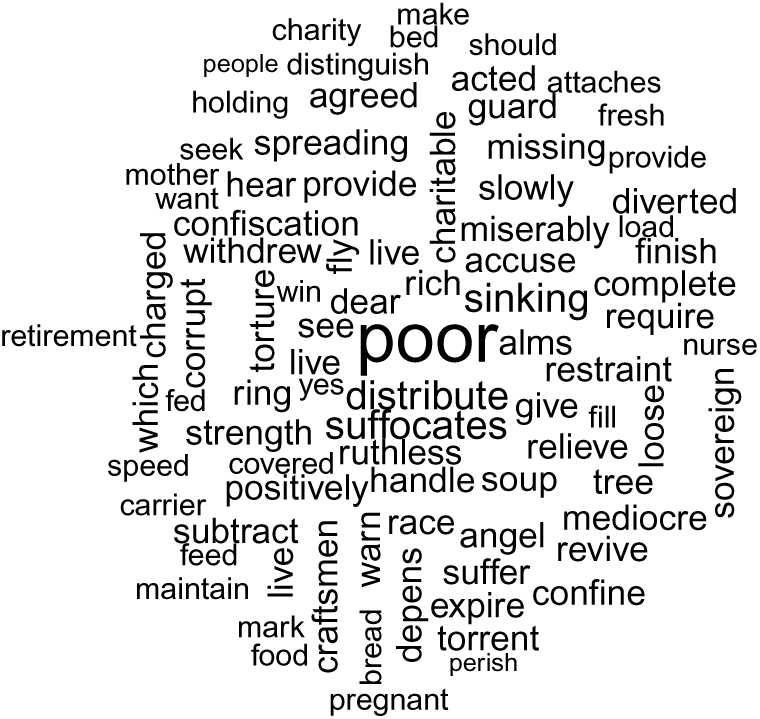
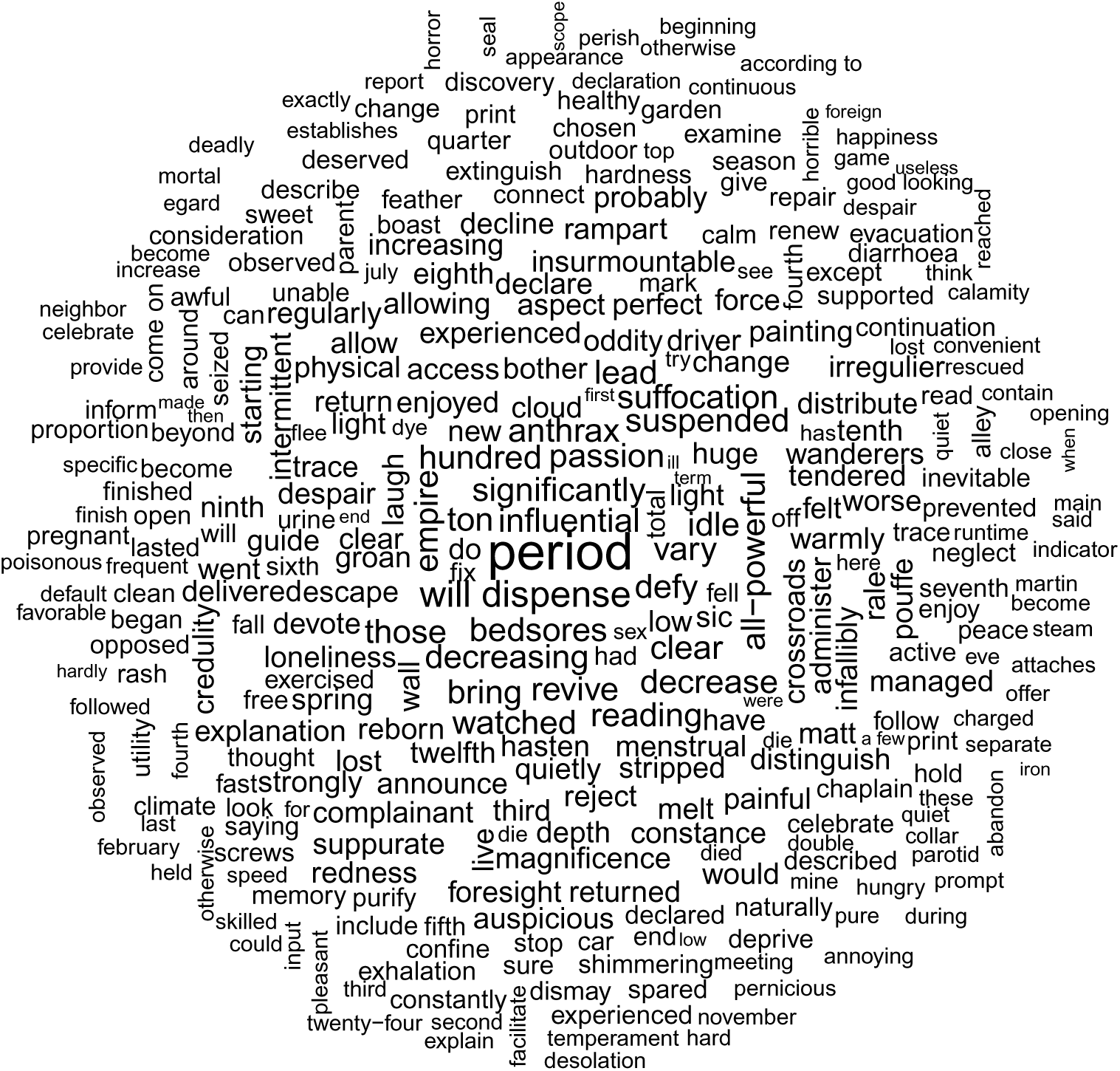
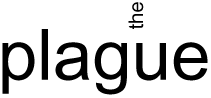
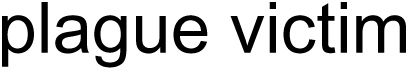
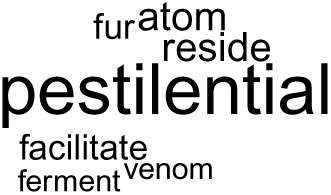
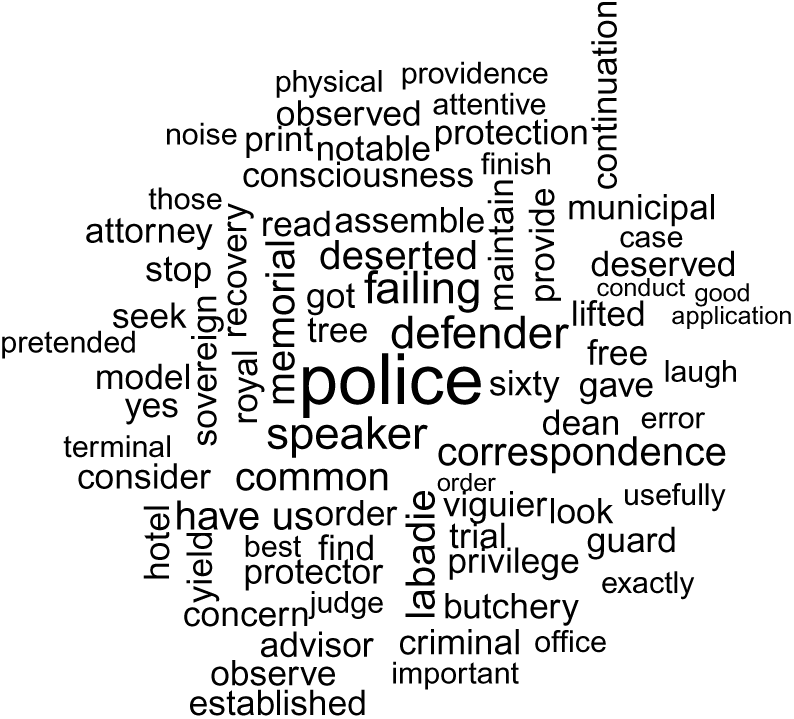
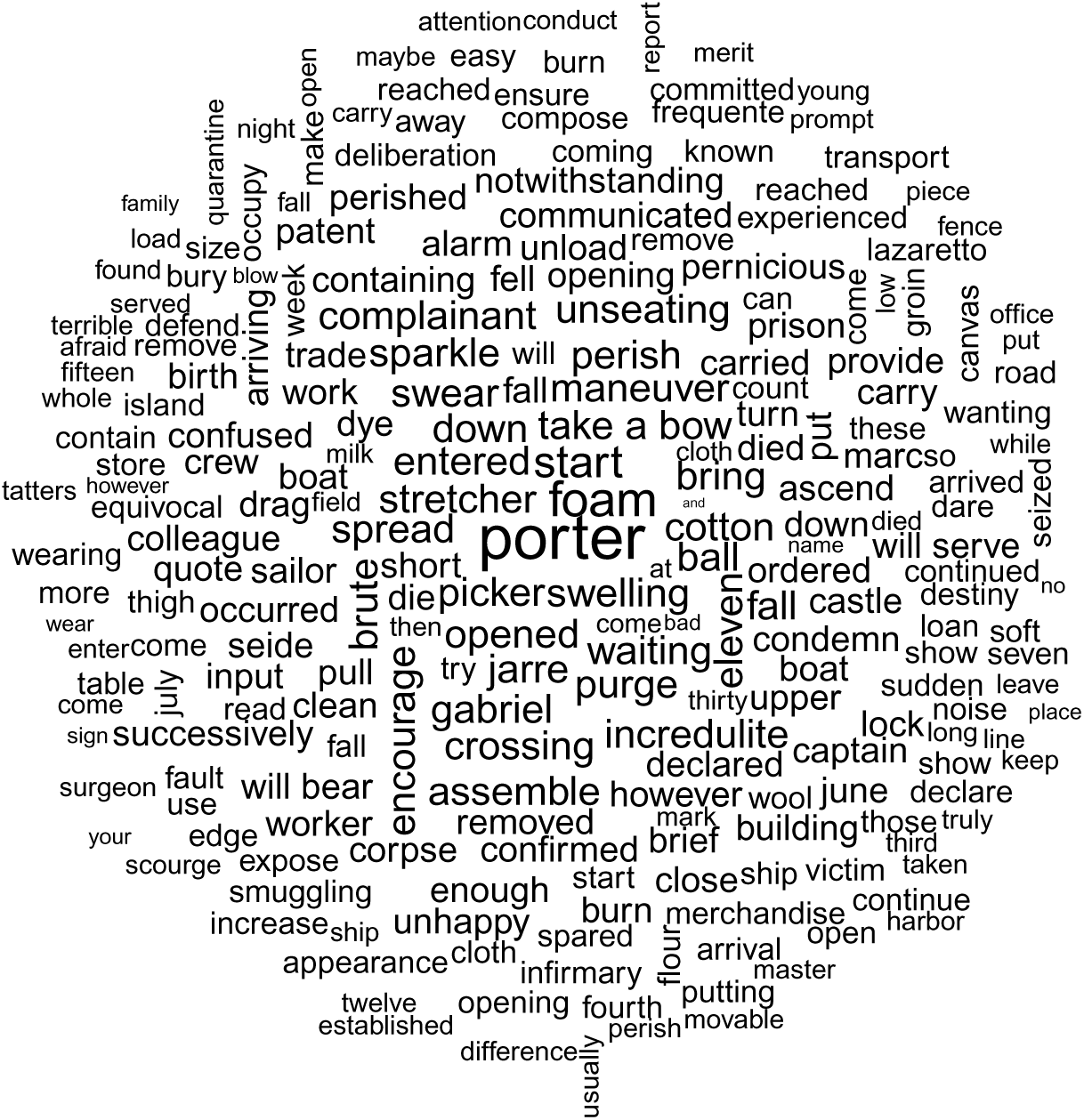
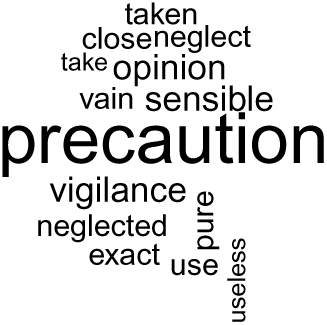
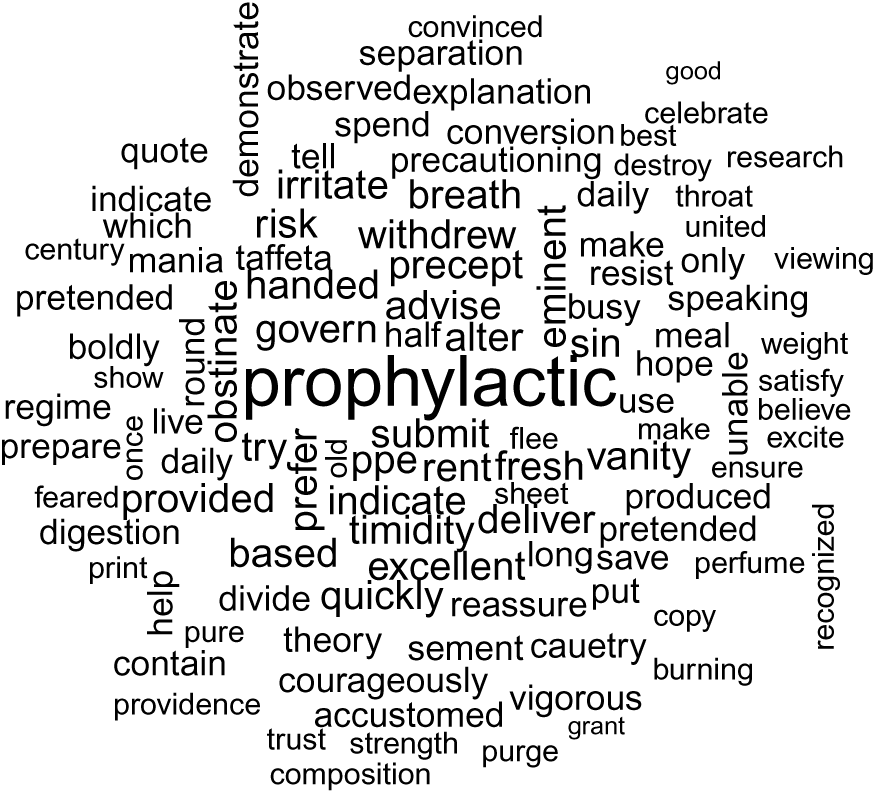
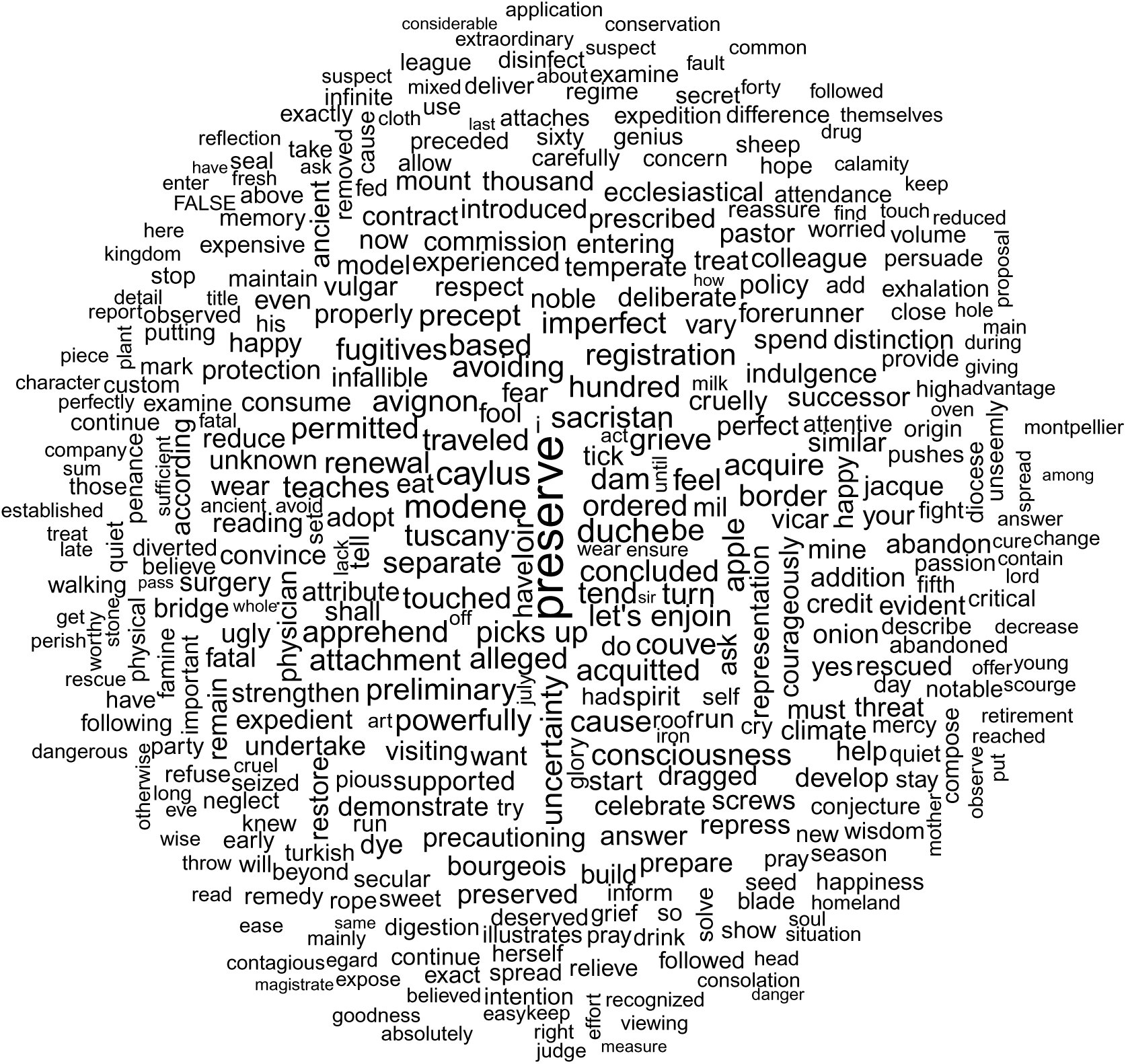
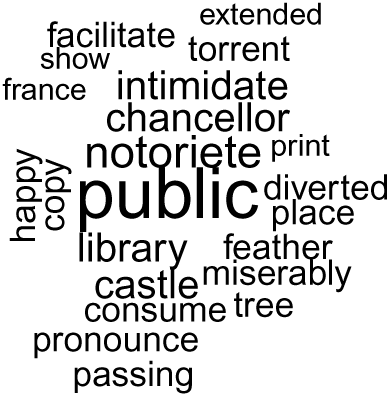
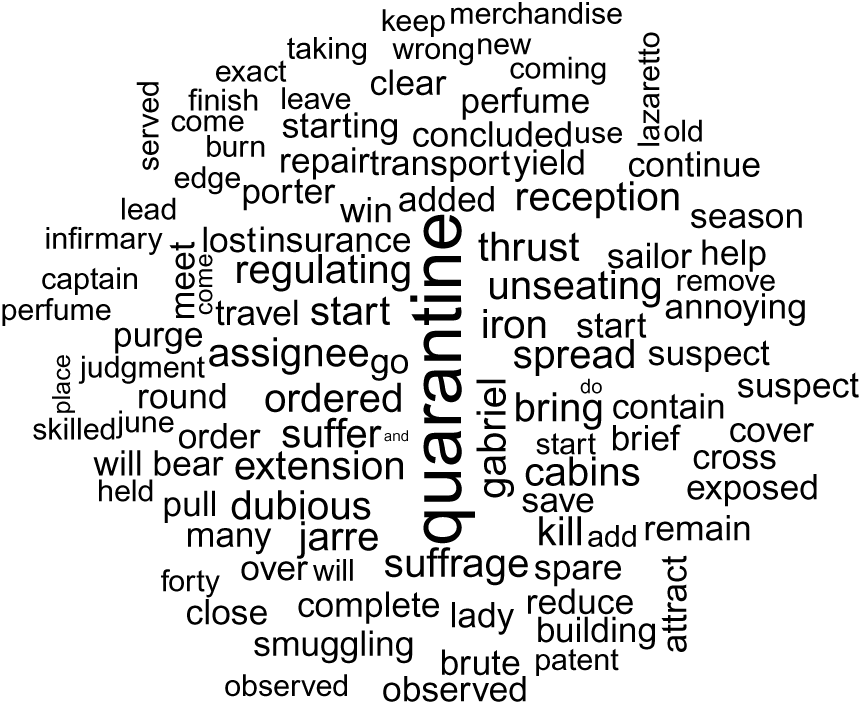
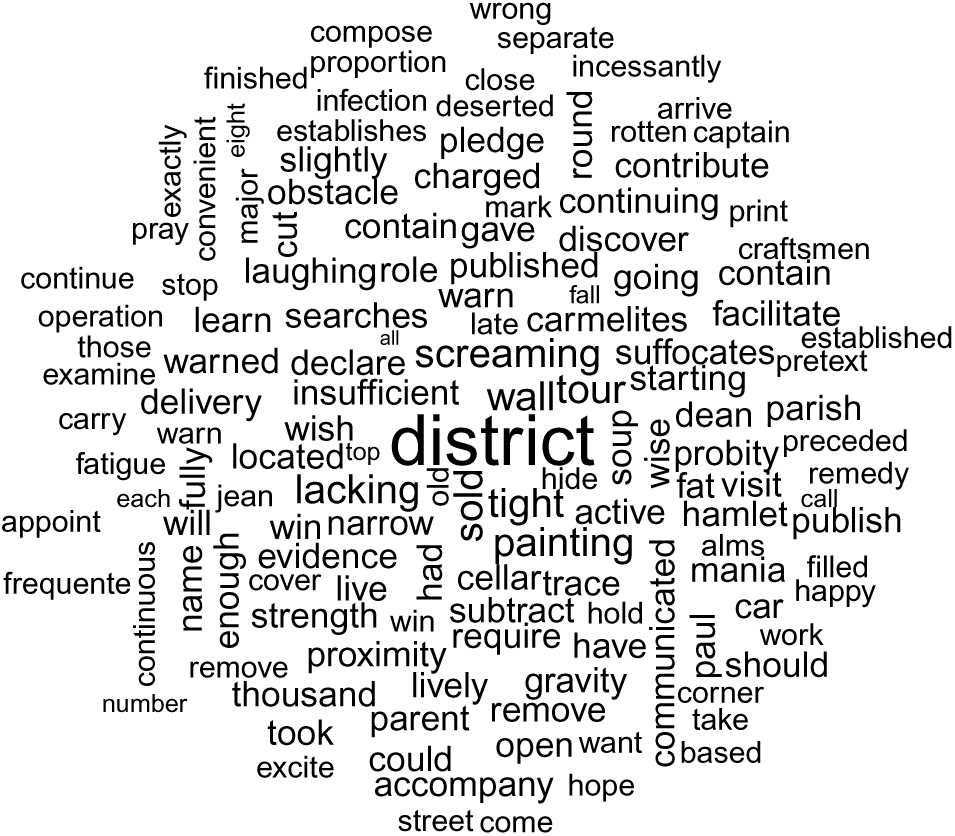
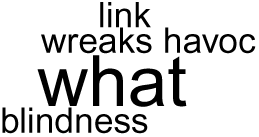
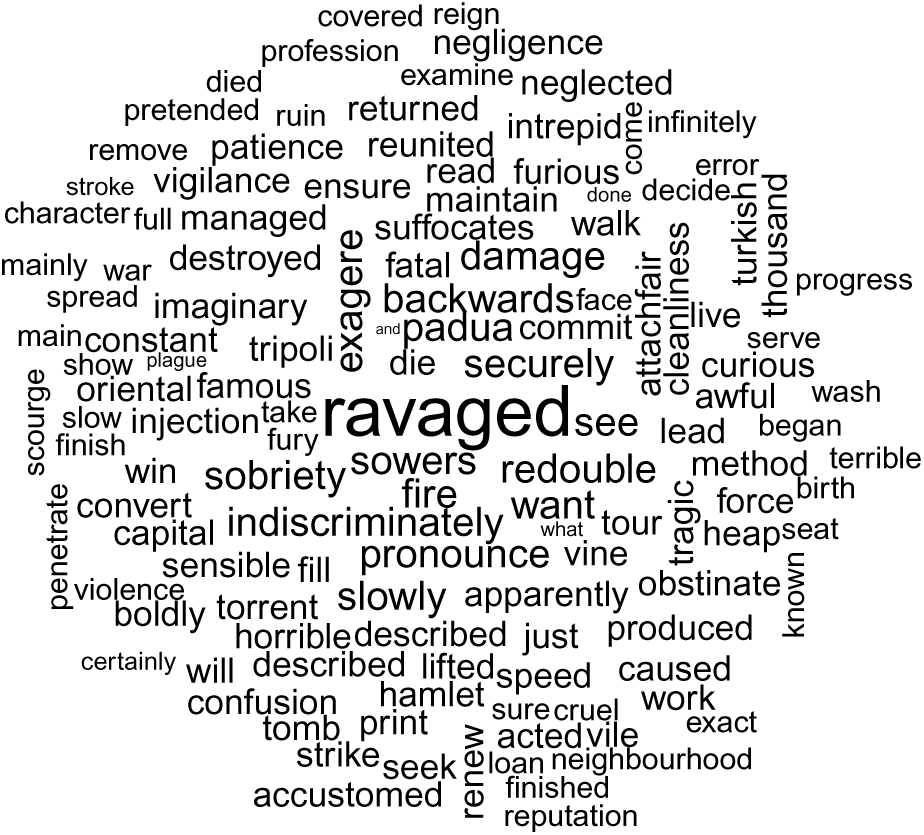
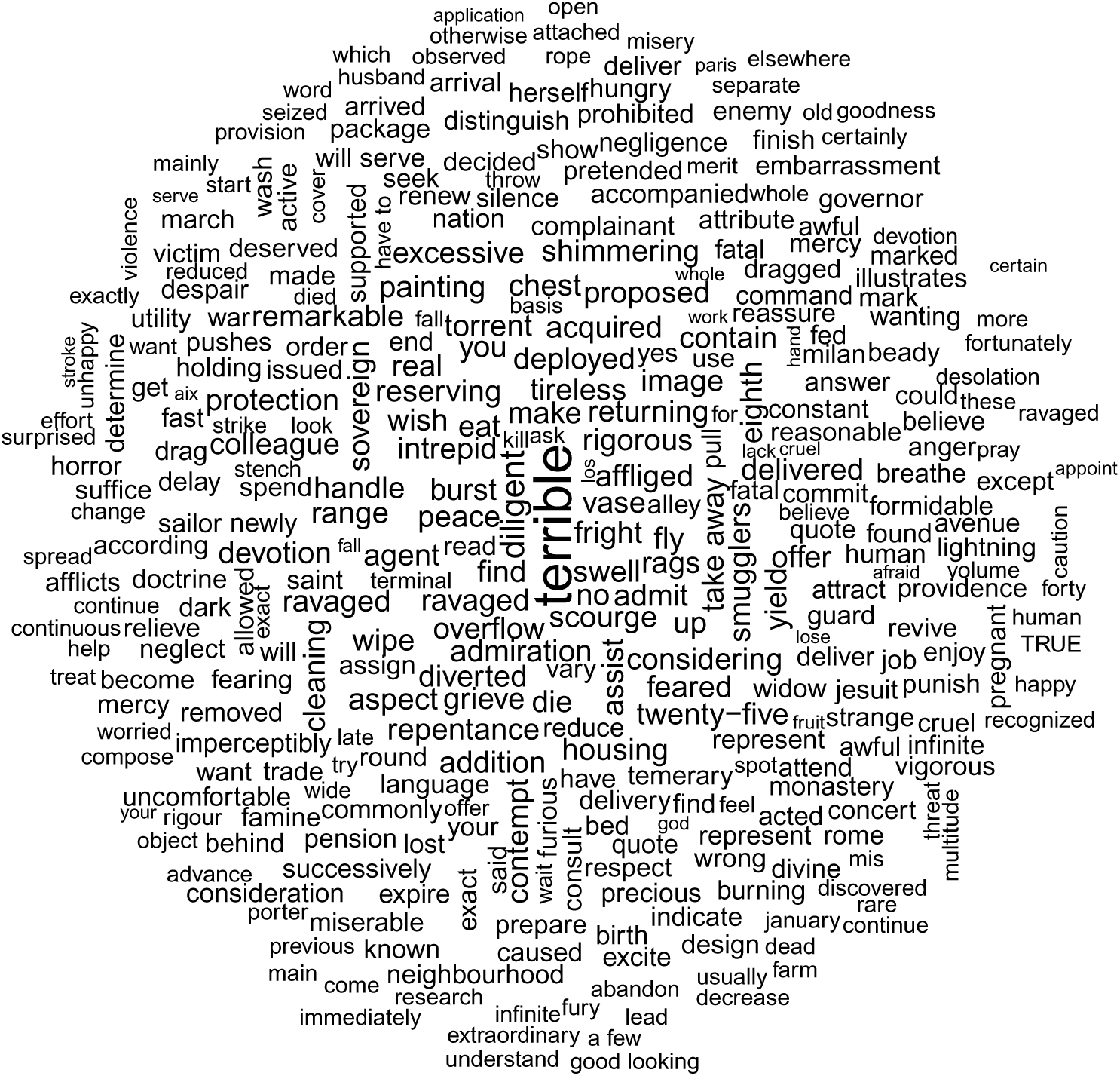
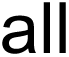
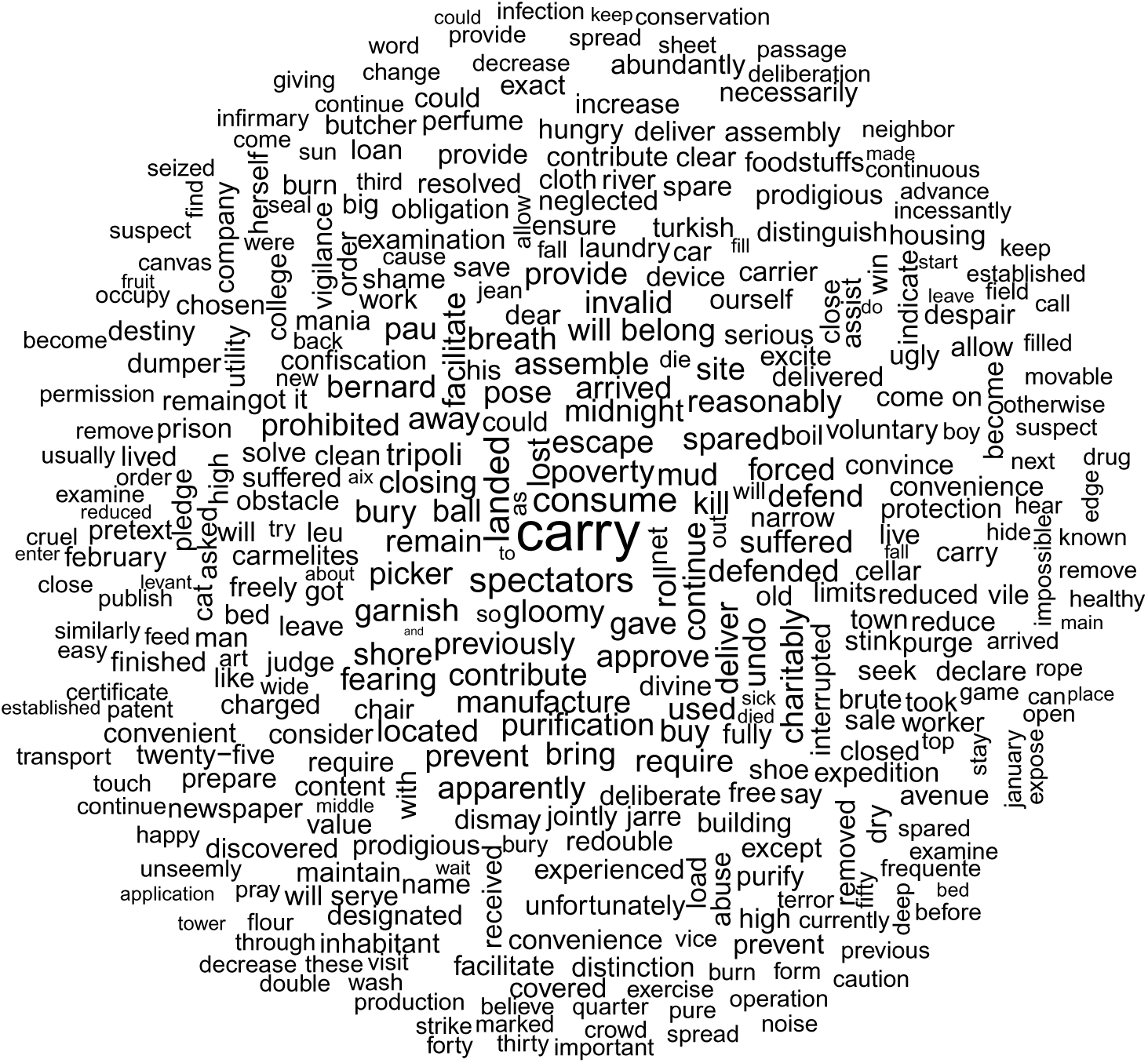
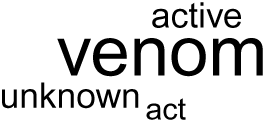
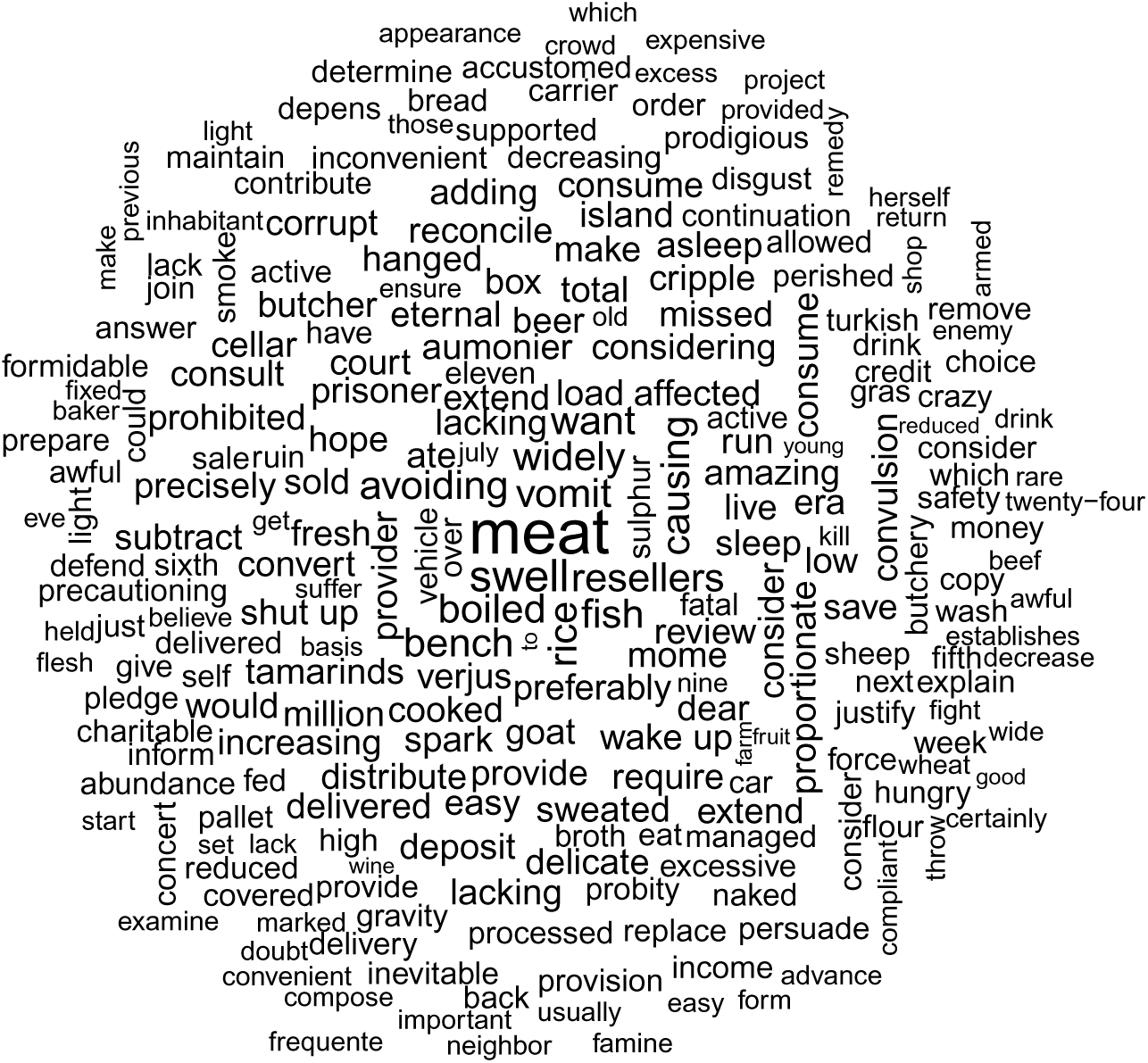
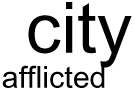
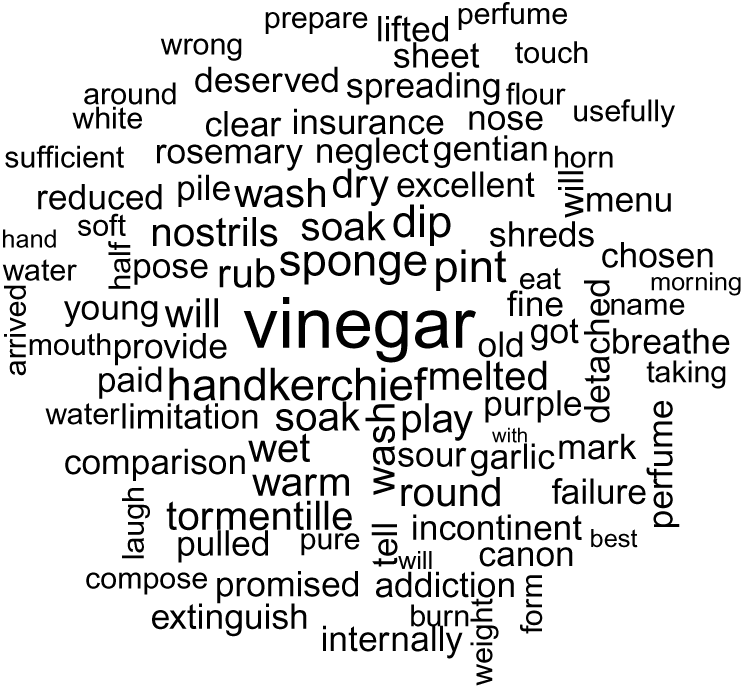
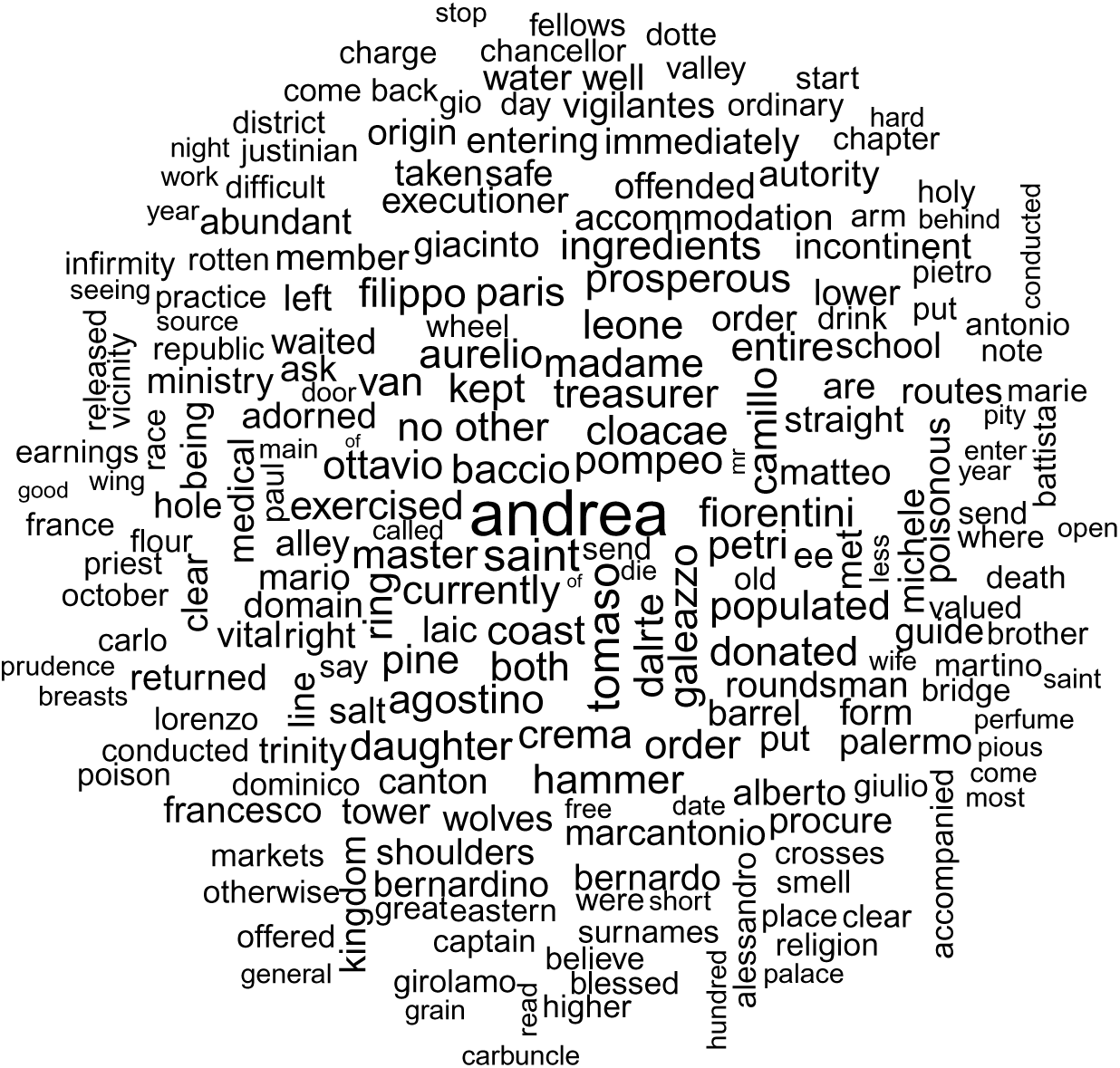
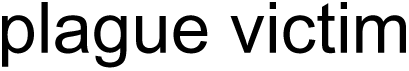
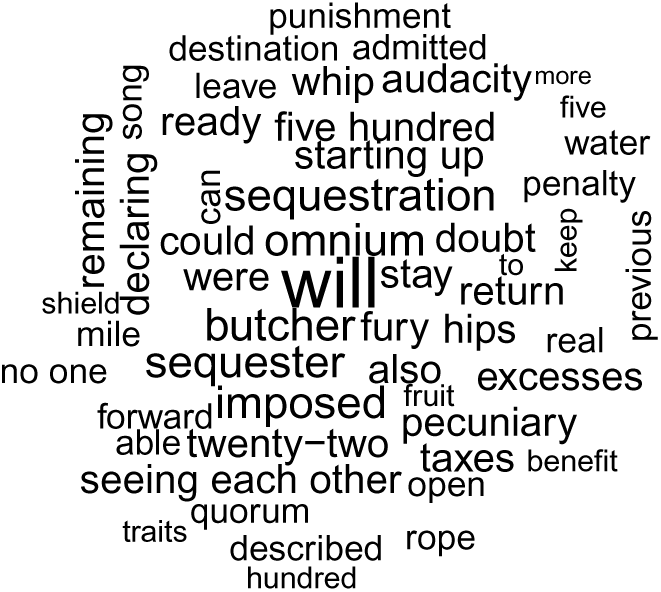
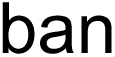
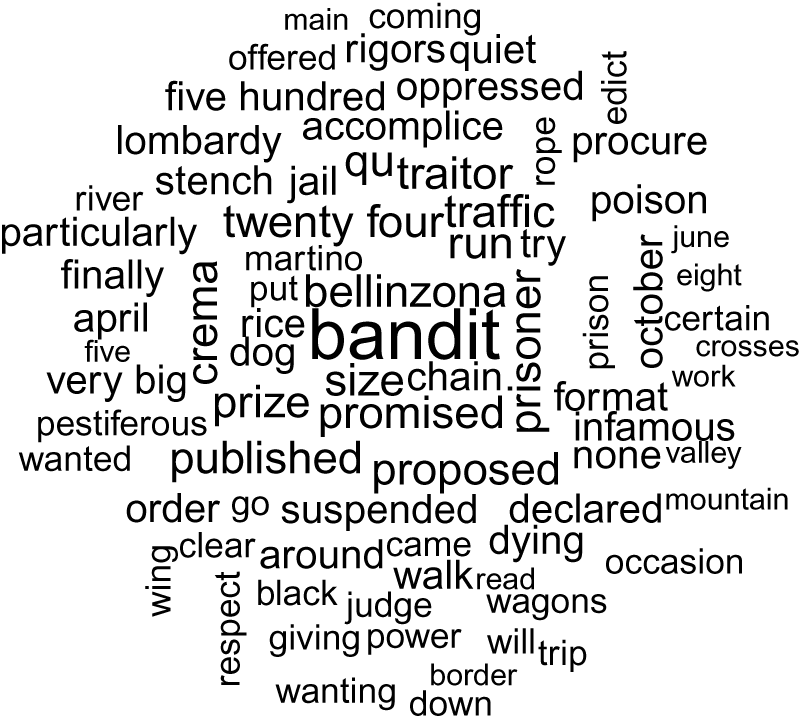
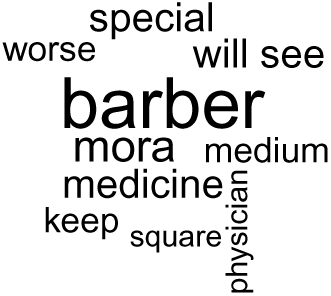
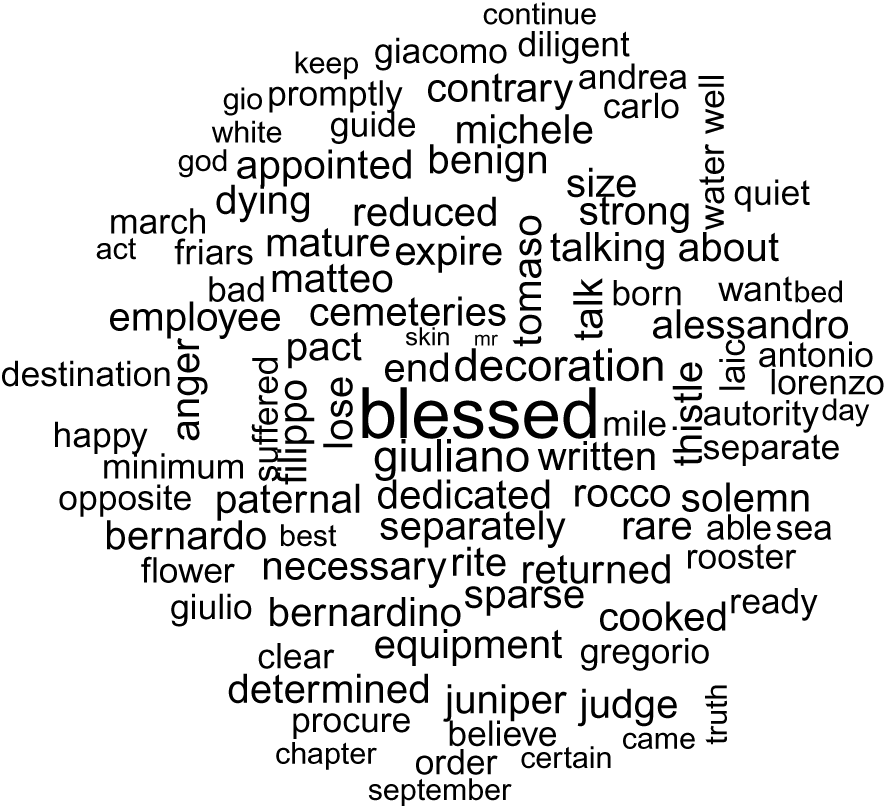
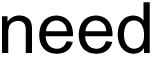
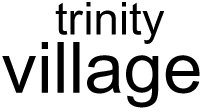
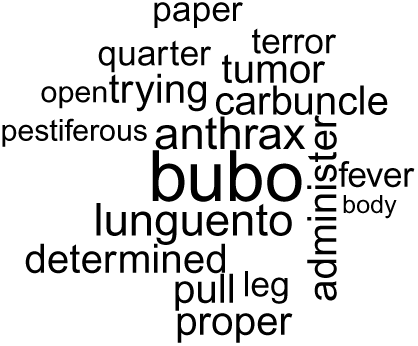
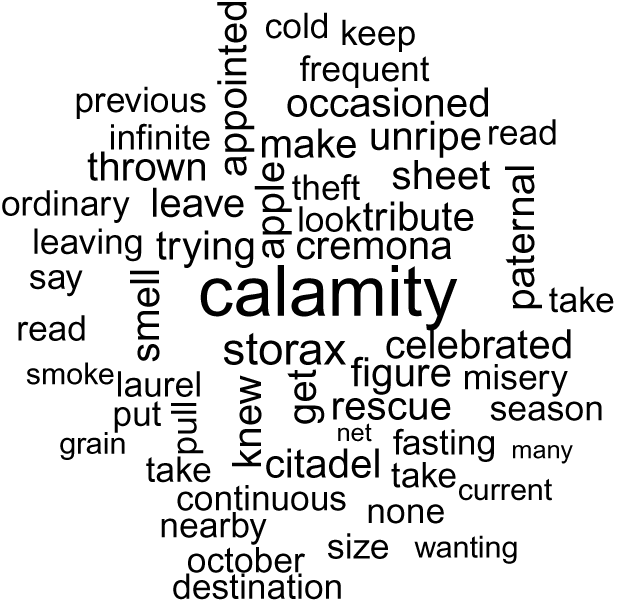
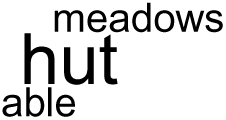
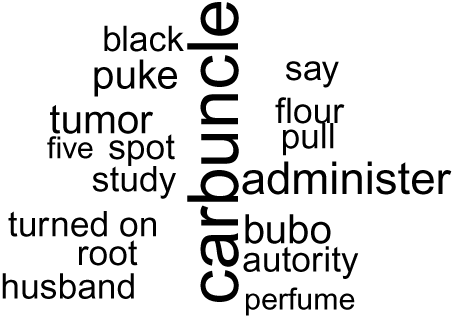
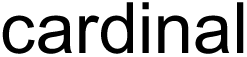
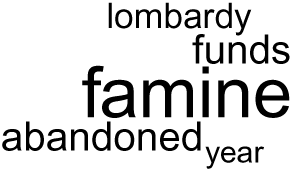
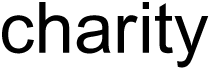
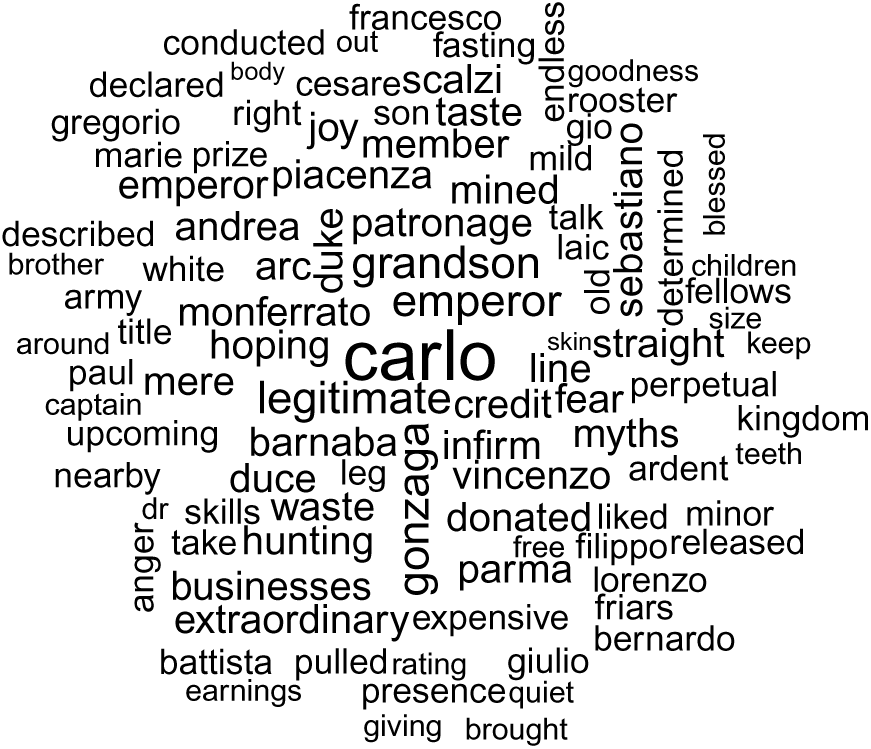
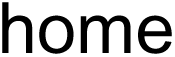
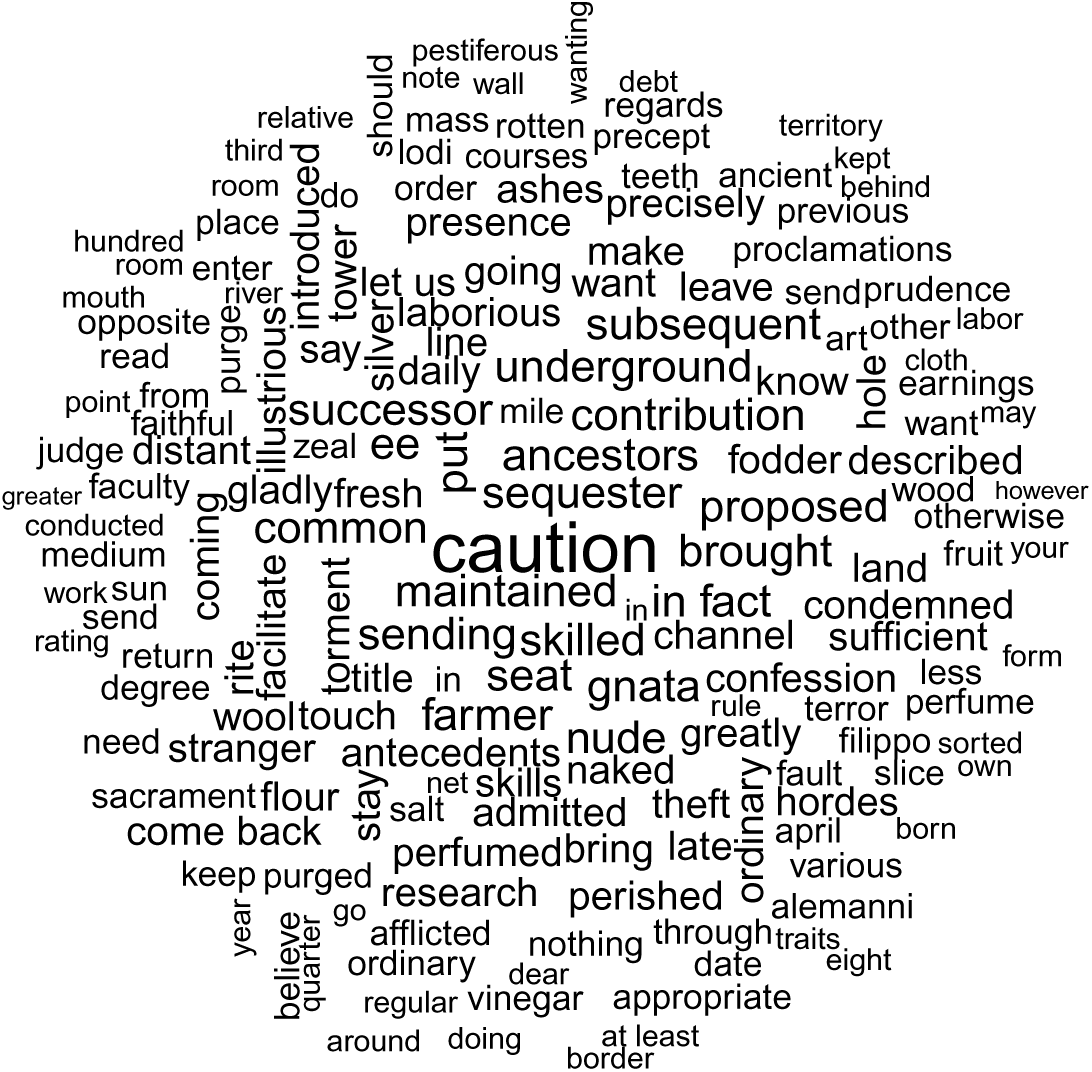
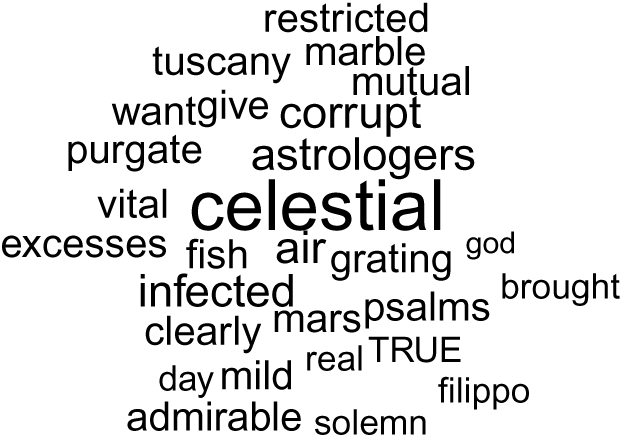
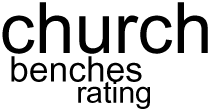
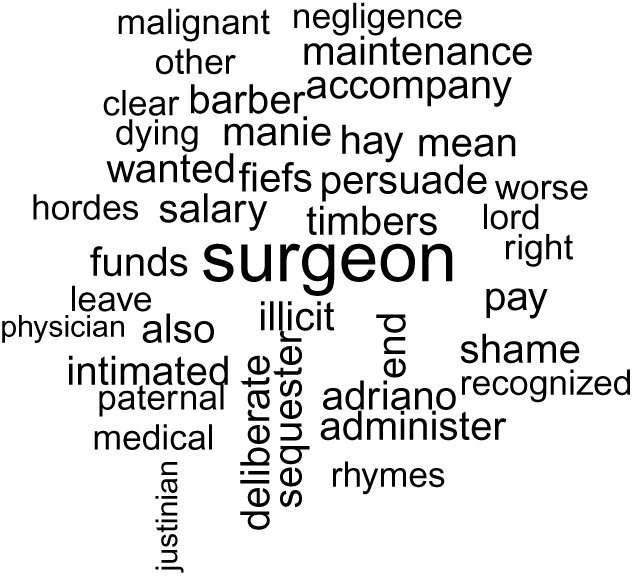
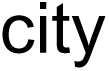
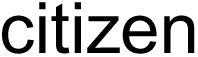
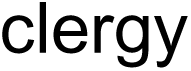
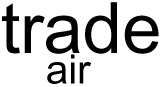
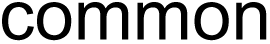
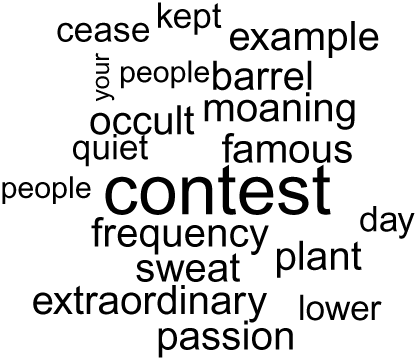
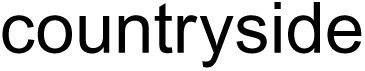
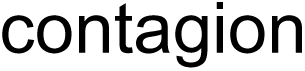
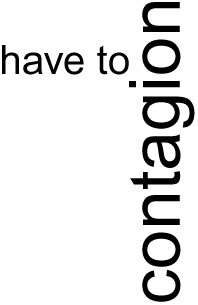
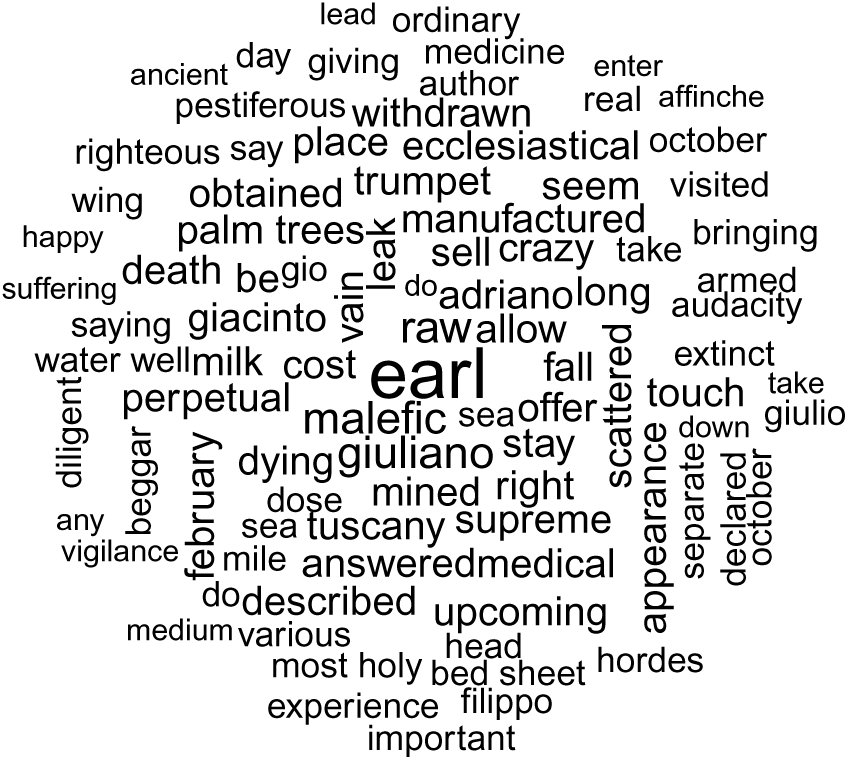
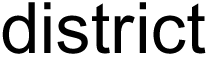
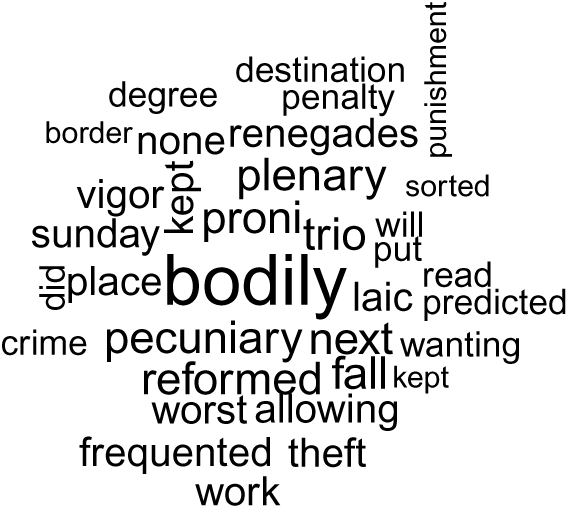
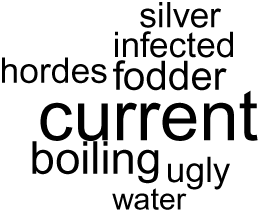
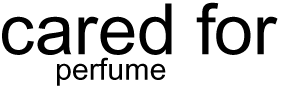
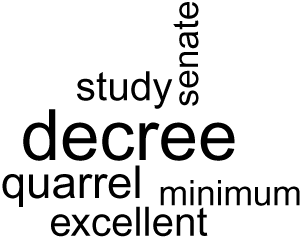
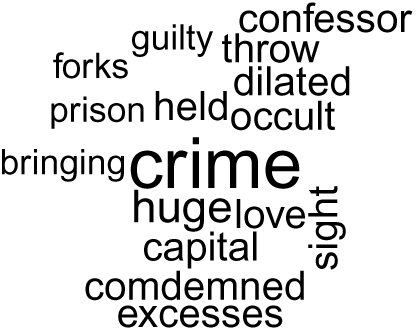
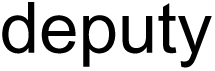
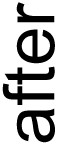
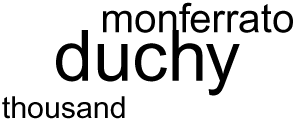
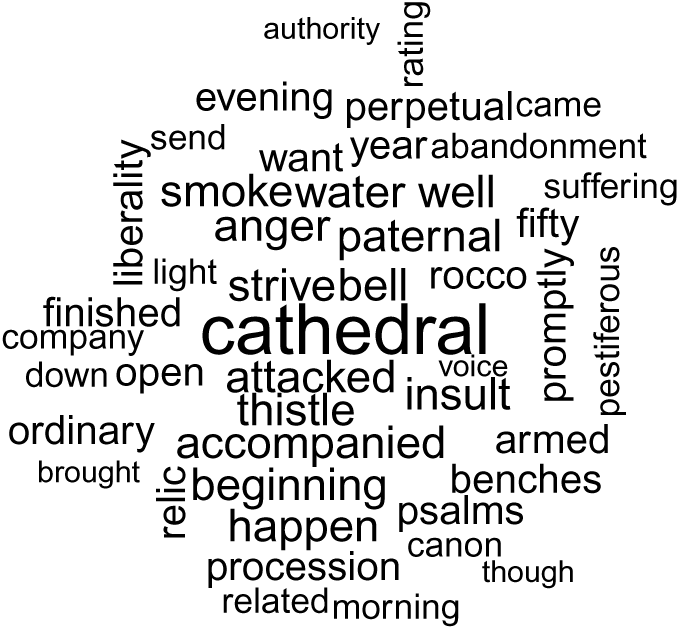
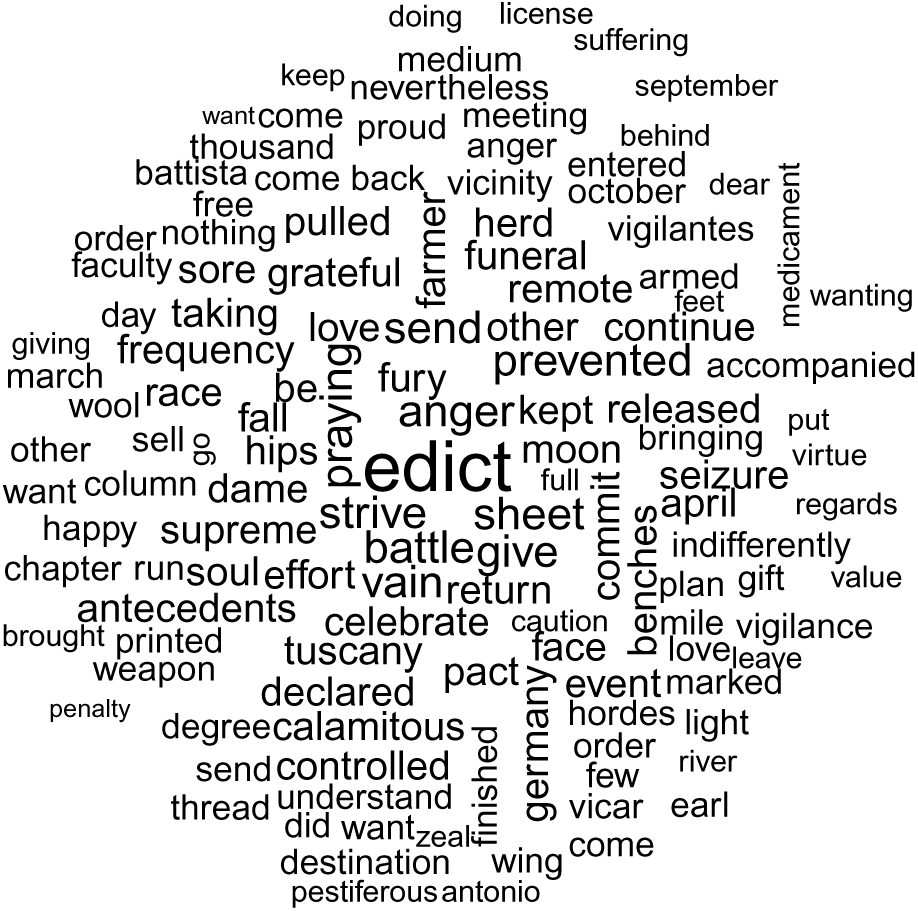
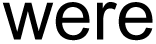
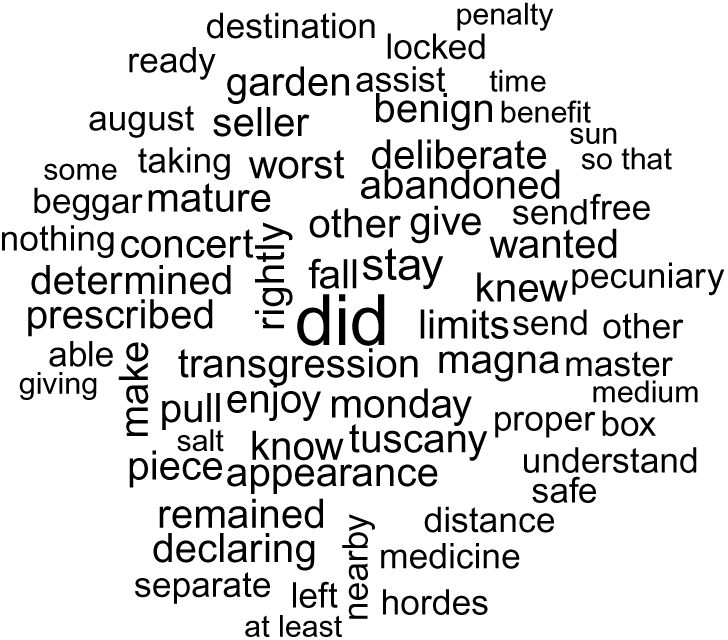
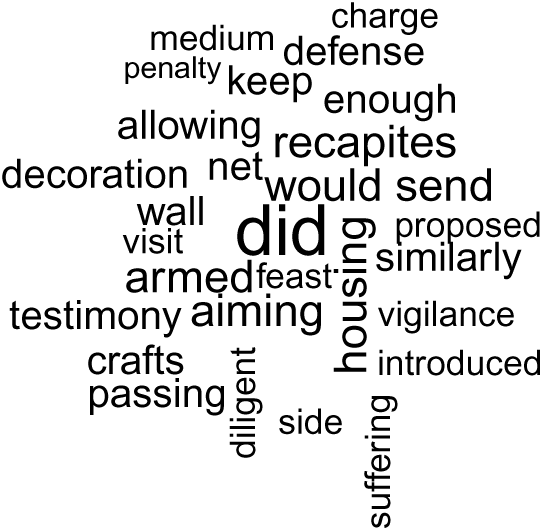
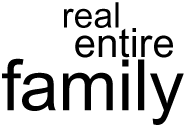
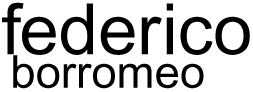
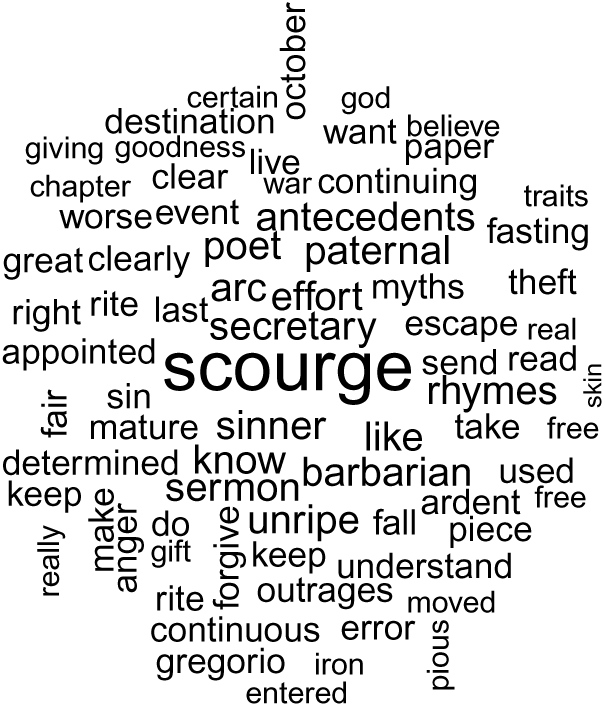
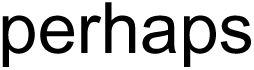
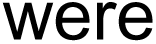
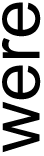
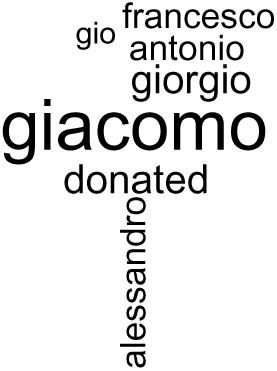
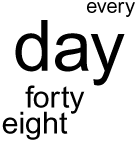
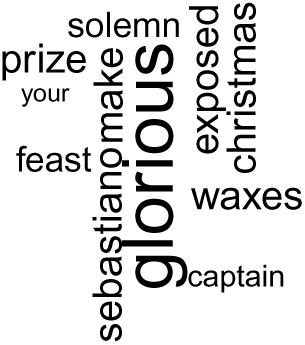
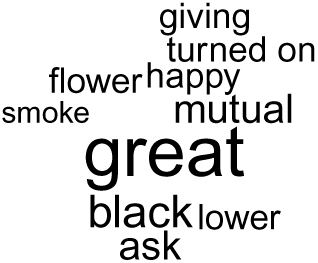
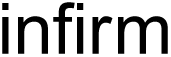
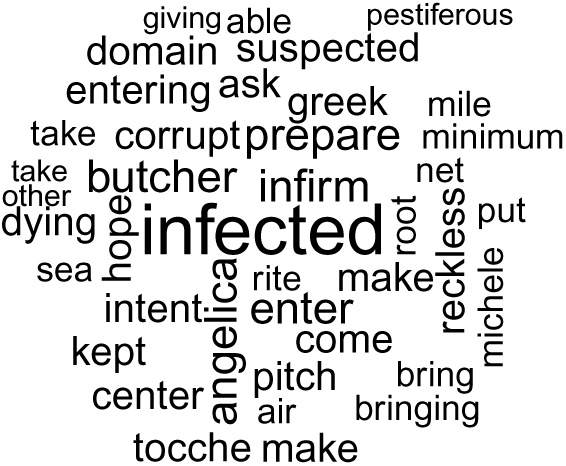
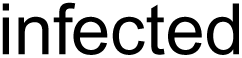
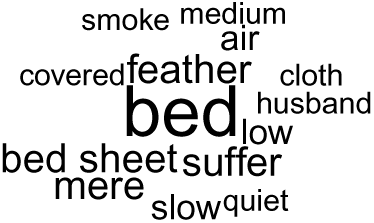
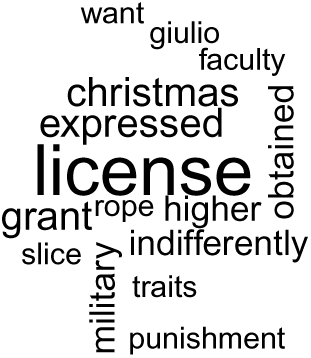
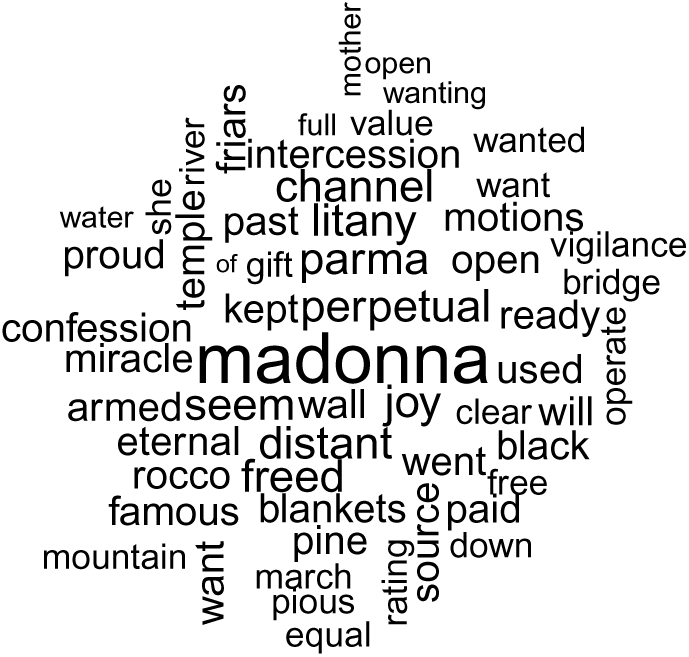
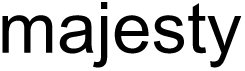
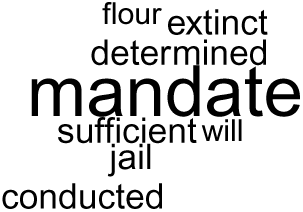
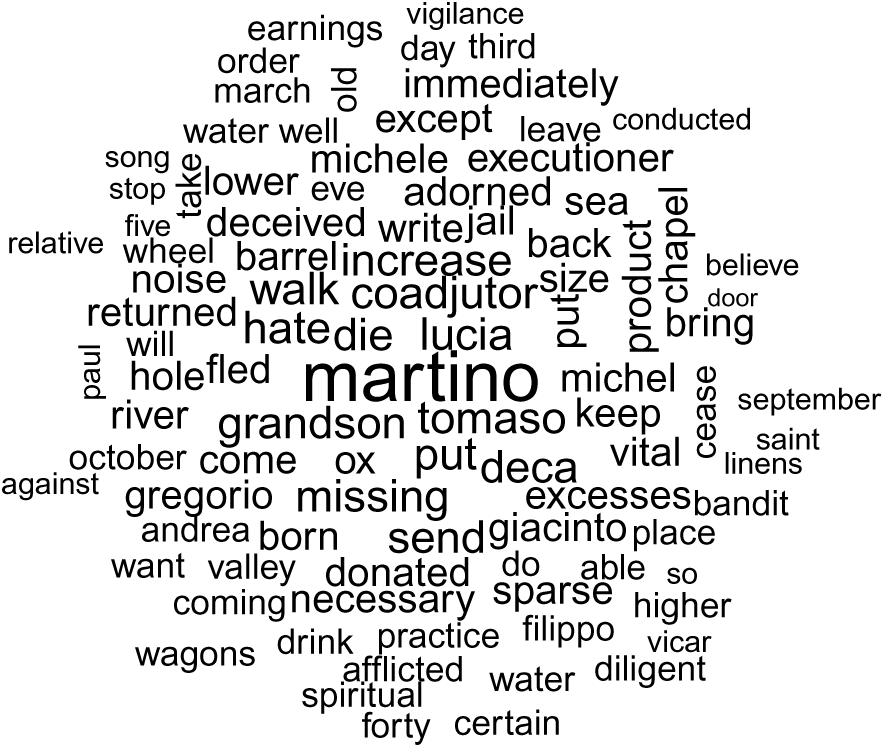
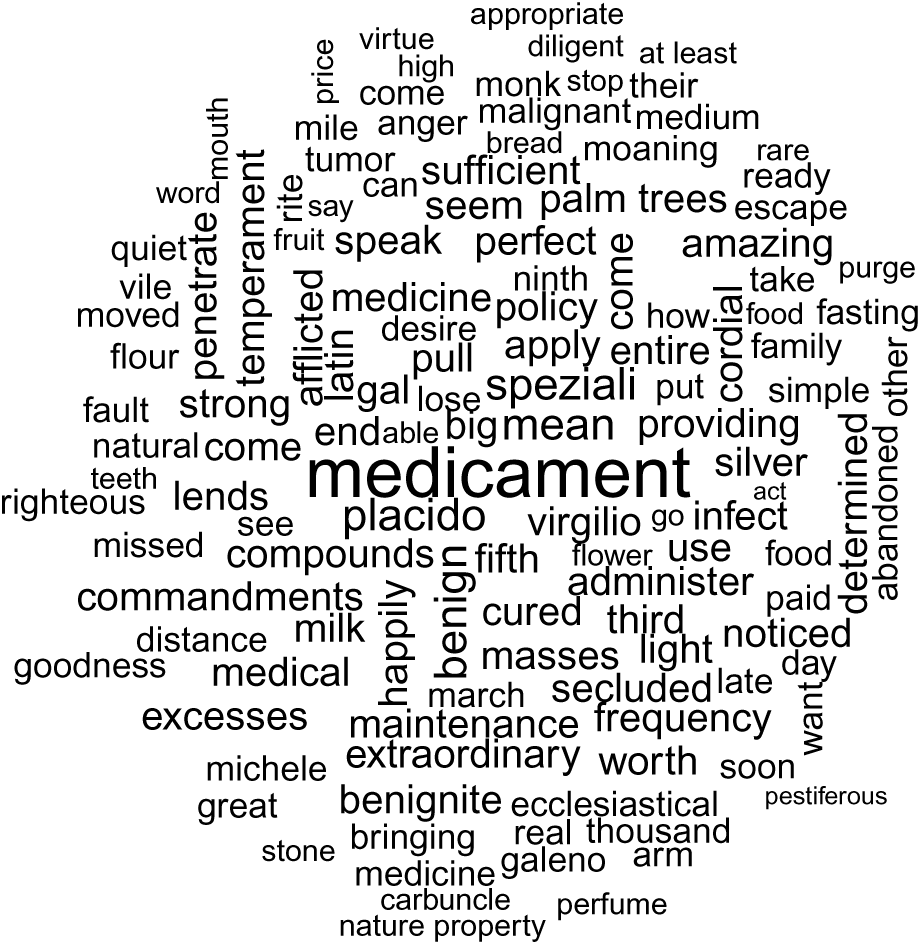
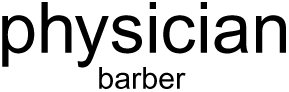
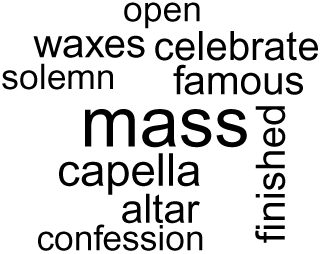
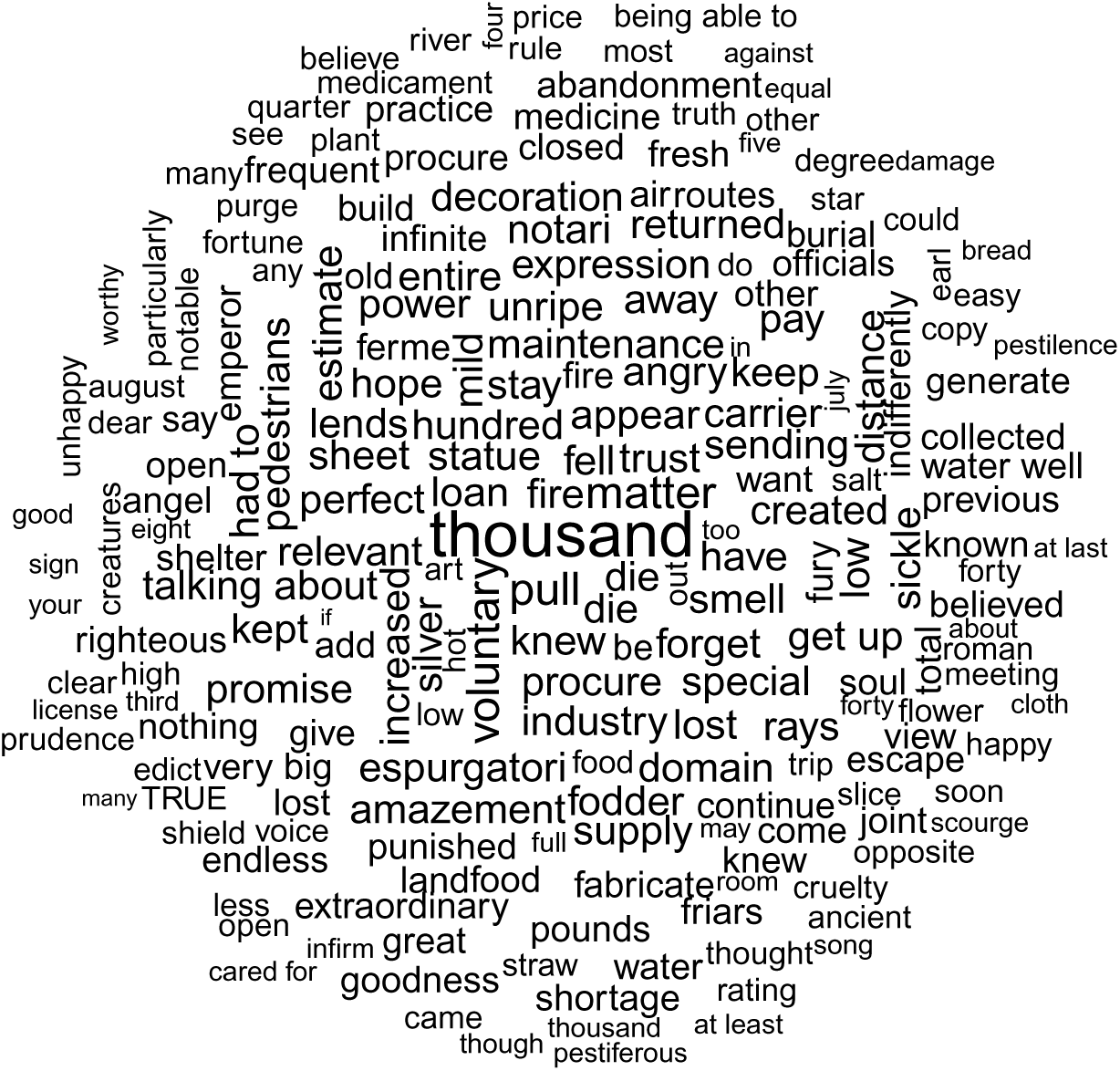
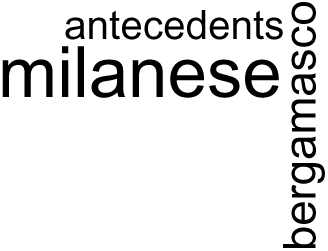
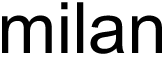
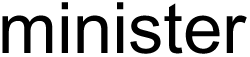
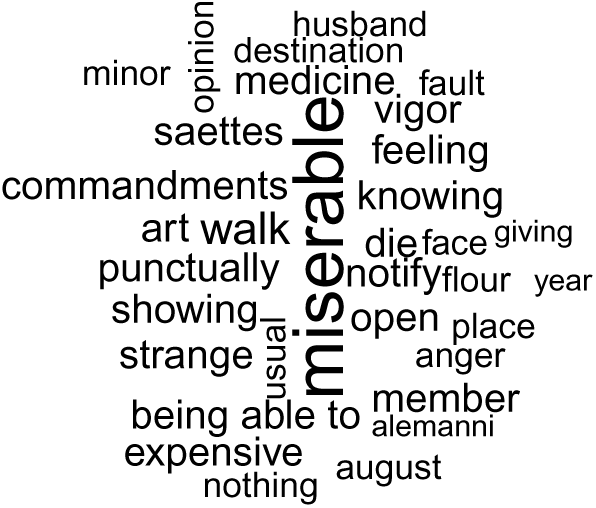
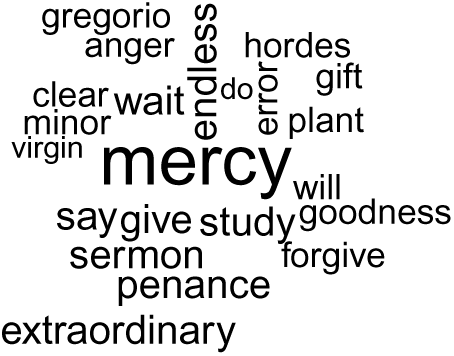
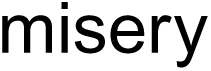
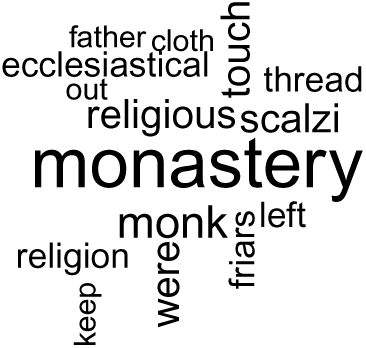
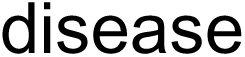
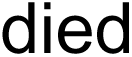
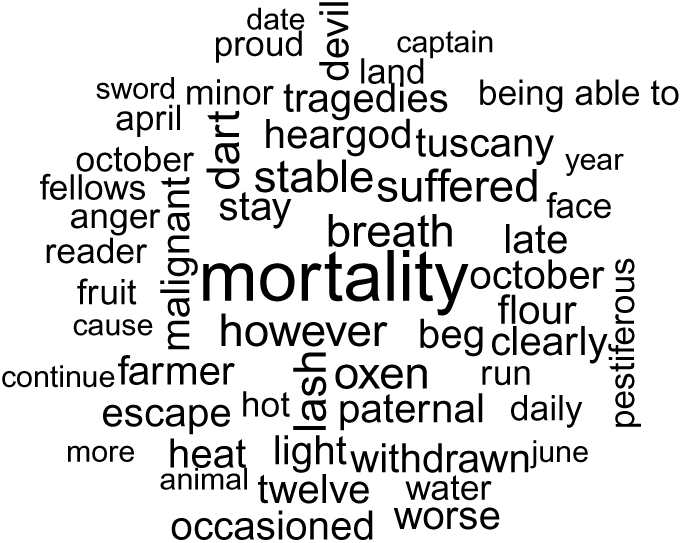
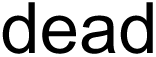
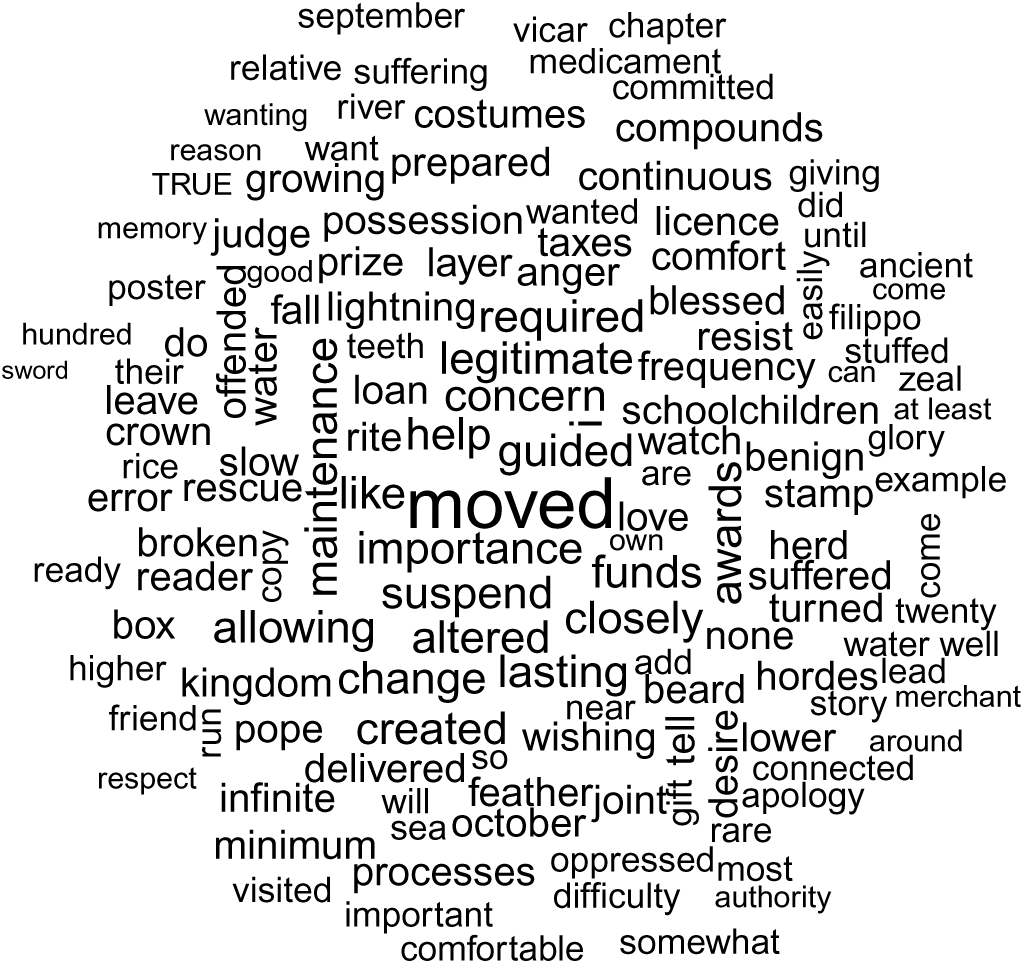
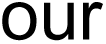
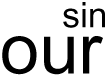
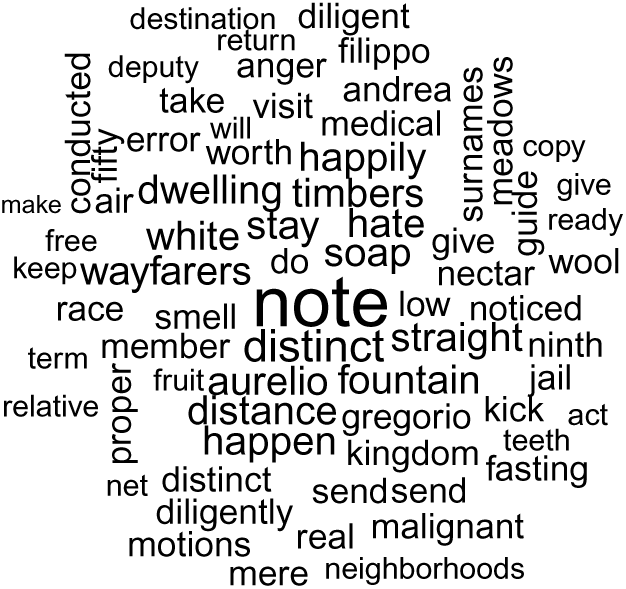
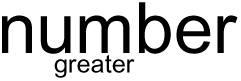
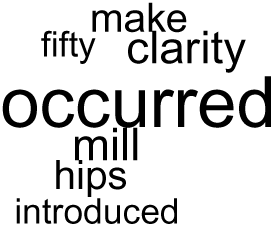
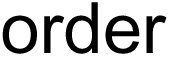
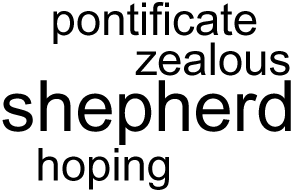
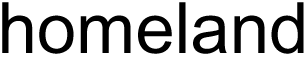
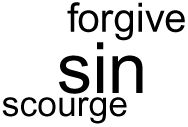
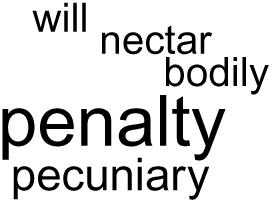
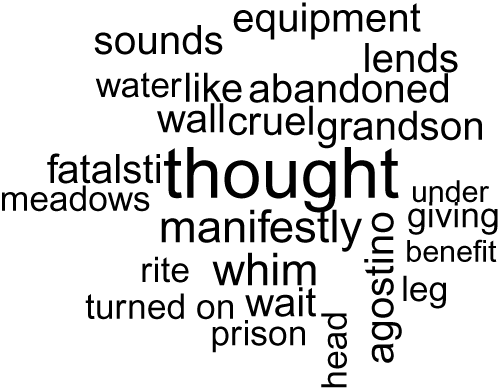
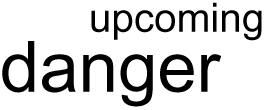
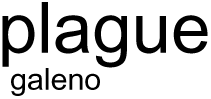
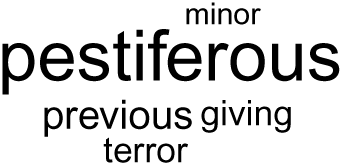
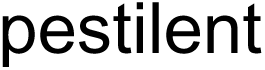
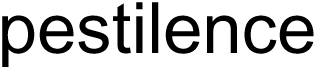
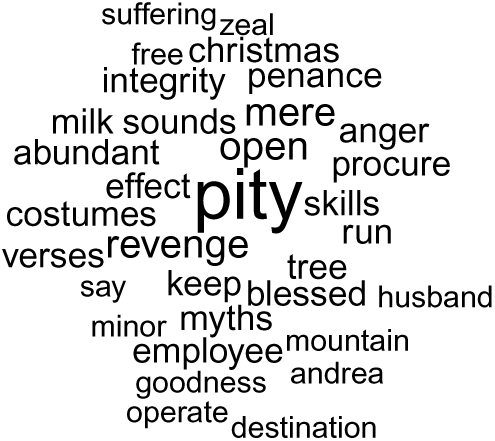
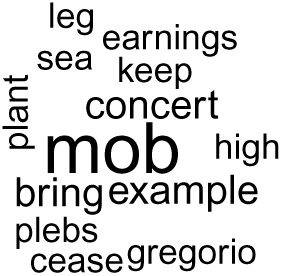
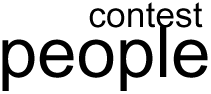
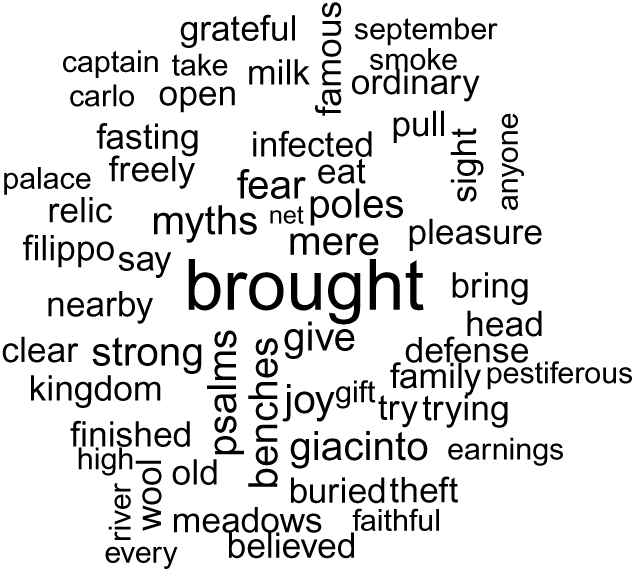
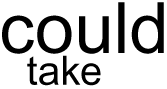
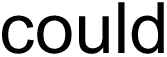
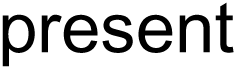
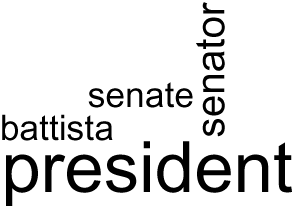
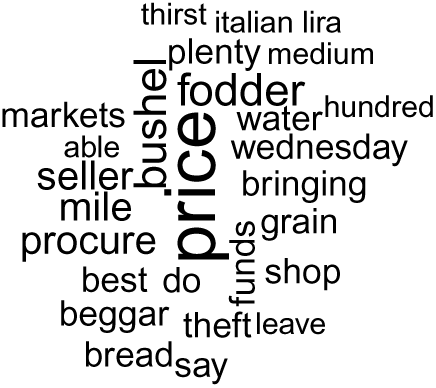
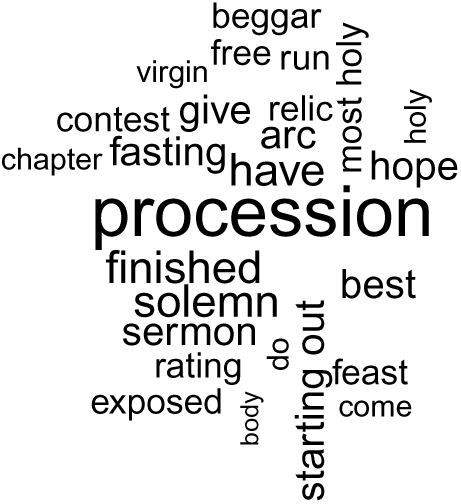
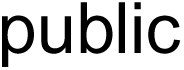
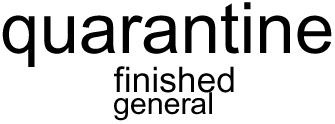
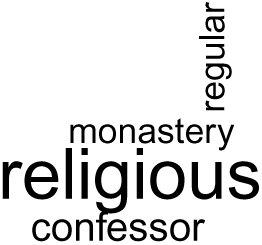
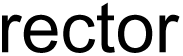
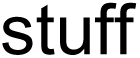
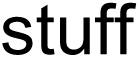
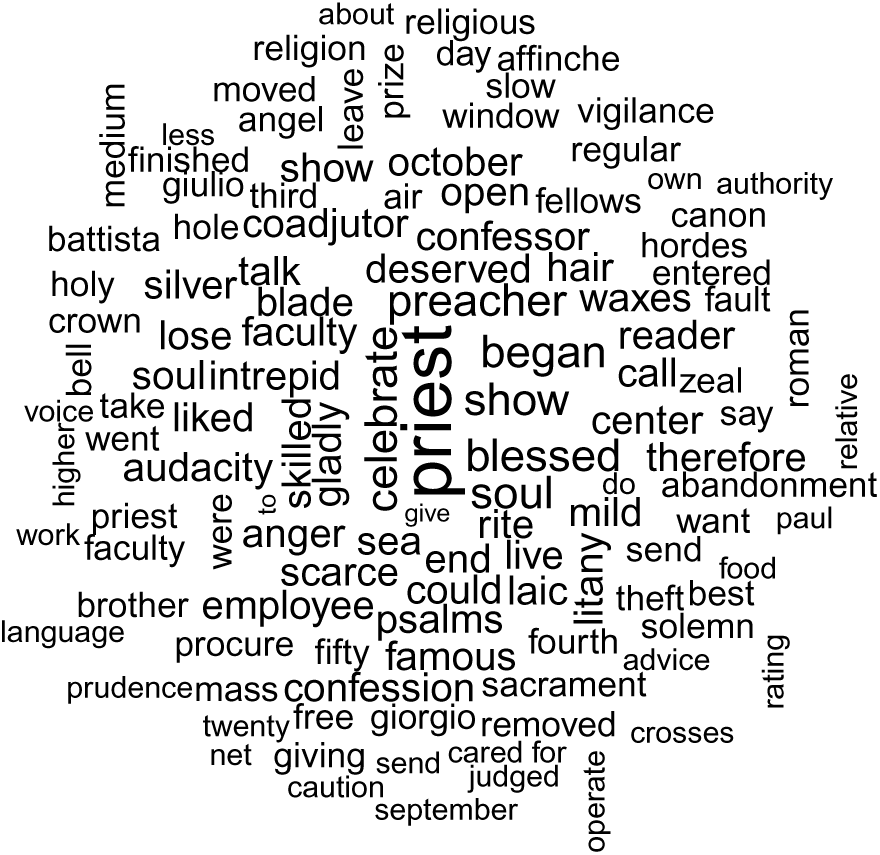
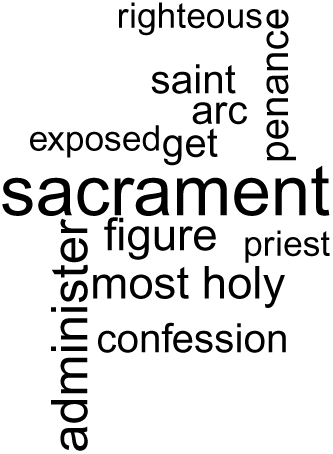
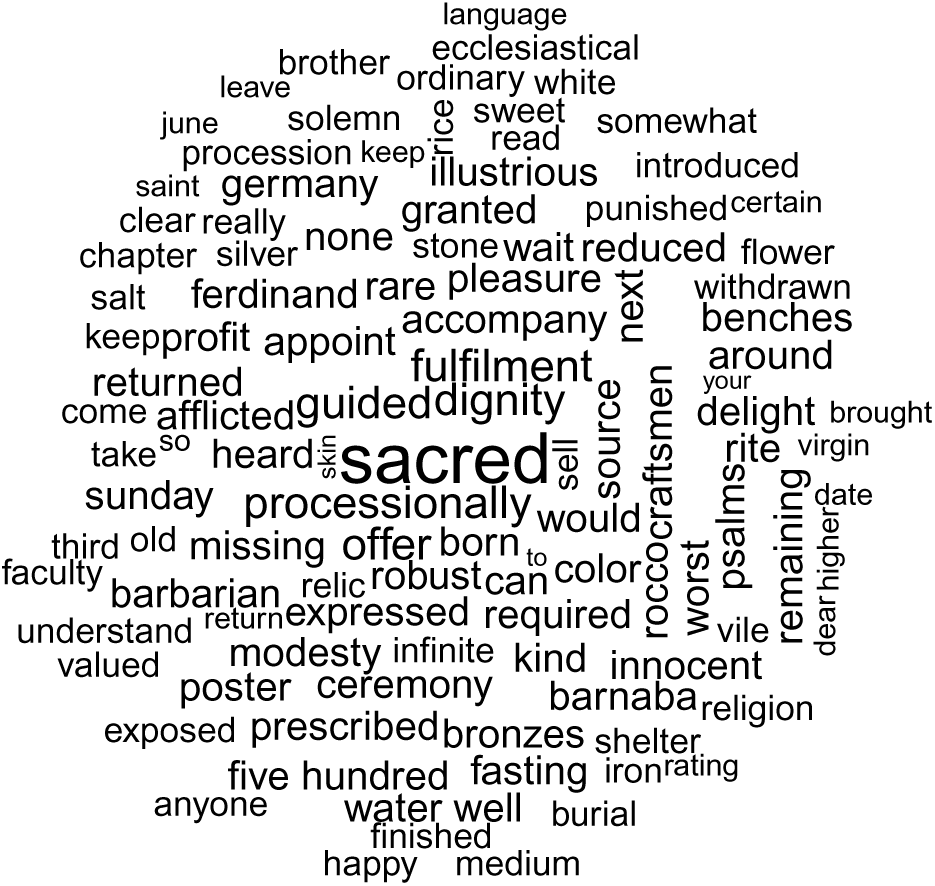
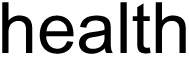
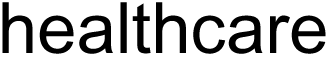
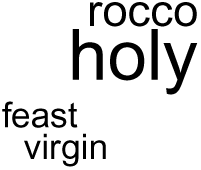
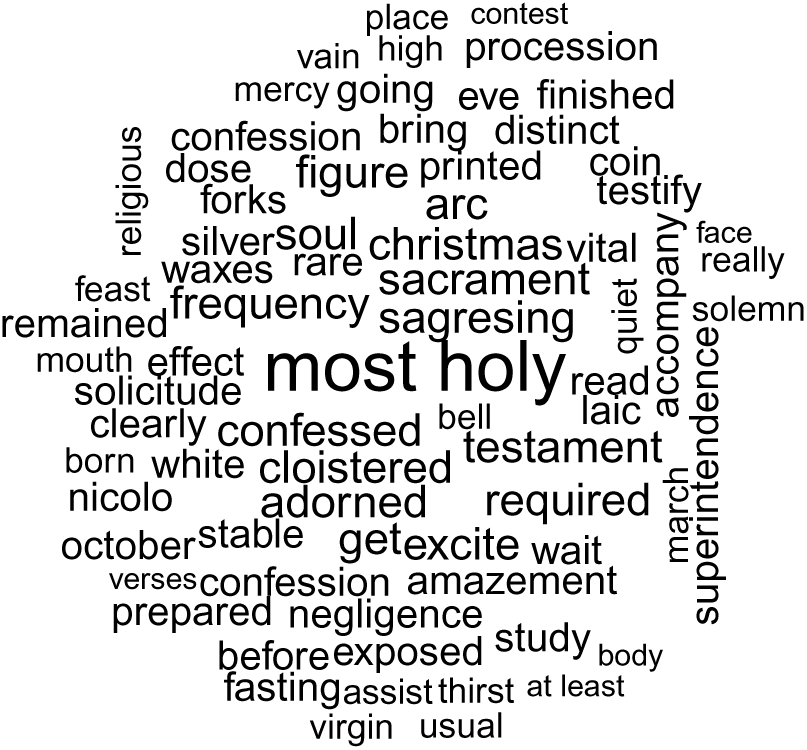
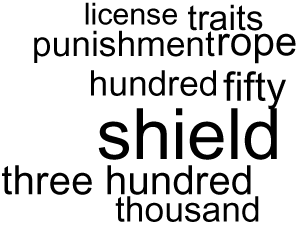
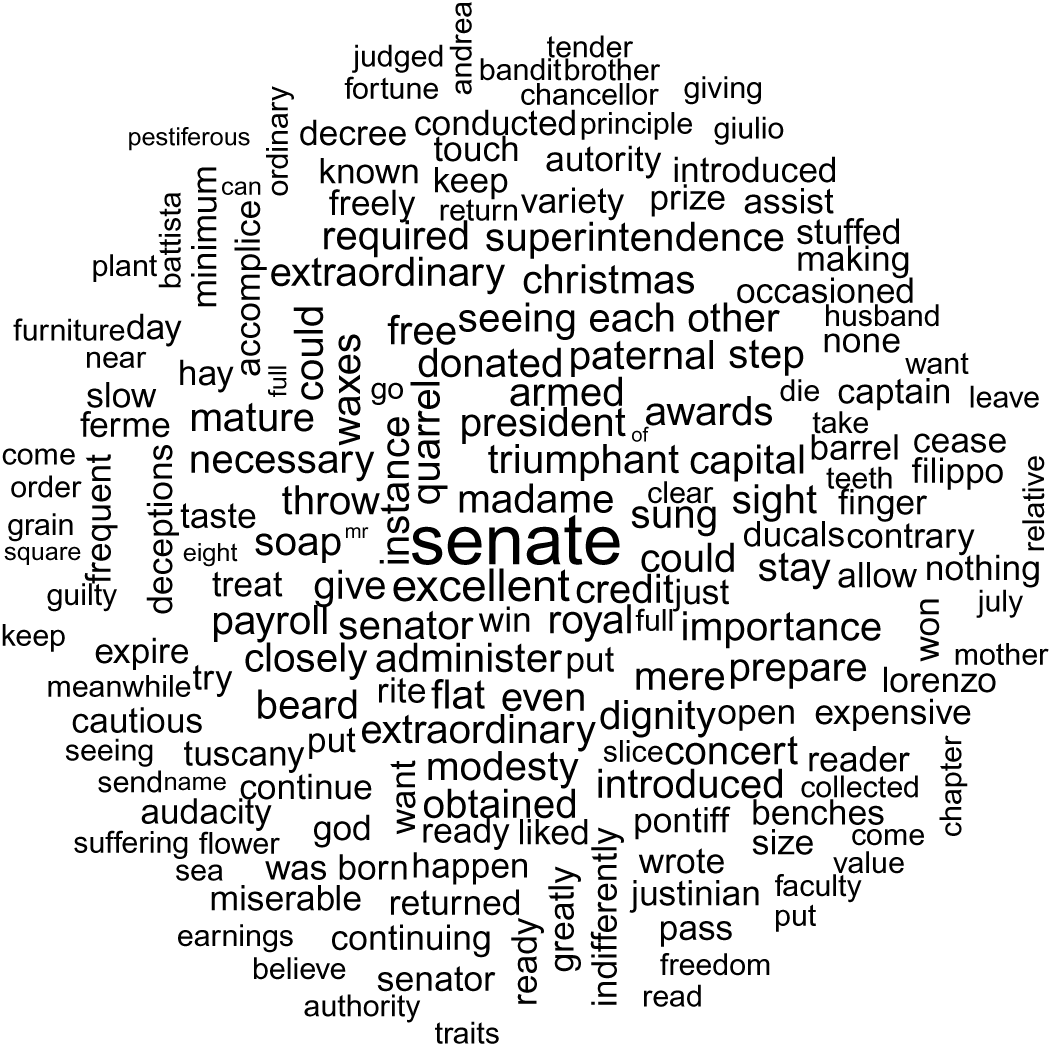
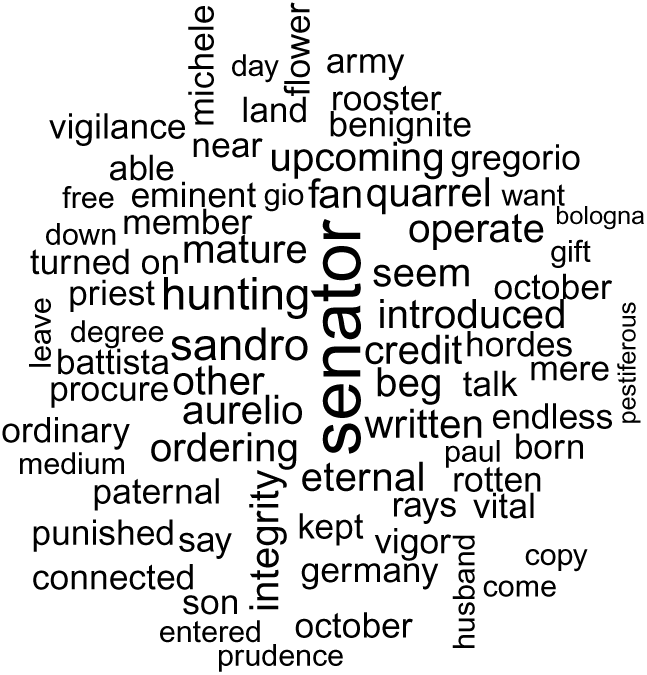
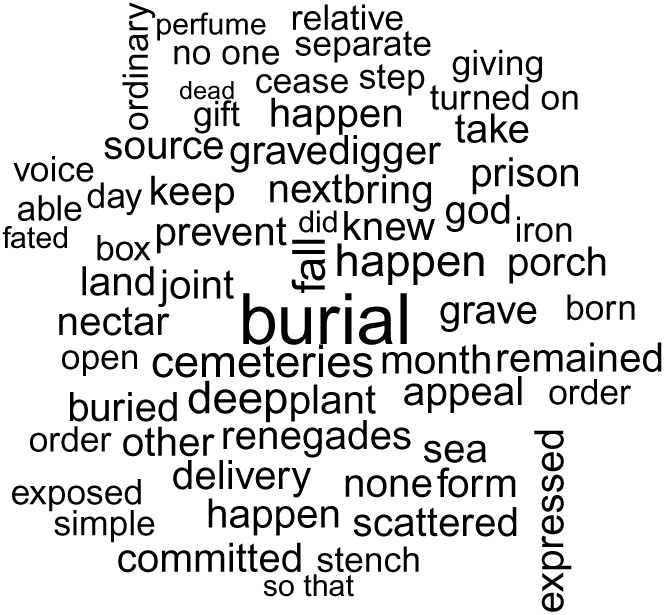
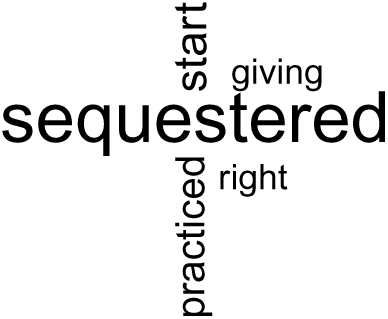
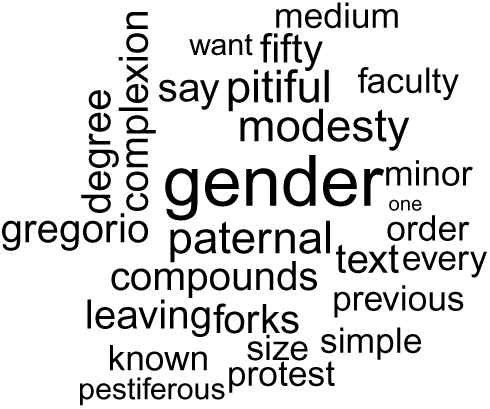
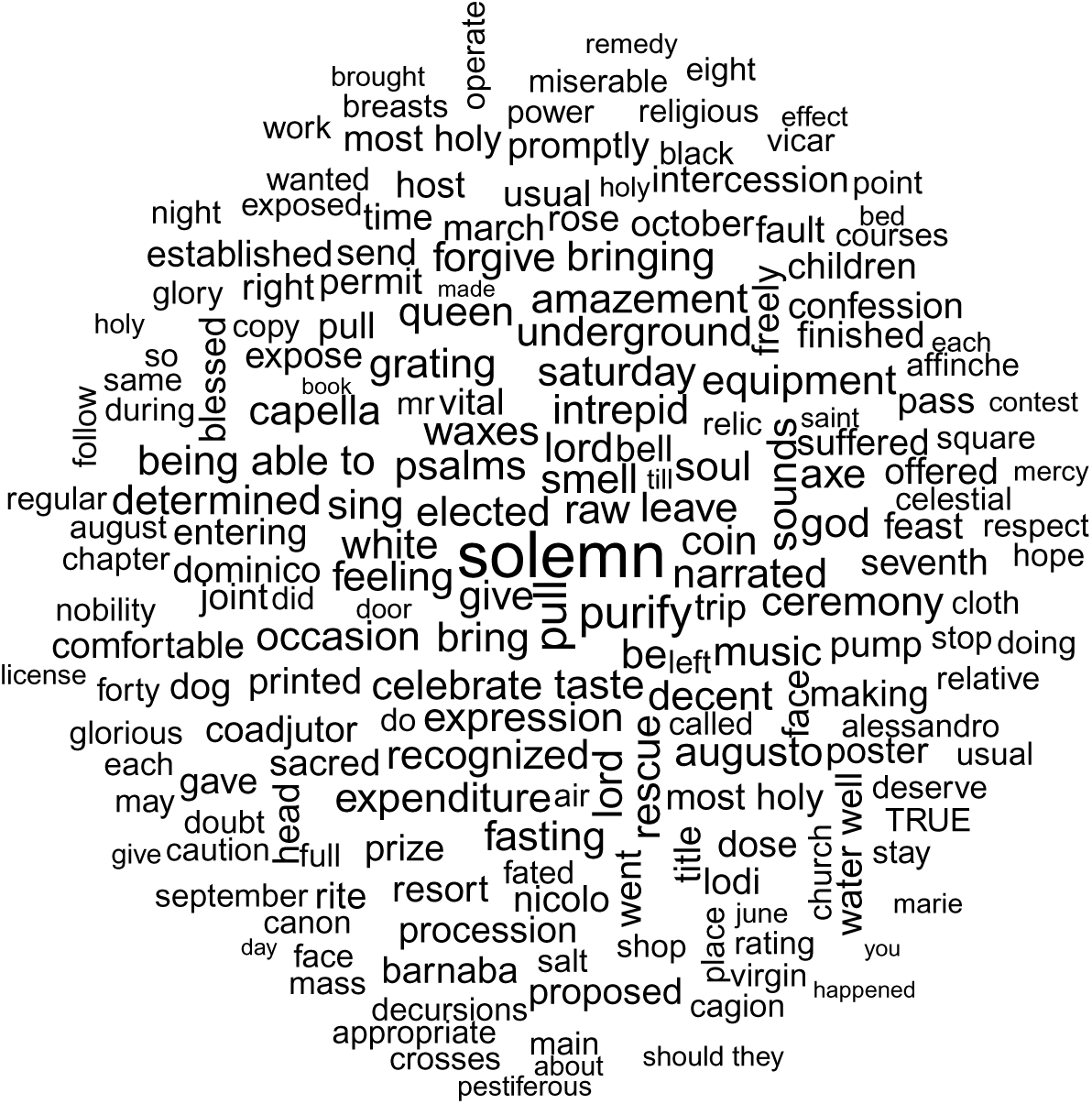
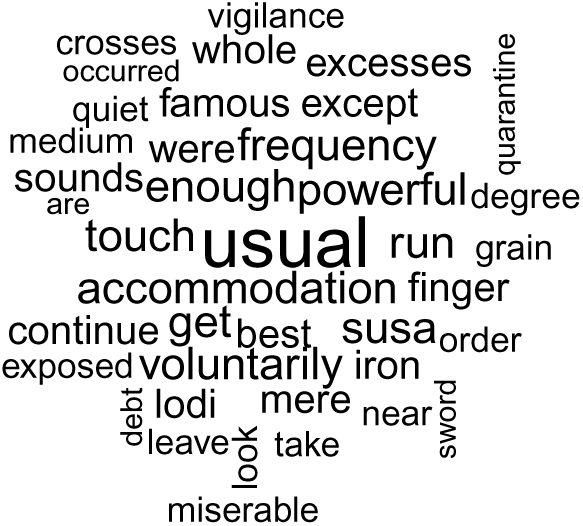
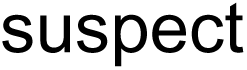
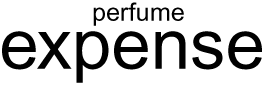
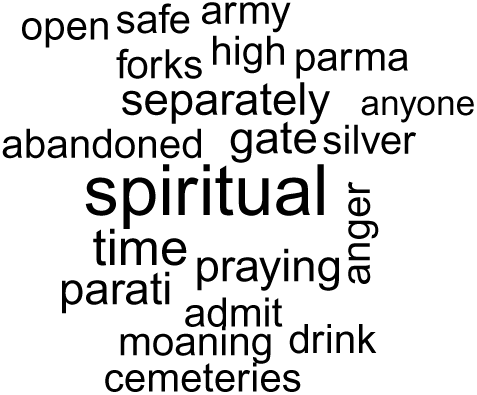
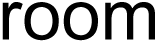
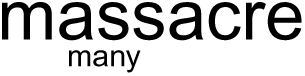
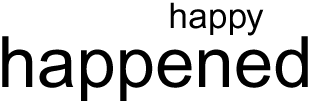
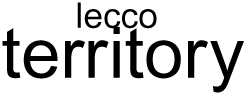
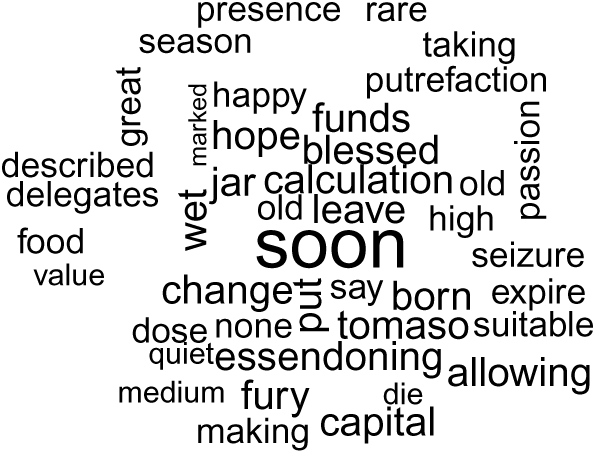
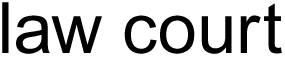
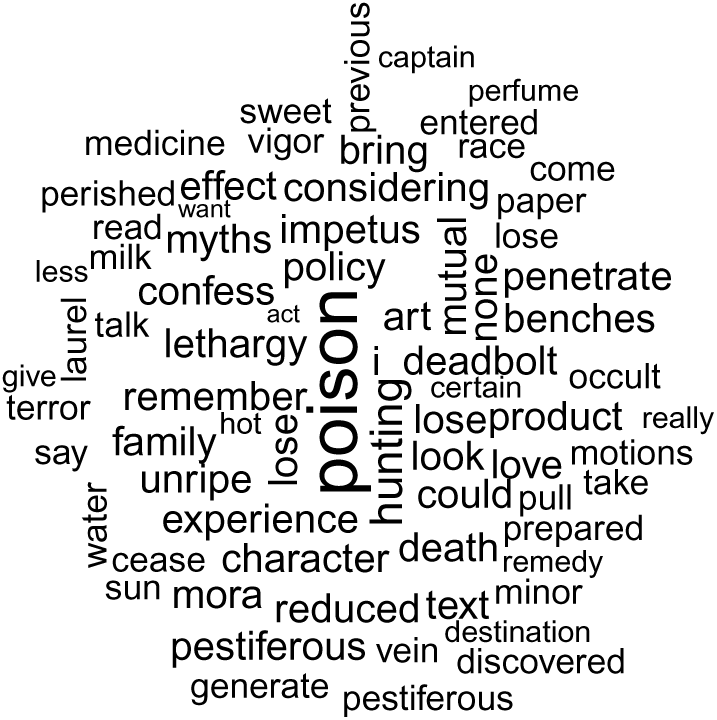
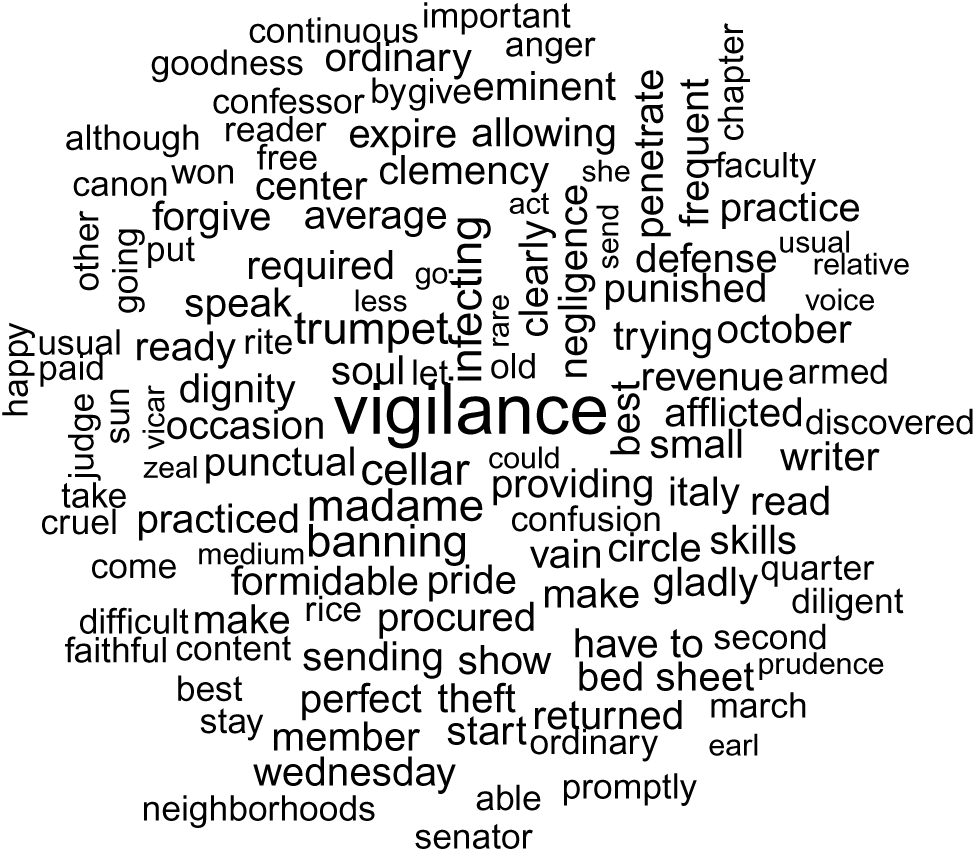
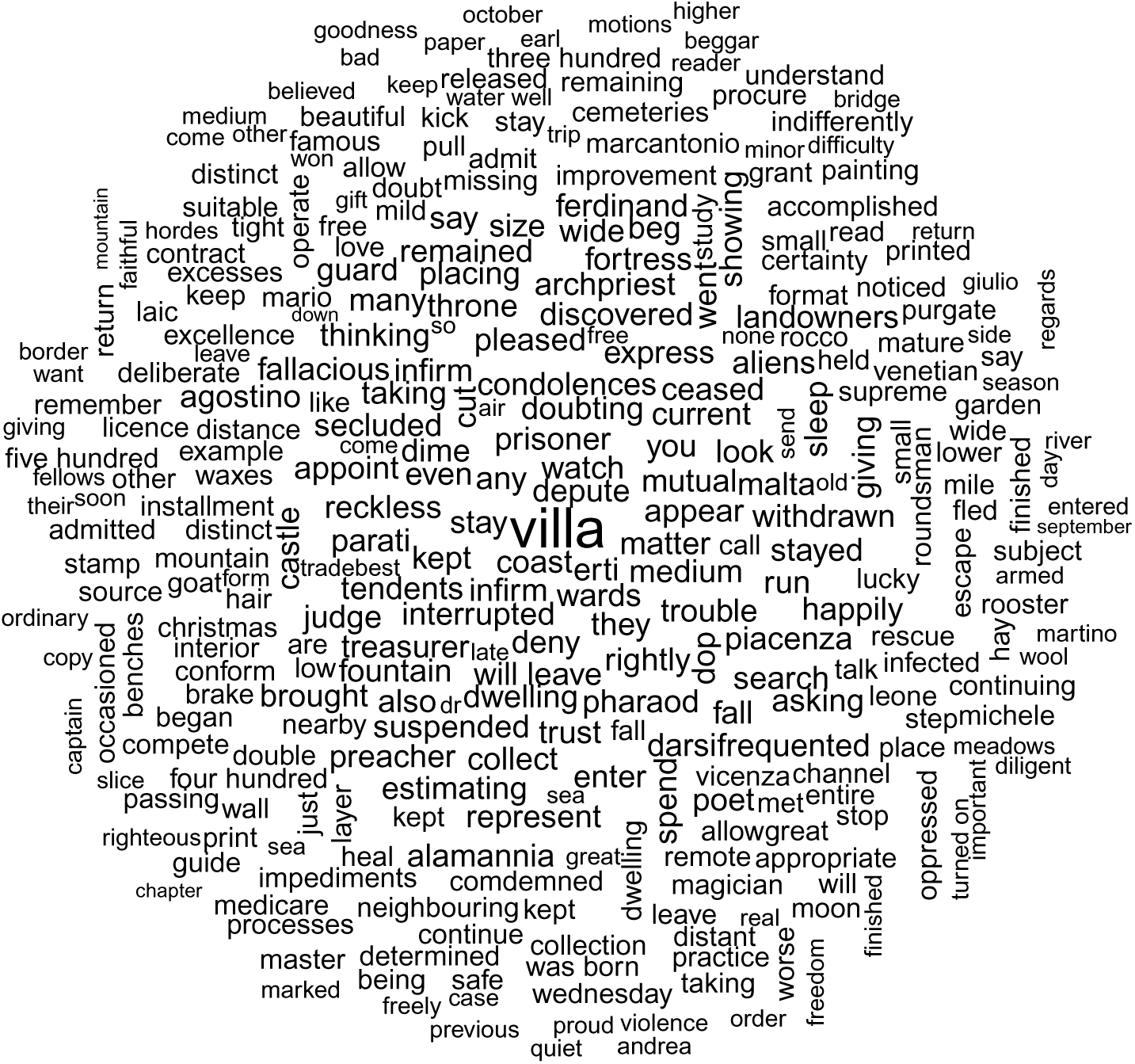
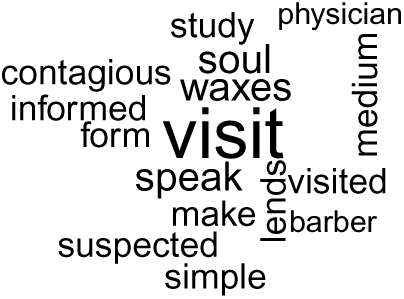
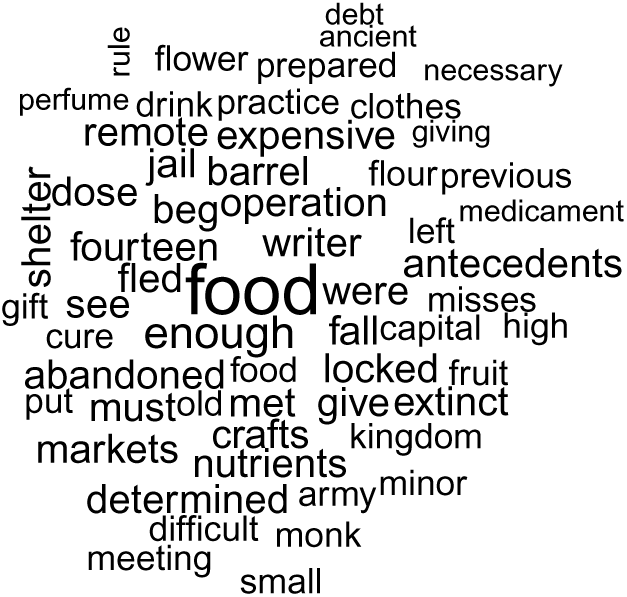
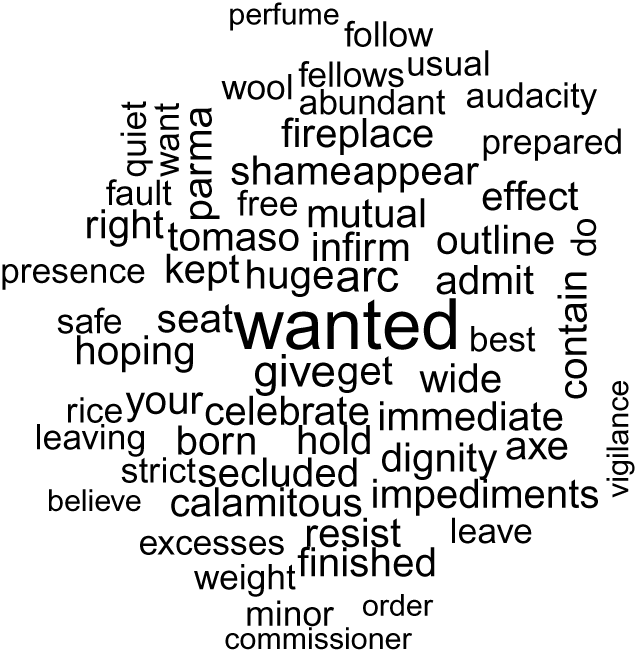

