## Supplementary Materials for "Unbiased lexicometry analyses illuminate plague dynamics during the second pandemic"

^1^ Aix Marseille Univ., IRD, MEPHI, IHU Méditerranée Infection, Marseille, 13005, France.

^2^ UMR 7268, Anthropologie bioculturelle, Droit, Ethique et Santé, Aix Marseille Univ, 11 CNRS, EFS, ADES, Marseille, 13344, France.

^3^ IHU Méditerranée Infection, Marseille, 13005, France.

^4^ Department of Biosciences and Pediatric Clinical Research Center “Romeo and Enrica Invernizzi”, University of Milan, Milan, 20133, Italy.

† These authors contributed equally to this work

IHU – Méditerranée Infection, 19-21 Boulevard Jean Moulin, 13005 Marseille.

Phone number: + 33 (0)4 13 73 20 01

Supplementary Materials and Methods

**Supplementary Text 1. Script.**

source("functions/functions.R")

require(data.table)

require(DESeq2)

require(stringi)

\# # Import Dictionaries

\# dic_FR <- fread("dictionnaries/dictionary_clean_FR.txt")

\# dic_IT <- fread("dictionnaries/dictionary_clean_IT.txt")

\#

\# # Import + clean texts

\# dat <- data.table(file= list.files("texts", full.names = T))

\# dat[, cdition:= ifelse(grepl("_FR_", file), "FR", "IT")]

\# dat[, sample:= gsub("_IT|_FR|.txt", "", basename(dat$file))]

\# dat <- dat[, .(raw_text= unlist(strsplit(readLines(file), " "))), c(colnames(dat))]

\# dat[cdition=="FR", filtered_text:= filtering(raw_text, dic_FR$words, plural_patterns = "FR")]

\# dat[cdition=="IT", filtered_text:= filtering(raw_text, dic_IT$words, plural_patterns = "IT")]

#------------------------------------------------------------#

\# 1- DESeq2 analyses

#------------------------------------------------------------#

\# dat[,

\#	{

\#	current <- na.omit(.SD)

\#	DF <- dcast(current, filtered_text~sample, fill = 0)

\#	DF <- data.frame(DF[, -1], row.names = DF$filtered_text)

\#	fwrite(current, paste0("results/word_counts_", cdition, ".txt"), col.names = T, row.names = F, sep= "\t", quote= F, na = NA)

\#	DF <- DF[rowSums(DF)>=5,]

\#	sampleTable <- data.frame(sample= unlist(tstrsplit(colnames(DF), "_", keep= 1)), row.names = colnames(DF))

\#

\#	# DESeq2

\#	dds <- DESeqDataSetFromMatrix(countData= DF, colData= sampleTable, design= ~sample)

\#	res <- DESeq(dds)

\#	diff <- as.data.table(as.data.frame(lfcShrink(res, contrast= c("sample", "plague", "control"))), keep.rownames= T)

\#	fwrite(diff, paste0("results/DESeq_FC_table_", cdition, ".txt"), col.names = T, row.names = F, sep= "\t", quote= F, na = NA)

\#	print("Global DESeq2 done")

\#

\#	# Nested

\#	sig <- diff[padj<=0.05 & log2FoldChange>0, .(tested_word= rn)]

\#	sig[, file:= tempfile(fileext = ".txt"), tested_word]

\#

\#	for(i in seq(sig$tested_word))

\#	{

\#	sub <- copy(current[!grepl("^control", sample)])

\#	.m <- which(sub$filtered_text==sig$tested_word[i])

\#	for(j in -25:25)

\#	{

\#	idx <- (.m+j)[between(.m+j, 0, nrow(sub))]

\#	sub[idx, check := T]

\#	}

\#	sub[is.na(check), check:= F]

\#	sub <- dcast(sub, filtered_text~check+sample, fill = 0)

\#	sub <- sub[filtered_text %in% diff$rn]

\#	DF_s <- data.frame(sub[, -1], row.names = sub$filtered_text)

\#	sampleTable_s <- data.frame(sample= unlist(tstrsplit(colnames(DF_s), "_", keep= 1)), row.names = colnames(DF_s))

\#

\#	# DESeq2

\#	dds <- DESeqDataSetFromMatrix(countData= DF_s, colData= sampleTable_s, design= ~sample)

\#	res_s <- try(DESeq(dds), silent = T)

\#

\#	if(class(res_s) == "DESeqDataSet"){

\#	diff_s <- as.data.table(as.data.frame(lfcShrink(res_s, contrast= c("sample", "TRUE", "FALSE"))), keep.rownames= T)

\#	diff_s <- diff_s[padj<=0.01 & log2FoldChange>0]

\#	}else{diff_s <- data.table()}

\#	saveRDS(diff_s, sig[i, file])

\#	print(paste0("Nested DESeq2", sig$tested_word[i], "done"))

\#	}

\#	sig <- sig[, readRDS(file), c(colnames(sig))]

\#	fwrite(sig, paste0("results/DESeq_FC_table_nested_", cdition, ".txt"), col.names = T, row.names = F, sep= "\t", quote= F, na = NA)

\#	}, cdition]

#--------------------------------------------------#

\# Save clean tables with traductions and so on...

#--------------------------------------------------#

t1 <- data.table(file= list.files("results", "DESeq_FC_table_nested", full.names = T))

t1 <- t1[, fread(file), file]

t1 <- t1[, c(1,2,4,5,6,7,8,9,10)]

t1[, type:= ifelse(grepl("_IT.txt", file), "nested_IT", "nested_FR")]

colnames(t1)[2] <- "nested_word"

t2 <- data.table(file= list.files("results", "DESeq_FC_table_IT|DESeq_FC_table_FR", full.names = T))

t2 <- t2[, fread(file), file]

t2$nested_word <- NA

t2[, type:= ifelse(grepl("_IT.txt", file), "global_IT", "global_FR")]

final <- rbind(t1, t2)

\# Translations

IT_EN <- fread("translations/translation_IT_EN_singular.txt")

IT_EN[, english:= tstrsplit(english, "/", keep= 1)]

final[IT_EN, translation:= ifelse(grepl("_IT", type), i.english, NA), on= "rn==italian"]

FR_EN <- fread("translations/translation_FR_EN_singular.txt", na.strings = "")

final[FR_EN, translation:= ifelse(grepl("_FR", type), i.english, translation), on= "rn==french"]

\# Signif cutoff

final[, significant:= ifelse(!is.na(padj) & padj<=0.05 & log2FoldChange>0, T, F)]

\# Correct words

final[rn=="meuble" & grepl("FR", type), translation:= "movable"]

final[rn %in% c("marchandise", "mercanzie", "merce"), translation:= "merchandise"]

\# Categories

cat <- unique(fread("categories/Mixed_word_categories.txt"))

cat <- cat[, .(corrected= tolower(unlist(tstrsplit(Word, "/")))), c(colnames(cat))]

cat_FR <- cat[Type %in% c("FR", "BOTH"), Category:= paste0(Category, collapse= ";"), corrected]

final[cat_FR, category:= ifelse(grepl("FR", type), i.Category, NA), on= "translation==corrected"]

cat_IT <- cat[Type %in% c("IT", "BOTH"), Category:= paste0(Category, collapse= ";"), corrected]

final[cat_IT, category:= ifelse(grepl("IT", type), i.Category, category), on= "translation==corrected"]

\# SAVE

final[, fwrite(.SD, paste0("results/final_table_", type, ".txt"), col.names = T, row.names = F, sep= "\t", quote= F, na= NA), type]

saveRDS(final, "results/all_data_final.rds")

**Supplementary Text 2. Information about the Italian word “roba”.**

A definition of the term in the XVII century is available from an Italian vocabulary published by the most important research institution of the Italian language (The Accademia della Crusca) at: https://books.google.fr/books?id=Nydc74IY6HUC&printsec=frontcover&dq=inauthor:%22Accademia+della+Crusca%22&hl=it&sa=X&ved=2ahUKEwiNl-Tngf3rAhUtz4UKHZBpCX0Q6AEwAHoECAIQAg#v=onepage&q&f=false

A detailed description of the origin and history of the Italian word “roba” (singular form of “robe”) is available at https://www.treccani.it/enciclopedia/roba-e-robone_%28Enciclopedia-Italiana%29: “All the clothes were formerly called "raubae" or "roboae" already in 1180 in the Roman d'Escloufe the set of all the garments that made up the complete male and female clothing is called robe.… In the seventeenth century, while robe and roboni have disappeared for over a century in the male costume, roba” takes up the meaning of a set of garments in women's clothing: the inventory of Isabella Infanta di Savoia (1610) lists roboni of very rich fabrics "with petticoats and jackets and detached sleeves ", and therefore" stuff (roba) "takes on the meaning of" stuff (roba) to put on ".

The term “robba” (singular form of “robbe”) is a dialectal form of the term “roba” (https://www.treccani.it/vocabolario/robba/).

**Supplementary Text 3. Plural rules for Italian word unification.**

The rules used are as follows:

1.	IF the word ends in "-co", change the last to letter to "-chi"

2.	IF the word ends in "-go", change the last to letter to "-ghi"

3.	IF the word ends in "-ca", change the last to letter to "-che"

4.	IF the word ends in "-ga", change the last to letter to "-ghe"

5.	IF the word ends in “-e”, change the last letter to “-i”

6.	IF the word ends in “-a”, change the last letter to “-e”

7.	IF the word ends in “-o”, change the last letter to “-i”

These rules work if the word is changed only if the “corresponding plural word” is in the word table (table containing all the words present at least 5 times in the texts used, control and plague-related). If the corresponding plural word is not present, the word remains unchanged. In this way, misconversions of words into an incorrect plural (nonexistent or not reflecting the correct meaning of the word) are greatly reduced. Clearly, errors are still generated.

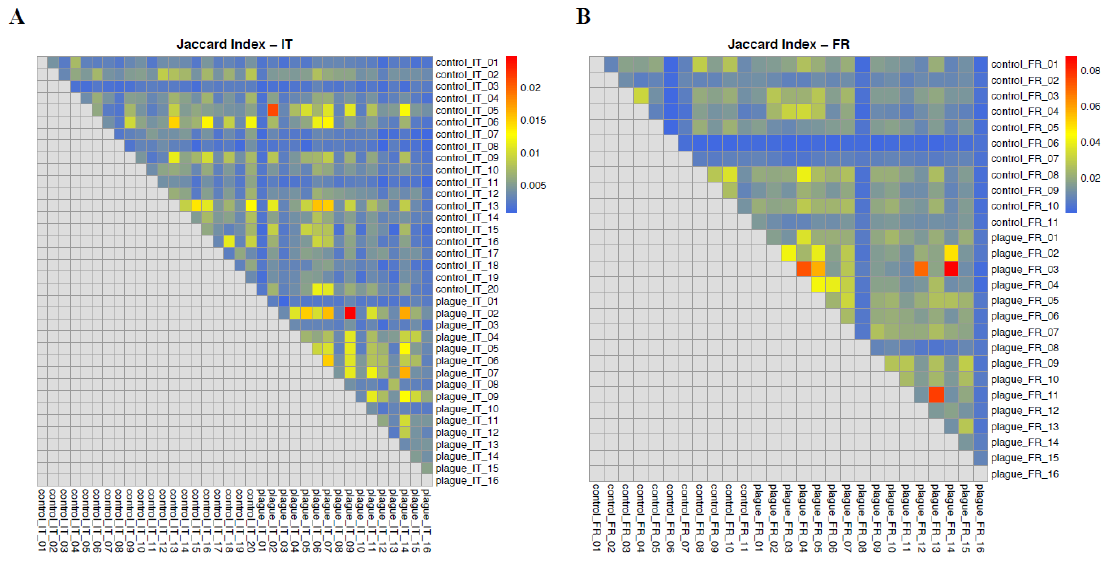

**Fig. S1.** **Jaccard index analyses for French texts corpus (A) (plague related-texts and control texts) and for Italian texts corpus (B).**

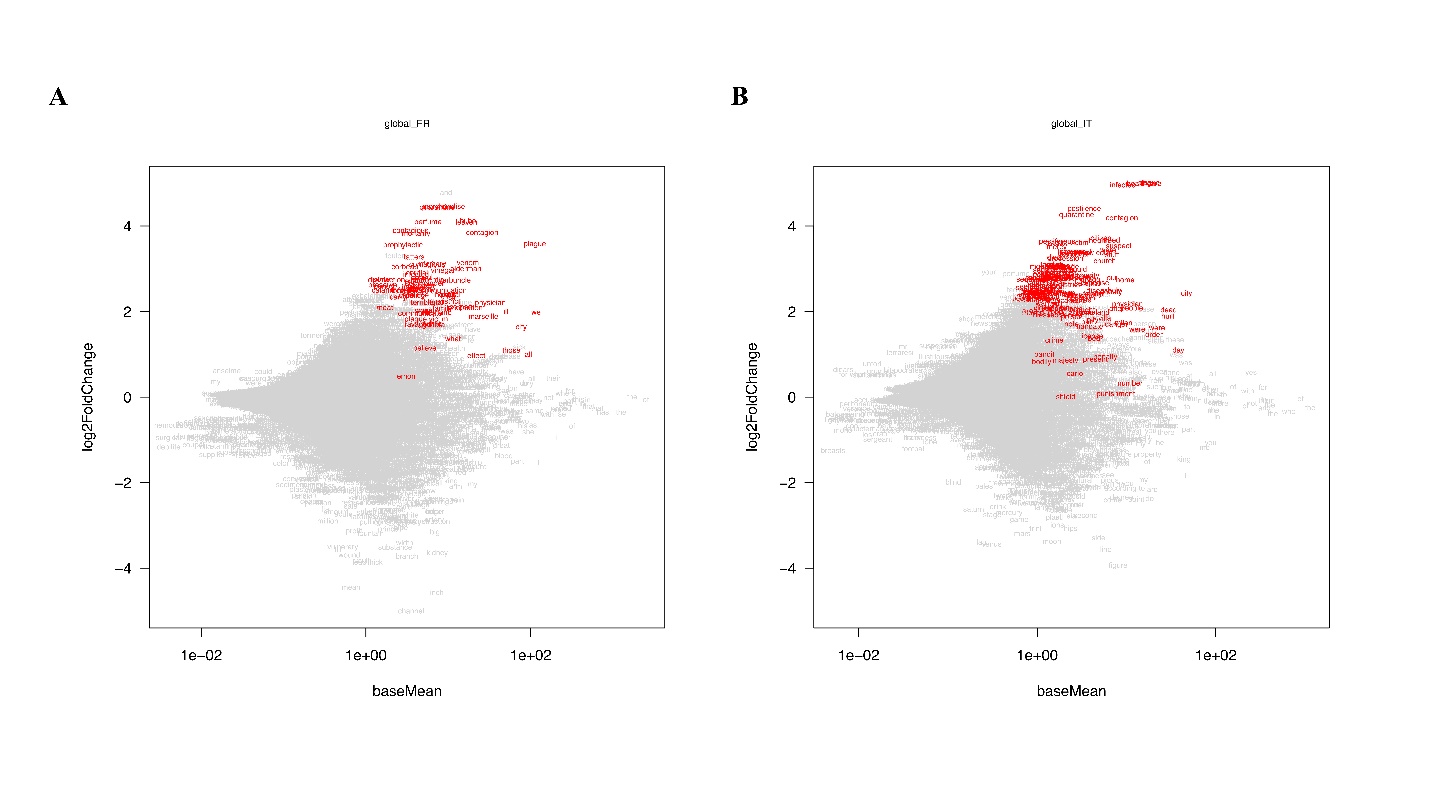

**Fig. S2. Volcano plot obtained from the analysis of the French texts corpus (A) and Italian texts corpus (B), generated using the DESeq2 package (log2foldchange >0 and P adjust<=0.05).**

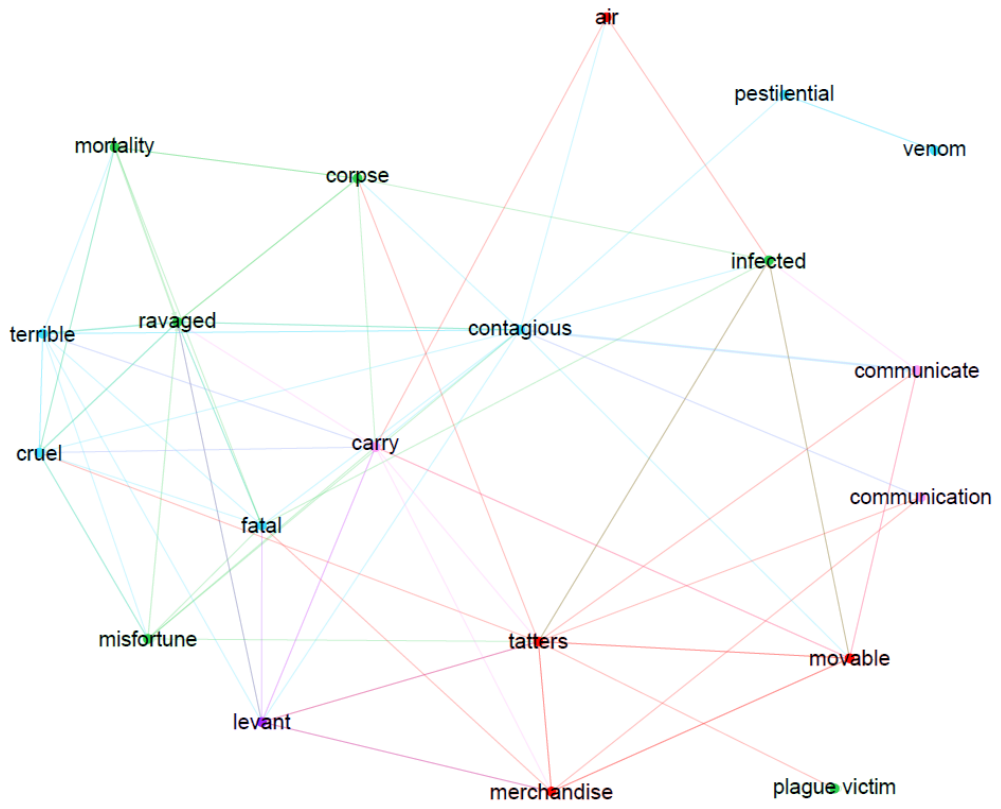

**Fig. S5. Word network representation for selected words from the Great Plague of Marseille (1720-1722).** Association between words that have been classified into five selected categories: plague sources (red), dissemination (pink), plague origin (violet), plague consequences (green), and plague nature (light blue). Network generated with Gephi software based on the adjusted P values from the nested analysis (P adj<0.01).

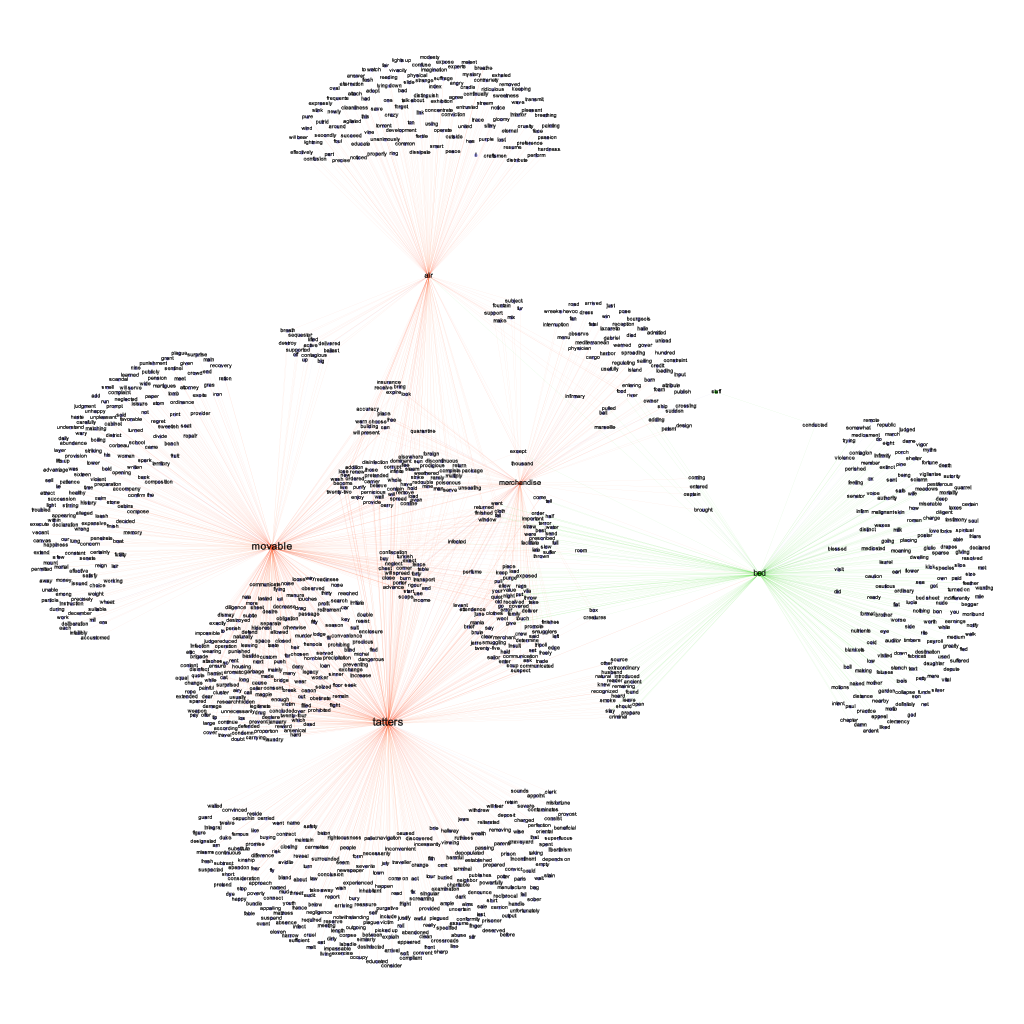

**Fig. S6**. **Word network representation of words associated to potential sources of plague.** Sources for the Marseille plague are displayed in green and for the Northern Italy plague are displayed in orange. Network generated with Gephi software based on the adjusted P values from the nested analysis (P adj<0.01).

Word label dimensions are proportional to the number of links with other words.

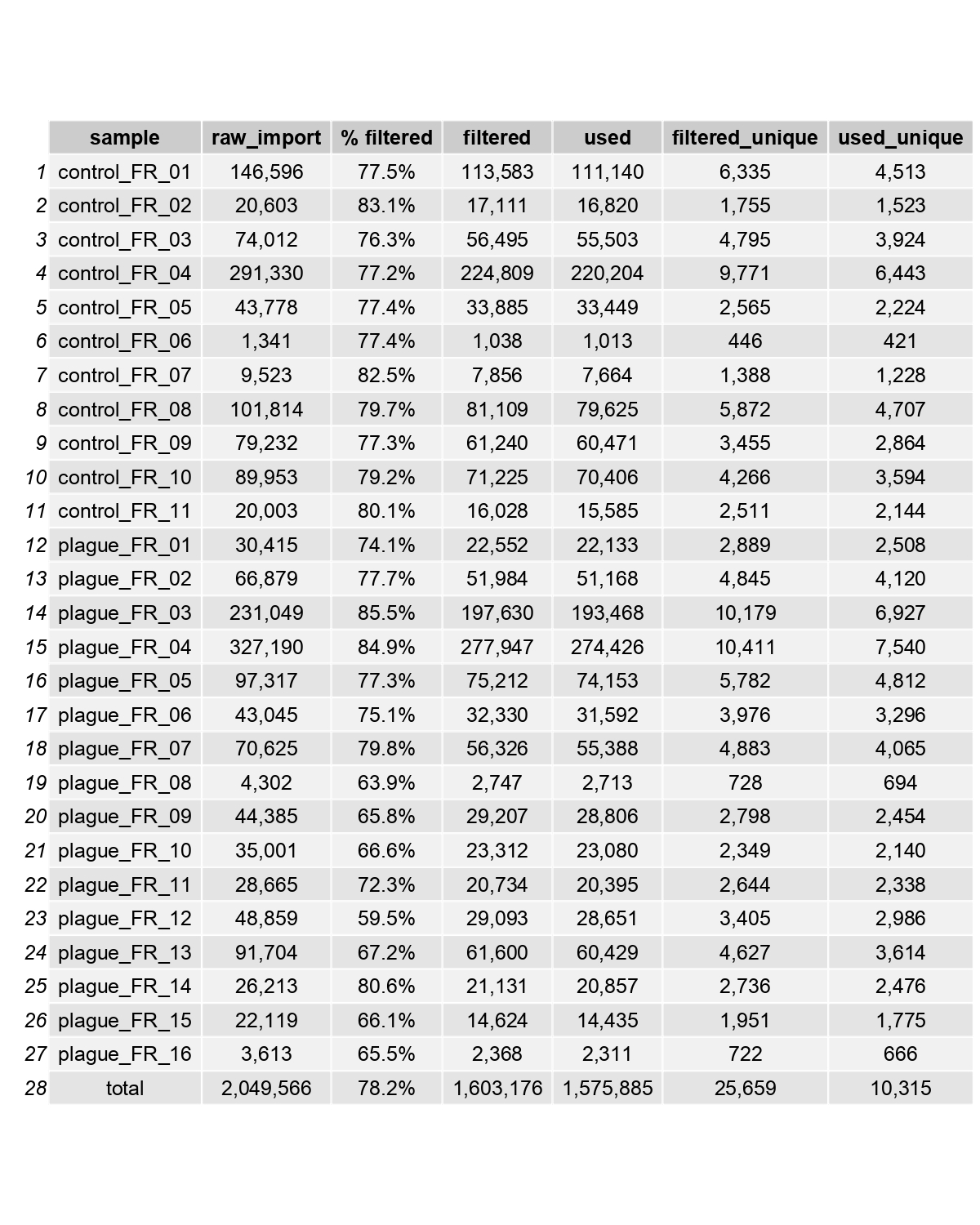

**Table S1. Statistical table summarizing the number of raw, filtered, used and unique words for each French plague-related texts and control texts.**

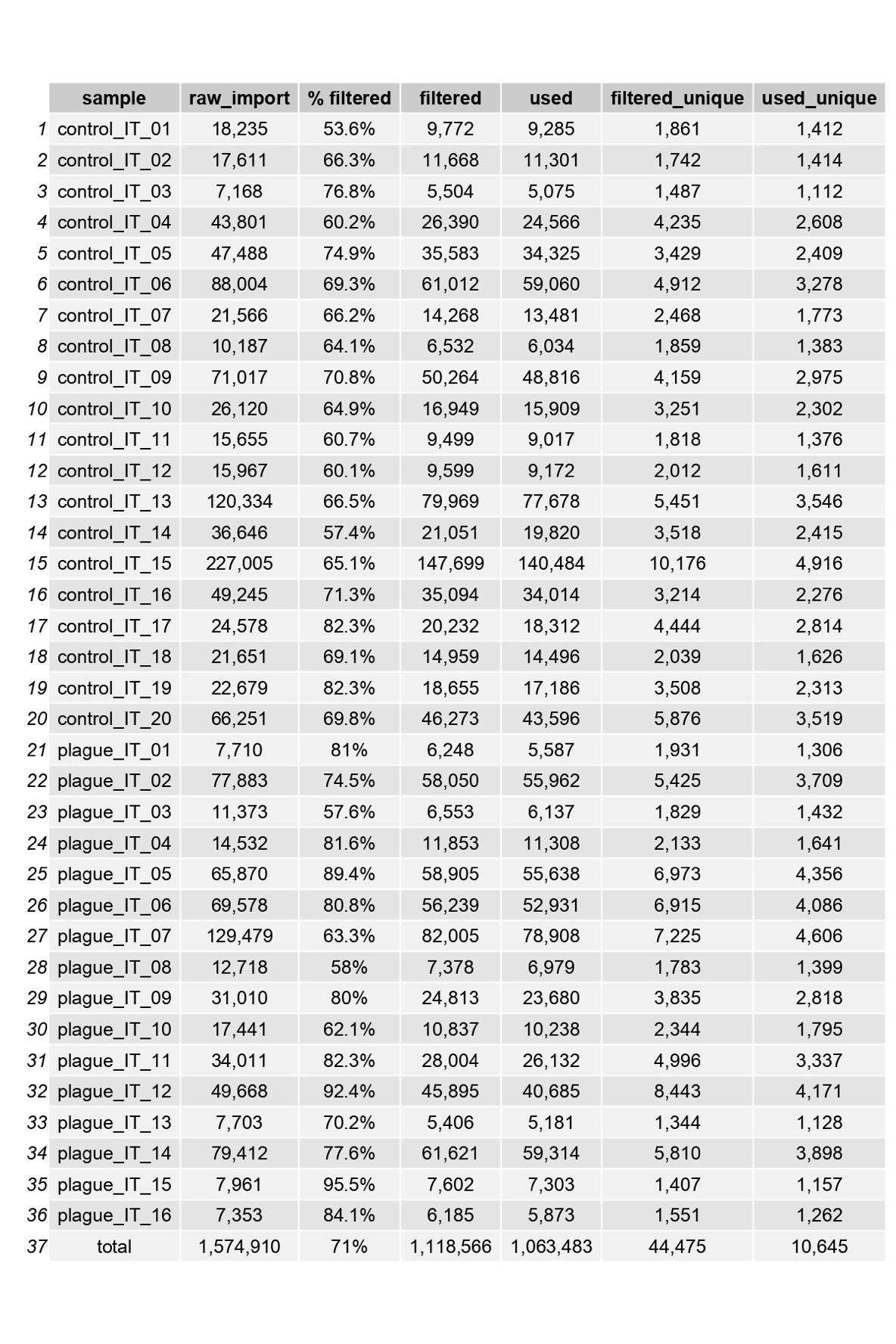

**Table S2.** **Statistical table summarizing the number of raw, filtered, used and unique words for each Italian plague-related texts and control texts.**

Other supplementary materials for this manuscript and available as separate files include the following:

Figures S3 and S4

Data file S1 to S9

**Fig. S3 (separate file).** **WordClouds generated from the nested analysis of French plague-related overrepresented words.** Each WordCloud is specific for one overrepresented word (present in the middle of the WordCloud) and displays every word that is associated with the relative overrepresented word (P value < 0.001 and log2foldchange>0).

**Fig. S4 (separate file).** **WordClouds generated from the nested analysis of Italian plague-related overrepresented words.** Each WordCloud is specific for one overrepresented word (present in the middle of the WordCloud) and displays every word that is associated with the relative overrepresented word (P value < 0.001 and log2foldchange>0).

**Data file S1 (separate file).** **Historical French text related to the 1720-1722 Great Plague of Marseille.**

**Data file S2 (separate file).** **Plague-unrelated French texts dating from the same historical period of the Great Plague of Marseille which were used as controls.**

**Data file S3 (separate file).** **All French words raw data generated by Deseq2 analysis.**

**Data file S4 (separate file).** **List of the overrepresented plague-related words with specific classification into 19 categories of words.**

**Data file S5 (separate file).** **All French words raw data generated by “nested” Deseq2 analysis.**

**Data file S6 (separate file).** **Historical Italian text related to the 1629-1631 plague epidemic in Northern Italy.**

plague_IT_03*. Originally written by the medic Alessio Alessandro during the epidemic and then printed again and amended in 1660;

plague_IT_05*. Originally written in 1631 and printed again in 1720 during the Plague of Marseille. The final part, not present in the original, reporting letters describing the Great Plague of Marseille have been removed before the analysis;

plague_IT_06*. Originally published in 1634 and printed again in 1714. The addition made in the edition of 1714 have been removed to only use the original material;

plague_IT_07*. Originally written during the epidemic in the city of Bergamo, but was published only in 1680 due mainly to political reasons (http://www.treccani.it/enciclopedia/lorenzo-ghirardelli_(Dizionario-Biografico)/);

plague_IT_12*. Text translation from latin by Francesco Cusani in 1841. The original book was written by Giuseppe Ripamonti in 1640. All the comments from the translator and the last book (5th book) were removed. The last book contains information about past epidemics not related to Milan and/or plague.

plague_IT_15*. Published in 1780 and it contains a report by Luca di Giovanni di Giovanni Targioni, granduncle of the author, from the years of the plague in Florence.

**Data file S7 (separate file).** **Plague-unrelated Italian texts dating from the same historical period of the plague epidemic in Northern Italy which were used as controls.**

**Data file S8 (separate file).** **All Italian words raw data generated by Deseq2 analysis.**

**Data file S9 (separate file).** **All Italian words raw data generated by “nested” Deseq2 analysis.**
